# Gene expression cartography of a developing neuronal structure

**DOI:** 10.1101/2025.03.30.646184

**Authors:** Leonardo Tadini, Lilia Younsi, Isabel Holguera, Félix Simon, Maximilien Courgeon, Nikos Konstantinides

## Abstract

The brain is a complex structure comprising thousands to billions of neurons that belong to thousands to millions of different neuronal types. These neurons often come from different progenitor domains and have very diverse developmental histories, yet they need to find their precise location in the brain and integrate into appropriate circuits. Therefore, describing the neuronal parts list of the brain in the form of single-cell mRNA sequencing atlases, while invaluable, lacks the spatial information that is instrumental to brain structures. While a large number of brain single-cell mRNA sequencing atlases have become available over the last years, spatial transcriptomic studies have lagged behind for a variety of reasons. Here, we use a gene expression cartography algorithm, called Novosparc, to reconstruct the spatial distribution of gene expression and cell type localization in a complex, yet tractable, developing brain structure, the *Drosophila* optic lobe. We generate a three-dimensional atlas of this structure, which we made available through a dedicated website (https://larva3dnovosparc.ijm.fr). This allowed us to identify spatially compartmentalized cell types and spatially patterned genes, to predict hotspots of neuronal cell death and identify the expression of neuronal type-defining transcription factors. Importantly, we identify caveats in such algorithms that could allow for the development of refined versions. Altogether, this work provides an invaluable tool for brain researchers to formulate hypotheses and paves the way for the generation of three-dimensional atlases of more complex brain structures, which will enhance our understanding of how neurons with diverse developmental lineages can integrate to form a functional brain.

## Introduction

The *Drosophila* visual part of the brain, i.e. the optic lobe, is a complex brain structure composed of ∼40,000 neurons (Apitz and Salecker, 2014; Nériec et al., 2016) that belong to, at least, 150 different neuronal types (Özel et al., 2021), as well of ∼5,000 glia of more than 15 different types (Lago-Baldaia et al., 2023). These neurons form synapses with each other in the four neuropils of the optic lobe, which are called lamina, medulla, lobula, and lobula plate (**Figure 1A**). The morphology and the transcriptome of the different cell types has been described extensively over the last thirty years(Fischbach and Dittrich, 1989; Konstantinides et al., 2018; Özel et al., 2021; Simon and Konstantinides, 2021), making the fly optic lobe one of the most comprehensively described adult neuronal structures.

**Figure 1:**
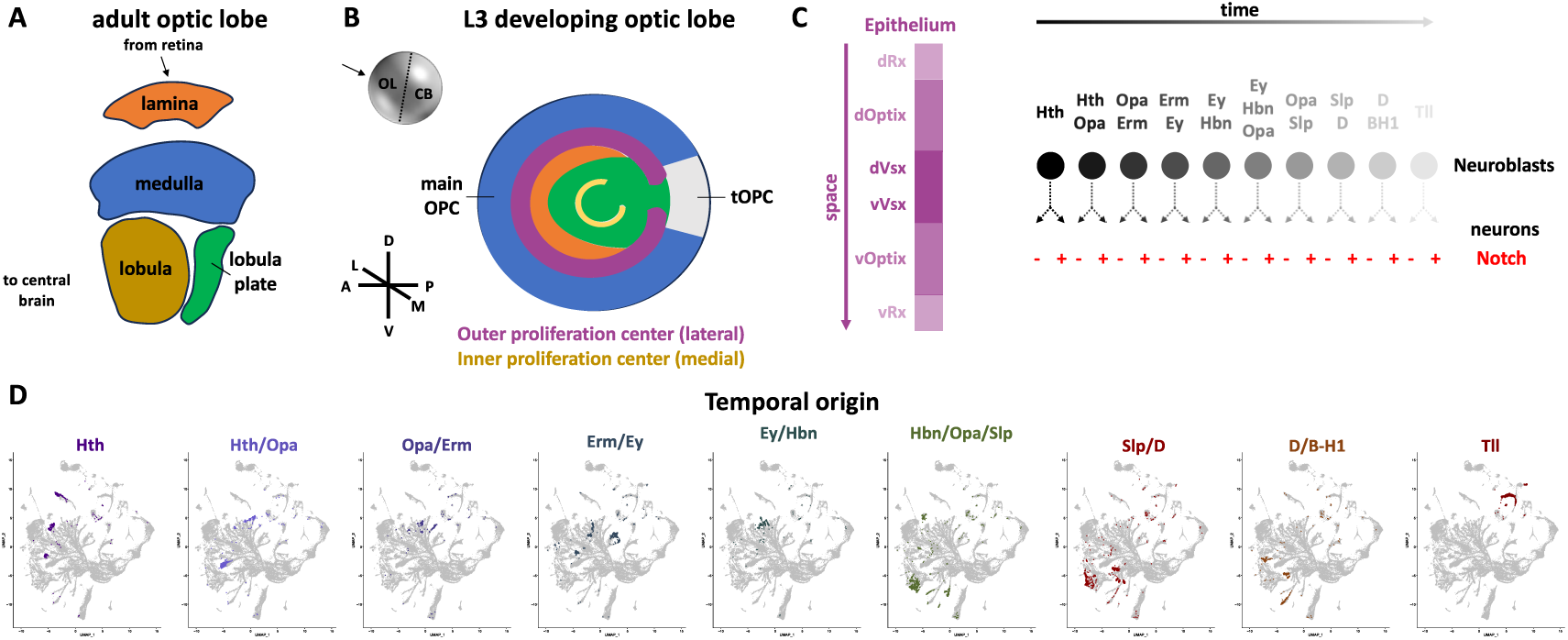
The developing *Drosophila* optic lobe. (A) Schematic of the adult optic lobe. It consists of four neuropils, the lamina, medulla, lobula, and lobula plate. (B) Schematic of a projection of the developing optic lobe in the late third instar larval stage, as seen from a lateral view. The medulla neurons (blue) come from the medial side of the main Outer Proliferation Center (magenta), while the lamina ones (orange) come from the OPC lateral sides. The tips of the OPC (tOPC) give rise to medulla and lobula neurons, while the Inner Proliferation Center (yellow) gives rise to lobula and lobula plate (green) neurons. (C) The main OPC gives rise to the majority of neuronal types of the optic lobe using an intersection of three mechanisms: first, the spatial patterning of the epithelium in six domains, second, the temporal patterning of neuroblasts in, at least, ten temporal windows, and, third, a Notch-driven binary cell fate decision upon the last division of the intermediate progenitors. (D) UMAP plots showing the localization of the different neuronal types that come from the respective temporal windows. In this two-dimensional representation, neurons with the same temporal origin tend to cluster together.

Like the more complex vertebrate brain structures, such as the cortex, the optic lobe neurons come from a variety of different developmental lineages. Most of the neurons are born from a neuroepithelial sheet called Outer Proliferation Center (OPC), which gives rise to the lamina cortex neurons in the lateral side and the medulla cortex neurons in the medial side (**Figure 1B**) (Nériec et al., 2016). The OPC is further divided in spatial domains, one of which (the tips of the OPC – tOPC) can give rise to some lobula cortex neurons as well (Bertet et al., 2014; El-Danaf et al., 2025; Erclik et al., 2017). The rest of the lobula cortex neurons come from another epithelial structure (the Inner Proliferation Center - IPC), which also gives rise to the most abundant neuronal cell type of the lobula plate, the T4/T5 neurons, as well as other C/T neurons **(Figure 1B)** (Apitz and Salecker, 2015). The rest of the lobula and lobula plate neurons come from the central brain neuroblasts (El-Danaf et al., 2025), which will not be discussed further in this study.

Apart from the comprehensive description of the neuronal cell type composition of the adult optic lobe, we also have a very good understanding of the developmental mechanisms that give rise to this diversity (Holguera and Desplan, 2018). A combination of spatial(Erclik et al., 2017) and temporal patterning(Konstantinides et al., 2022; Zhu et al., 2022) of the neuroepithelial cells and neuronal stem cells (neuroblasts), respectively, in combination with a Notch binary cell fate decision during the terminal division of an intermediate progenitor (the ganglion mother cell – GMC) is responsible for the generation of the different cell types of the medulla **(Figure 1C)**, which compose ∼70% of the optic lobe(Apitz and Salecker, 2014).

In an earlier study, we published a single-cell sequencing atlas of the third instar larval developing optic lobe (Konstantinides et al., 2022) **(Supplementary Figure S1)**. By sequencing ∼40,000 single cells (1X coverage of the brain) and using trajectory analysis algorithms, we were able to identify all the temporal transcription factors that are expressed in the medulla neuroblasts. While this was not part of the study, it did not escape our attention that the two-dimensional representation of the single-cell sequencing atlas (in the form of UMAP plot) retained a lot of spatial information, i.e. the lamina and the medulla were separated along the UMAP1 axis, while the lobula plate was distinguished along the UMAP2 axis **(Supplementary Figure S1)**. Moreover, neuronal types with similar temporal origins were clustered together **(Figure 1D)**, although this was not immediately obvious for the ones with similar spatial origins, as there are many cell types generated from more than one spatial domain (Malin et al., 2024; Simon et al., 2025) (**Supplementary Figure S1B**). While this information proved helpful, it lags considerably from a real three-dimensional spatial transcriptomic atlas.

To reconstruct the spatial localization of neuronal cell types and the spatial gene expression of the developing fly optic lobe, we decided to use Novosparc(Moriel et al., 2021; Nitzan et al., 2019), a gene cartography tool that has been used in the past to spatially reconstruct single-cell mRNA sequencing atlases of tissues, such as the stage 5 *Drosophila* embryo (Nitzan et al., 2019) and the mid-planula stage of *Nematostella (He et al., 2023)*. Novosparc relies on the hypothesis that physically neighbouring cells share a similar gene expression profile. Novosparc reconstructs spatial gene expression by leveraging a geometric framework that maps single-cell transcriptomic data onto spatial coordinates, preserving the tissue’s structural organization. It uses optimal transport-based algorithms to infer spatial positions of cells based on known spatial reference information.

Here, we used Novosparc to reconstruct in three dimensions gene expression emanating from the developing fly optic lobe single-cell mRNA sequencing atlas. To achieve this, we used pre-existing knowledge to divide the brain in different regions, based on their developmental lineage (Ngo et al., 2017), and generate a three-dimensional object representing the developing brain in Blender (https://www.blender.org). We used existing single-cell mRNA sequencing datasets (Konstantinides et al., 2022; Simon et al., 2025) to identify spatially and temporally informative genes that we used as landmarks while running the algorithm. We validated the performance of the algorithm by confirming the expression of the landmark genes, as well as predicting the expression of genes, for which the algorithm didn’t have any information. We validated their predicted expression using immunohistochemistry, existing Gal4 lines and Hybridization Chain Reaction Fluorescence *in situ* hybridization (HCR-FISH) (Tsuneoka and Funato, 2020). Finally, we used the reconstructed gene expression to predict hotspots of neuronal cell death, as well as temporally, spatially, and both temporally and spatially regulated neuronal type-defining transcription factors. Our work generates a powerful tool for predicting gene expression in a developing brain, sets a pathway for the use of such approaches for more complex brains and highlights the importance that developmental lineage plays in their generation.

## Results

To apply Novosparc, we required (a) a three-dimensional spatial framework onto which single cells could be mapped and (b) a set of marker genes to serve as spatial landmarks for this projection. To construct this framework, we first generated a regionalized three-dimensional spatial reference that enabled precise alignment of marker gene expression.

### Regionalization of the three-dimensional structure

The developing *Drosophila* optic lobe is a complex structure that is composed of different neuronal sets that are born from different neuronal linages. These different neuronal sets are adjacent to each other; however, their transcriptomes differ significantly reflecting their different developmental histories. Therefore, we first generated a three-dimensional structure corresponding to the developing optic lobe that reflected these different parts, using Blender. This was a half-sphere consisting of 41,287 vertices (corresponding to a similar number of cells in the L3 optic lobe and the respective single-cell mRNA sequencing dataset). This half-sphere was then separated in nine different regions after the (Ngo et al., 2017) 3D model that is based on confocal imaging z-projections of the fixed tissue (**Figure 2A**): a) the OPC epithelium, b) the OPC neuroblasts, c) the IPC epithelium, d) the IPC neuroblasts, e) the posterior medulla cortex neurons (T and C cells), f) the lamina neurons (L cells), g) the medulla neurons, h) the lobula plate neurons (T4/T5 cells), and i) the lobula neurons **(Figure 2B)**. In addition to these regions, we further added three more regions for j) surface glia, k) ganglion mother cells (**Figure 2C)**, and l) lamina wide-field neurons (Lawf1 and Lawf2), which were part of the sc-mRNA-seq datasets, but not of the 3D model.

**Figure 2:**
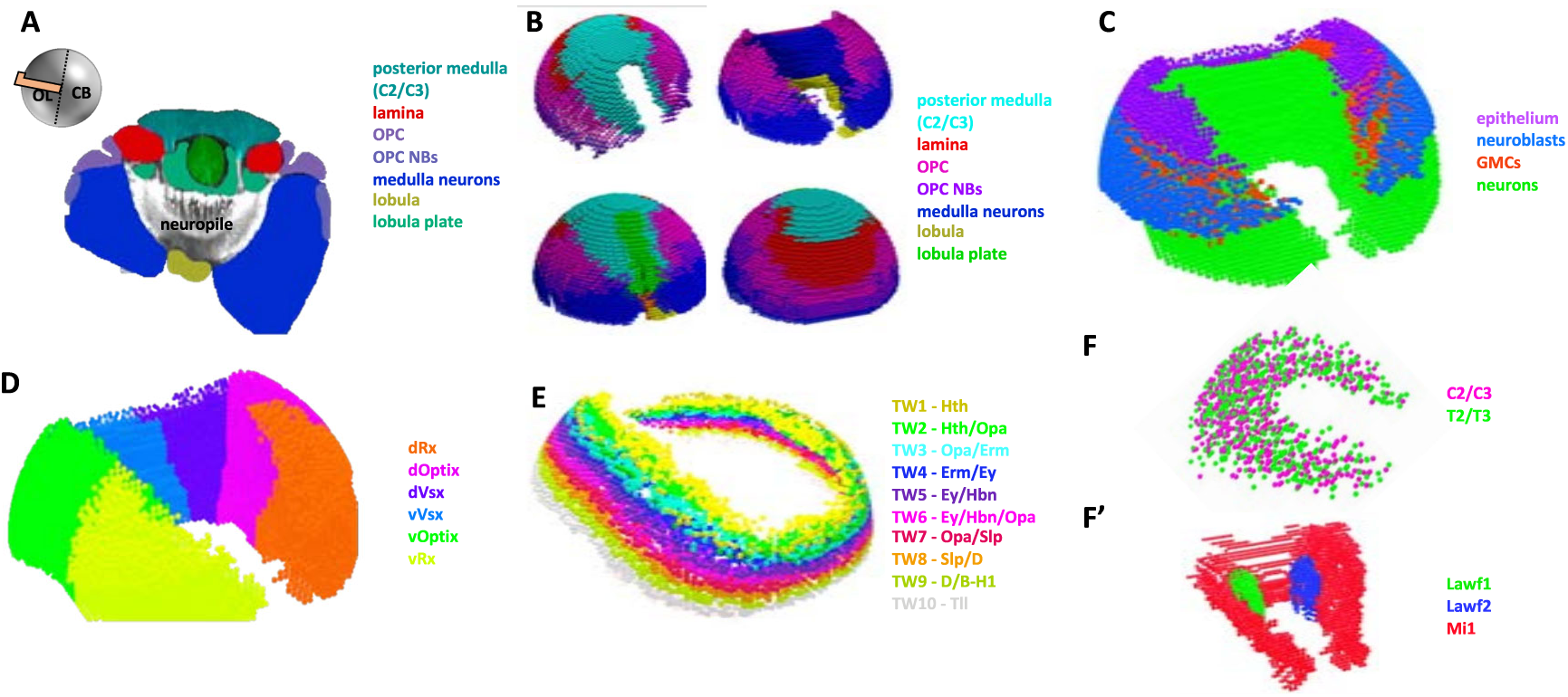
Regionalization of the three-dimensional model. (A) Color-coded cross-section of the developing optic lobe. Region annotation comes from (Ngo et al., 2017). (B) The three-dimensional model of the optic lobe was respectively divided into seven domains, i.e. the posterior medulla, lamina, OPC, OPC neuroblasts, medulla neurons, lobula, and lobula plate. (C) Visualization of the OPC epithelium (magenta), neuroblasts (blue), GMCs (red), and neurons (green). (D) The OPC epithelium, neuroblasts, GMCs, and neurons were further divided into six domains, based on the spatial patterning of the epithelium. (E) The medulla neuroblasts and GMCs were further subdivided into ten different temporal windows. (F-F’) The posterior medulla was subdivided into two populations mixed in a salt-and-pepper manner, C2/C3 and T2/T3 cells, while the Lawf1 and Lawf2 cells were added adjacent to the Mi1 neurons.

We then further subdivided some of these twelve primary regions into subregions, for which we could identify specific markers in the sc-mRNA-seq dataset. Therefore, we subdivided:

- the OPC epithelium into six spatial domains, corresponding to the dorsal and ventral Vsx (which account for 25% of the epithelium), Optix (50%), and Rx (25%) domains. The respective size of each spatial domain was decided based on the actual size of these domains (Malin et al., 2024) **(Figure 2D)**
- the OPC neuroblasts, GMCs, and neurons into the aforementioned six spatial domains and into 10 equally sized temporal domains, which correspond to the 10 temporal windows **(Figure 2E)**
- the posterior medulla cortex neurons into an equal number of intermingled C and T cells **(Figure 2F)**
- and the Lawf neurons into an equal number of Lawf1 and Lawf2 cells **(Figure 2F’)**

As a result, our final 3D model comprised 41,287 cells separated into 245 different regions (**Supplementary Table 1**). We then asked whether the cell number of each of these domains corresponds to the cell number from the sc-mRNA-seq dataset. For this purpose, we extended the annotation of the sc-mRNA-seq atlas, so that each of the cells corresponds to one of the above regions based on existing markers. The correspondence between the cell number of each of the regions in the Blender model and the estimated cell number from the sc-mRNA-seq dataset can be seen in **Supplementary Table 1**.

### Selection of appropriate marker genes and parameters

We then decided on spatially informative genes that would allow the algorithm to discriminate between the majority of the aforementioned 245 regions. For this purpose, we first removed from the dataset cells that correspond to the central brain (as this were not accounted for in the three-dimensional model in Blender) (**Figure S2A**) and then selected 43 different genes:

- we first selected genes that would differentiate neuroepithelium (shg, dpn), neuroblasts (shg, dpn, ase), GMCs (ase, elav), neurons (elav), and glia (repo, wrapper) (**Figure 3A** and **Supplementary Figure S2B**).
- we also selected the temporal (hth, Dll, erm, opa, ey, hbn, scro, slp1, D, B-H1, tll) and spatial (Vsx1, Optix, Rx) transcription factors (**Figure 3B** and **Supplementary Figure S2C**).
- we selected genes that would discriminate the different neuropils (acj6, eya, Lim1) (**Figure 3C**).
- finally, we selected different genes that would differentiate the different neuronal types. We focused on genes that are expressed differentially in neurons based on their spatial, temporal, and Notch origins (Simon et al., 2025) (aop, ap, bsh, dac, CG34340, Ets65A, fd59A, fkh, Hmx, kn, Lim3, oc, run, sim, Sox102F, svp, tj, toy, tup, vvl) (**Figure 3D** and **Supplementary Figure S2D**).

**Figure 3:**
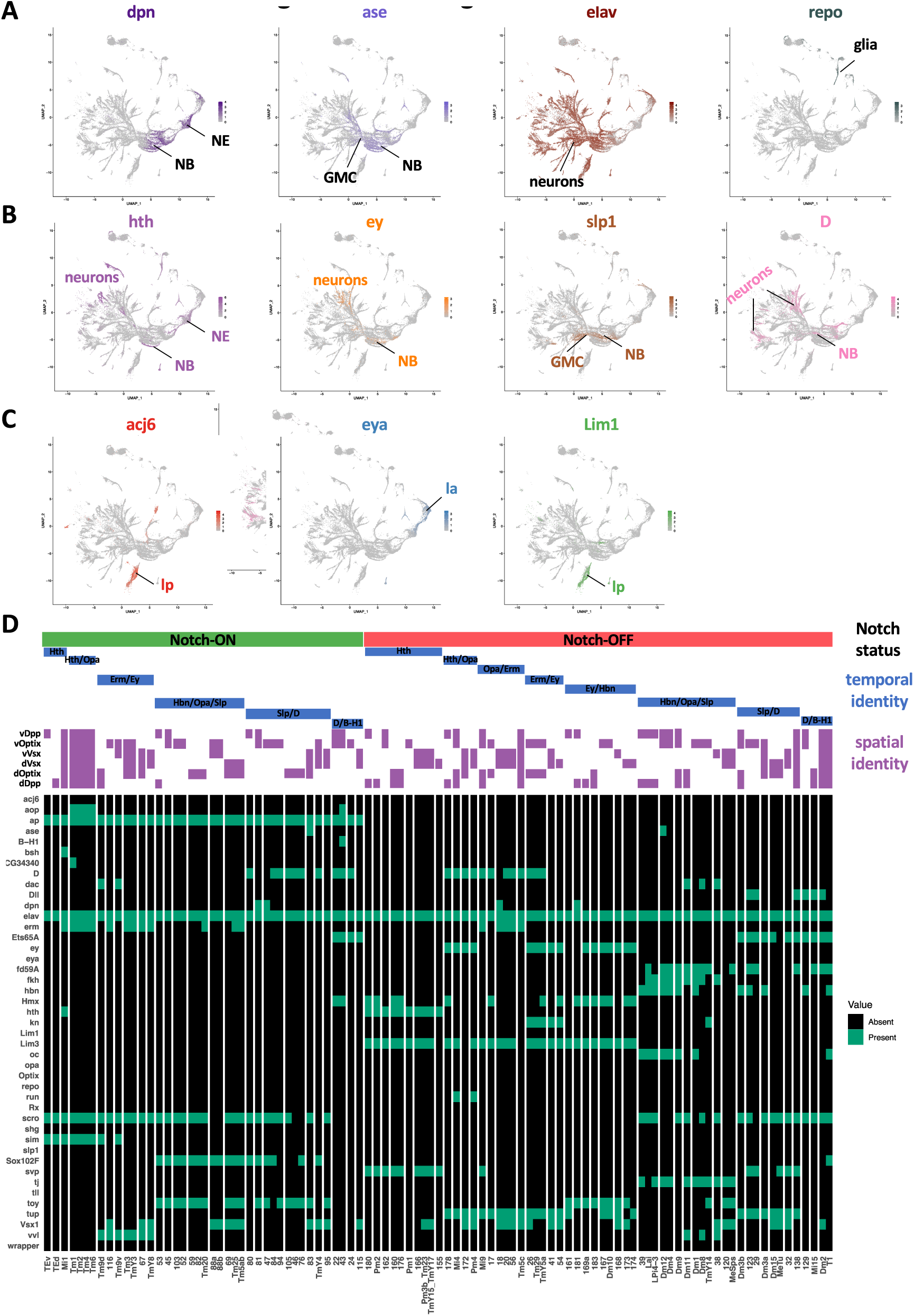
Marker gene selection. (A) UMAP plots showing the expression of gross cell types markers. Combinations of gross cell type markers, such as *dpn*, *ase*, *elav*, and *repo* can differentiate between neuroepithelium, neuroblasts, GMCs, neurons, and glia. (B) UMAP plots showing the expression of temporal transcription factors. Combinations of temporal transcription factors can differentiate between the different neuroblast and GMC temporal windows, as well as many neuronal types. (C) UMAP plots showing the expression of different neuropil neuron markers. Combinations of these markers can differentiate the lobula plate and lamina from the medulla neurons. (D) On/off expression heatmap of the different landmark genes used in neuronal types of different spatial, temporal, and Notch origin. Data come from the mixture modeling (Davis et al., 2020) used in (Simon et al., 2025).

All the marker genes and their corresponding landmark expression can be found in **Supplementary Table 2.**

We also tested extensively the different parameters that influence the performance of Novosparc (Moriel et al., 2021), i.e.

a) the alpha parameter, which controls the interpolation between two modes of reconstruction: de novo spatial reconstruction (**α = 0**) and reconstruction based solely on the information provided by the reference atlas (**α = 1**),
b) the number of genes used for the reconstruction: This defines the number of genes used for the reconstruction. It is generally advisable to include only the most differentially expressed genes, as these are more likely to be spatially informative (i.e., expressed in some cells but not others, ideally across a well-defined spatial domain). Based on the above criterion, *elav* was not used in the reconstruction as it was not among the most differentially expressed genes.
c) the number of nearest neighbors, which determines the number of neighbors used to construct the k-nearest neighbors (kNN) graphs for cells and spatial locations, and
d) the epsilon parameter, which is associated with the entropic regularization term. A low epsilon value leads to more localized mapping, while a higher epsilon value results in higher-entropy mappings, approaching a uniform distribution.

Details about parameter selection can be found in the Materials and Methods and in **Supplementary Figure S3.**

### Evaluation of Novosparc run

Using the above spatial reference, parameters, and markers, we reconstructed in three dimensions the sc-mRNA-seq dataset of the third instar larval developing optic lobe using Novosparc.

The first test for the performance of Novosparc was to evaluate the expression of the genes that were used as landmarks. Since the alpha parameter was set to 0.5, Novosparc was free to navigate between a *de novo* and a guided reconstruction. The reconstructed expression of the landmark genes can be visualized in **Figure 4** and **Supplementary Figure S4**. We compared the reconstructed expression with the actual expression of some of these marker genes using antibody staining (**Figure 4**).

**Figure 4:**
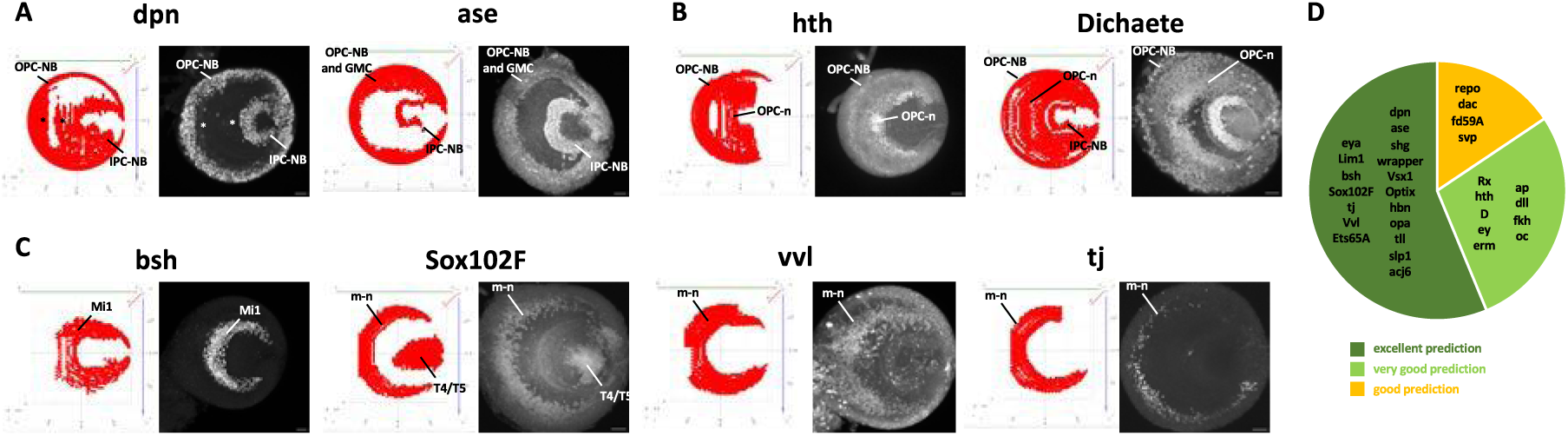
Reconstructed marker gene expression. Comparison of reconstructed marker gene expression as seen in https://larva3dnovosparc.ijm.fr and as seen by antibody staining against the respective proteins. (A) *dpn* and *ase* are expressed as expected, the neuroblasts (medulla and lobula plate neuroblasts, m-NB and lp-NB) and neuroblasts and medulla GMCs (m-GMCs), respectively. Notably, *dpn* transcript can be obviously detected in the OPC and IPC neuroepithelia (asterisks), in which protein expression is very faint. (B) *hth* can be detected in young medulla neuroblasts (m-NB), as well as medulla neurons (m-n), in both reconstructed gene expression and antibody staining. Similarly, *D* can be detected in medulla and lobula plate neuroblasts, as well as many medulla neurons in both representations. (C) Neuronal markers, such as *bsh,* Sox102F, vvl, and tj are detected in the same expression pattern in both reconstructed gene expression and antibody stainings. (D) Pie chart showing the prediction quality of the 31 tested reconstructed marker genes. 18/31 (55%) were reconstructed almost perfectly, 9/31 (29%) were reconstructed very well, while only 4/31 (16%) were reconstructed well.

Among the genes that differentiate cell classes (neuroepithelium, neuroblasts, GMCs, neurons, and glia), *dpn* is expressed in the OPC and IPC neuroepithelium (marked with asterisk) and the neuroblasts of the medulla and the lobula plate (**Figure 4A**), as it is obvious also from the sc-mRNA-Seq (**Figure 3A**). Notably, Dpn protein is only expressed very faintly in the neuroepithelium, as shown by the respective antibody staining and it is predominantly evident in the neuroblasts. *ase* mRNA and Ase protein are found in neuroblasts and GMCs (**Figure 4A**) that come from the OPC and IPC, while *shg* mRNA can be found in the neuroepithelium and the neuroblasts, as expected (**Supplementary Figure S4A and Supplementary Figure S2B)**. On the other hand, the expression of *repo* can only be found in the glia that are born during the last division of the main OPC neuroblasts (**Supplementary Figure S4A**), which shows that migrating cells like glia are very hard to reconstruct with such approaches (discussed further in the “Limitations of this study” Discussion section). Wrapping glia do not express *repo*, but express *wrapper* mRNA as expected (**Supplementary Figure S4A and Supplementary Figure S2B).**

Among the spatial transcription factors, Vsx1 is expressed in the central part of the OPC neuroepithelium, as well as in medulla neurons that predominantly, but not exclusively, come from this neuroepithelial region (asterisk), and Optix and Rx are expressed in the respective regions of the neuroepithelium (**Supplementary Figure S4B)**. We also tested for the expression of the temporal genes *hth, D, erm, ey, slp1, hbn, opa,* and *Tll*. *Hth* is expressed in the medulla NBs, as well as some medulla neurons, as also shown by antibody staining (**Figure 4B**). Interestingly, *hth* is also expressed in glia, as will be further discussed later. Similarly, *D* is expressed in medulla neuroblasts and neurons, as well as the IPC neuroblasts, as was already known (Apitz and Salecker, 2015) (**Figure 4B**). *erm* is expressed very broadly as can also be confirmed by antibody staining, *ey* is expressed in medulla neuroblasts and many different medulla neurons, *slp1* is expressed exclusively at a single temporal window of the medulla neuroblasts, *hbn* is expressed in medulla neuroblasts and neurons, C/T cells, and Lawf neurons, *opa* is expressed in medulla neuroblasts and GMCs in two temporal windows, and *tll* is expressed in old medulla neuroblasts, lobula plate neuroblasts, and the lamina (**Supplementary Figure S4C)**.

We also looked for the spatial reconstruction of genes that differentiate different neuropil origin. As expected, *acj6* is expressed exclusively in the lobula plate T cells, *eya* is expressed in the lamina and lamina wide-field cells (Lawf1 and Lawf2), while *Lim1* in T4/T5 cells and Lawf2 (**Supplementary Figure S4D)**.

Finally, we probed for the expression of the neuronal type-specific genes*, bsh, Sox102F, vvl, tj, ap, dac, dll, ets65A, fd59A, fkh, kn, oc, svp,* and *tup*. As can be seen in **Figure 4C** and **Supplementary Figure S4E**, their reconstructed expression matches to a large extent their actual expression as revealed by antibody stainings. Overall, we evaluated the quality of the reconstruction of 31 marker genes, 27 of which were reconstructed accurately (**Figure 4D**).

### Prediction of expression of non-reference genes

We then tested the reconstruction with a more difficult task, which was to predict the expression of genes that were not used as a reference. For this purpose, we decided to probe different types of genes, such as transcription factors, neuronal effector genes (in particular, different cell surface molecules), as well as signaling molecules (in particular, members of the Wnt family, which is well known to regulate neuronal diversification in the visual system (Bertet et al., 2014; Özel et al., 2021; Suzuki et al., 2016)) (**Figure 5**).

**Figure 5:**
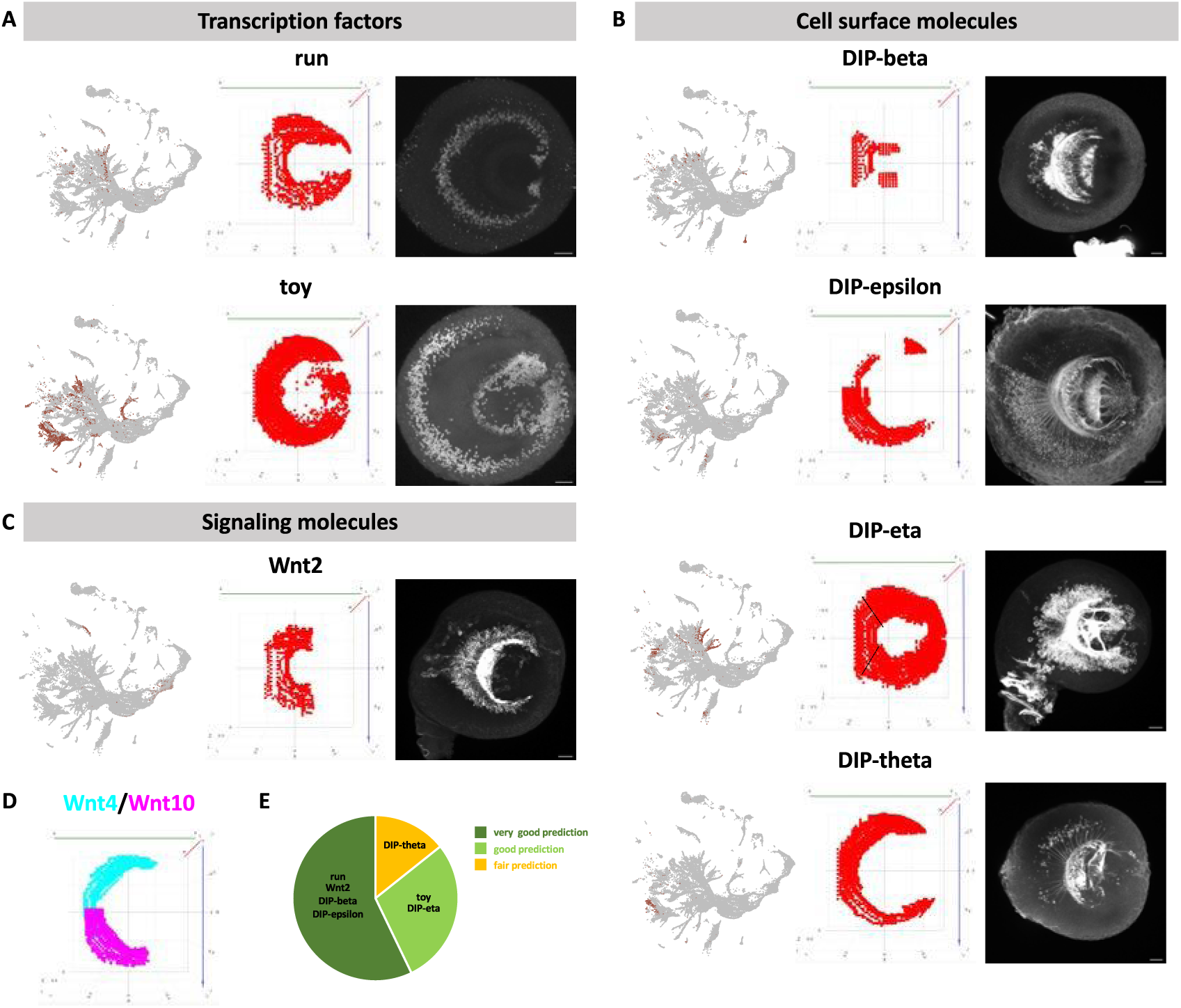
Non-reference gene expression. (A) Transcription factor expression in the larval developing optic lobe, as can be seen in a UMAP plot, the Novosparc reconstruction, and by antibody staining. The reconstructed expression of both *run* and *toy* matches their actual expression pattern. (B) Cell surface molecule expression in the larval developing optic lobe, as can be seen in a UMAP plot, the Novosparc reconstruction, and by antibody staining of the respective T2A-Gal4 line driving GFP. The reconstructed expression of *DIP-beta, DIP-epsilon*, and *DIP-eta* matches their actual expression pattern. The expression of *DIP-theta* is similar, but the Novosparc reconstruction predicted broader expression. (C) Cell surface molecule expression in the larval developing optic lobe, as can be seen in a UMAP plot, the Novosparc reconstruction, and by antibody staining of the Wnt2-Gal4 line driving GFP. The reconstructed expression of *Wnt2* matches its actual expression pattern. (D) The reconstructed expression of Wnt4 (cyan) and Wnt10 (magenta) matches their published expression pattern, in which Wnt4 is expressed ventrally and Wnt10 is expressed dorsally. (E) Pie chart showing the prediction quality of the 7 tested reconstructed non-reference genes. 4/7 (57%) were reconstructed very well, 2/7 (29%) were reconstructed well, while only 1/7 (14%) was reconstructed fairly.

*run* is expressed in progeny of neuroblasts of the Hth/Opa temporal window, such as Mi4 and Pm4 (Konstantinides et al., 2022), as also shown in the UMAP plot of the sc-mRNA-seq. Indeed, according to both antibody stainings and Novosparc reconstructed 3D expression, *run* is expressed in a thin concentric ring in the medulla cortex, indicative of its expression in the neuronal progeny of a single temporal window (**Figure 5A**). On the other hand, toy is expressed much more broadly, in particular in neurons that are born during the Ey/Hbn, Ey/Hbn/Opa, and Scro/Opa/Slp temporal windows (Konstantinides et al., 2022), as indicated also by the broad expression in the UMAP plot of the sc-mRNA-seq. Indeed, the toy mRNA reconstructed expression and the Toy antibody staining reveals a broad ring of expression. Importantly, Toy is also expressed in C2/C3 neurons which can be seen in the center of the developing lobe (**Figure 5A**).

We then looked for the cell type-specific expression of cell surface molecules. As these proteins are predominantly localized at synaptic sites, which would not be informative for the cell expression to compare to the Novosparc reconstruction, we tagged *DIP-beta, DIP-epsilon, DIP-eta*, and *DIP-theta* with a T2A-Gal4 using the “plug-and-play” system (Diao et al., 2015) to drive expression of GFP in the cell types, where these genes are expressed. *DIP-beta* is expressed in few cell types, as can be seen by the UMAP and the antibody staining, including Lawf1 and Lawf2 and other cells that likely originate from the Vsx1 domain (**Figure 5B**). The 3D reconstruction of Novosparc predicts perfectly this expression, but it misses *DIP-beta*-expressing cells that are found in the Optix domain. Similarly, *DIP-epsilon* is also expressed in few cell types with an obvious enrichment in cells in the ventral side of the optic lobe. The 3D reconstruction identifies accurately this difference in the dorsoventral localization of *DIP-epsilon-*expressing cells (**Figure 5B**). *DIP-eta* is expressed across the entire medulla neuronal cortex with a slight but noticeable depletion in the neurons that come from the Vsx1 domain, which is obvious in both antibody staining and 3D reconstruction (**Figure 5B**). Finally, *DIP-theta* is also expressed in different neuronal types that are primarily localized in the ventral side of the optic lobe. Novosparc reconstruction identifies this difference along the dorsoventral axis, however it seems less pronounced, as it identifies more cells in the dorsal side than the ones that are marked with the Gal4 line (**Figure 5B**).

Finally, we looked for the localized expression of different Wnt proteins. *Wnt2* is expressed in immature Mi1 cells (as indicated by the UMAP) and can be seen in few neurons that span the medulla cortex, as indicated by both 3D reconstruction and Gal4 expression (**Figure 5C**). Finally, we have previously published that *Wnt4-* and *Wnt10-*expressing cells are localized ventrally and dorsally, respectively (Özel et al., 2021). The Novosparc reconstruction can accurately recapitulate this expression pattern (**Figure 5D**). In total, out of the 7 genes that we tested, 4 were predicted very well, two were predicted well, while only *DIP-theta* expression was less accurate (**Figure 5E**). Notably, the percentage of prediction accuracy of non-reference matches to that of the marker genes (**Figure 4D**), indicating that this percentage of accuracy (60% very good, 30% good, 10% fair) is expected for the rest of the genes.

### Novosparc predicts localized programmed cell death driven by *reaper* and *grim*, as well as *hth*- expressing glia

Finally, we wanted to take advantage of this newly constructed three-dimensional atlas to make predictions about the expression of interesting genes. Programmed cell death is an essential and conserved mechanism that regulates neuronal diversity and the ultimate neuronal number in a developing nervous system (Pinto-Teixeira et al., 2016). Notably, cell death is regulated by both internal factors (such as temporal patterning) (Bertet et al., 2014) and external mechanisms (such as metabolism and hormones) (Draizen et al., 1999). Four pro-apoptotic genes (Cashio et al., 2005) regulate apoptosis in the developing *Drosophila* optic lobe, *hid*, *reaper*, *grim*, and *skl*. Their expression in the UMAP plots show that *hid* is expressed in the tips of neuronal trajectories of neurons that do not undergo apoptosis, which could be indicative of stress or of a non-apoptotic role for *hid*, while *reaper, grim*, and *skl* are expressed in specific cell types that are fated to die and are concentrated in a specific part of the plot (**Figure 6A**). We wanted to see how these cells are distributed in the 3D reconstruction. As expected, *hid-*expressing cells can be found spread in the entire medulla cortex. On the other hand, the expression of *reaper, grim,* and *skl* appears to be partially overlapping and spatiotemporally regulated (**Figure 6A**), which suggests that specification mechanisms regulate the neuronal cell types that will undergo programmed cell death in the entire medulla cortex, as has been published for the tOPC neuronal types before (Bertet et al., 2014).

**Figure 6:**
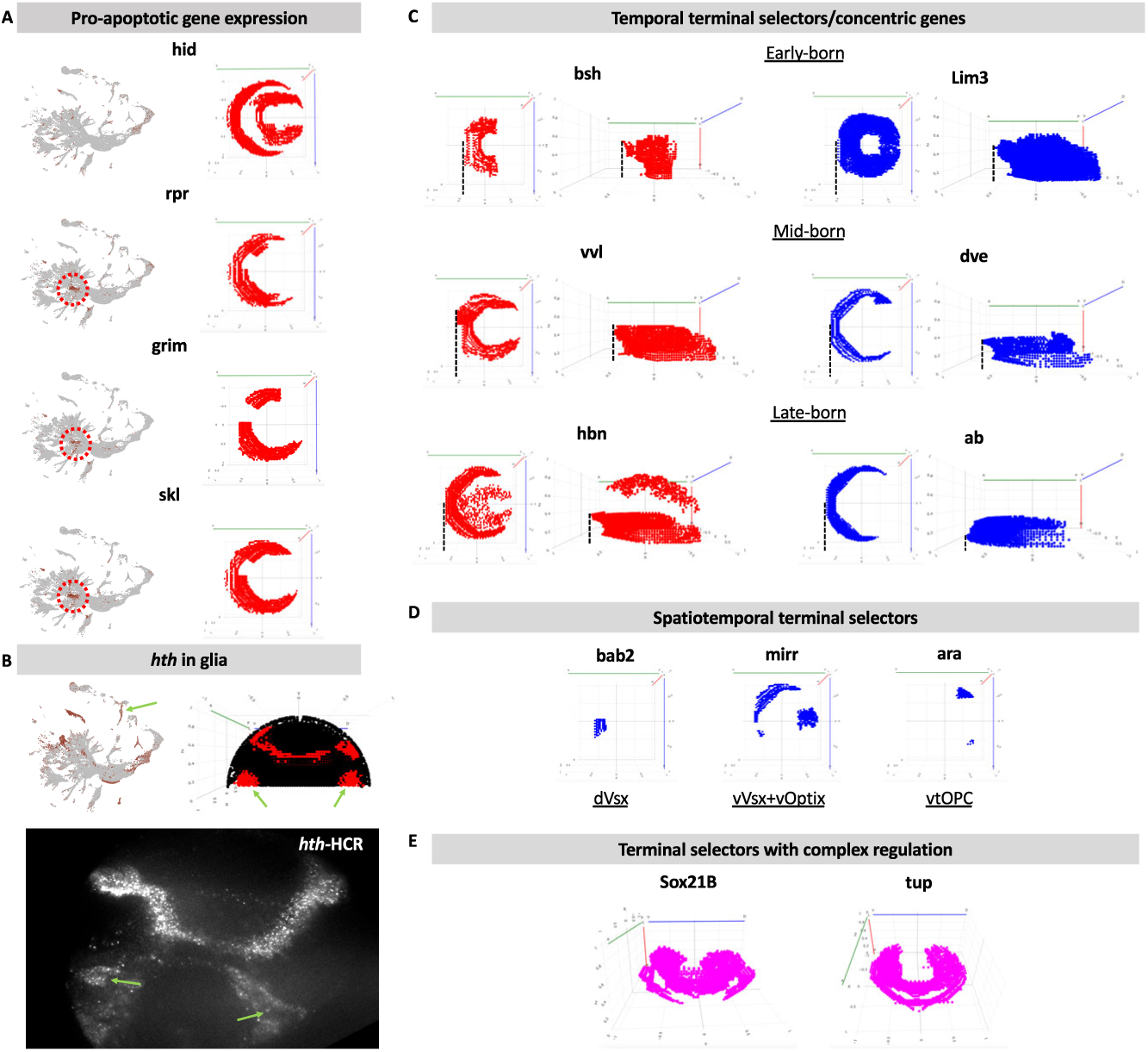
Spatio-temporal regulation of pro-apoptotic genes and terminal selectors. (A) Pro-apoptotic gene expression in the larval developing optic lobe, as can be seen in a UMAP plot and in the Novosparc reconstruction. *hid* is expressed in all differentiating neuronal cells types, as can be seen by its presence in the tips of all trajectories in the UMAP plot and the broad expression in the medulla cortex in the Novosparc reconstruction. *rpr, grim*, and *skl* expression is localized in a cluster of cells in the UMAP plot, as well as in distinct locations along the medulla cortex, which indicates that their expression is regulated by temporal and spatial transcription factors. (B) *hth* is expressed in the glial cells that are born during the *tll* temporal window of the OPC neuroblasts as shown in the UMAP plot (green arrow), the Novosparc reconstruction (green arrows) and by HCR-FISH against *hth* (green arrows). (C) Novosparc-reconstructed expression of temporally regulated terminal selectors. We use the 3D reconstruction to identify all concentric genes (whose expression is regulated by the temporal transcription factors) and to classify whether the neurons that express these genes are born early (such as *bsh-* and *Lim3-*expressing neurons), mid (*vvl-* and *dve-*expressing neurons), or late (*hbn-* and *ab-*expressing neurons). (D) Novosparc-reconstructed expression of spatiotemporally regulated terminal selectors. We use the 3D reconstruction to identify all terminal selectors which are spatially restricted in the medulla cortex, which indicates that they originate from one (or more) of the spatial domains, such as *bab2* (dVsx domain), *mirr* (vVsx and vOptix domains), and *ara* (vtOPC). (E) Novosparc-reconstructed expression of terminal selectors with complex regulation. We identify terminal selectors whose expression pattern is more complex, such as *Sox21B*, which is expressed in neurons from two temporal windows in a spatially restricted manner, and *tup,* which is also is expressed in neurons from two distinct temporal windows.

As mentioned earlier, we also noticed that *hth* expression was not only restricted in the first temporal window of the medulla neuroblasts and its progeny, but the 3D reconstruction suggested extensive expression in glia (**Figure 6B**), which was also supported by the UMAP plot. To verify the presence of the *hth* transcript in glia, we performed HCR-FISH against *hth*. Indeed, we verified the presence of *hth-* expressing cells at the same location as predicted by the 3D reconstruction.

### Novosparc predicts complex regulation of candidate terminal selector expression

Terminal selectors are transcriptional regulators that coordinate the acquisition and maintenance of neuron-type-specific terminal features, including neurotransmitter identity, ion channel expression, and synaptic connectivity (Hobert, 2008; Hobert, 2011). Their presence has been shown predominantly in *C. elegans;* however, candidate terminal selectors have also been published in the *Drosophila* optic lobe (Özel et al., 2022; Simon et al., 2025). We recently noticed that concentric transcription factors (which are neuronal transcription factors regulated by temporal transcription factors, which results in a concentric expression in the medulla cortex (Hasegawa et al., 2011)) are all predicted terminal selectors (Simon et al., 2025), highlighting the crucial role of temporal patterning in establishing the great majority of neuronal diversity in this neuronal structure. We wondered whether we could use the 3D reconstruction of the optic lobe to a) identify the expression pattern of all published candidate terminal selectors and categorize them and b) identify the relative expression of concentric genes with respect to the neuroblast temporal windows.

Indeed, by plotting the expression of all 95 candidate temporal selectors (**Figure S5**):

- we noticed that about one third of them (31) form concentric rings in the medulla cortex, suggesting that they are under the regulation of temporal transcription factors,
- 18 of them have specific spatiotemporal patterns indicating that they are regulated by both spatial and temporal transcription factors,
- 9 show even more complex expression patterns raising the possibility that they are regulated by multiple temporal and/or spatial transcription factors,
- 9 are mostly expressed in specific neuropils,
- 16 have restricted expression patterns in single progenitor or neuronal types, while
- 12 have broader, albeit specific, expression patterns.

Importantly, we noticed that by looking into the spread of the predicted gene expression in the anteroposterior axis, we could predict whether a gene was expressed in neurons that are generated during early, mid, or late temporal windows, as exemplified by the known early, mid, and late markers, *bsh*, *vvl*, and *hbn* respectively (Konstantinides et al., 2022) (**Figure 6C**). Similarly, looking at the distribution of the gene expression along the medulla cortex, we could predict the domains where each of the neurons that express these genes come from. For example, *bab2* is expressed in neurons that originate from the dVsx domain, *mirr* is expressed in neurons that come from the vVsx and vOptix domains, and *ara* in neurons that come from the vtOPC (**Figure 6D**). Finally, genes that are under more complex regulation, such as *Sox21B* and *tup*, which are expressed in neurons from at least two different temporal windows can also be visualized (**Figure 6E**).

In total, this shows that our 3D atlas can serve as a reliable hypothesis generator and can facilitate the study of different stages and different mechanisms of neuronal development.

### A dedicated webpage to explore the three-dimensional atlas

To maximize the use of this atlas, we made this three-dimensional reconstruction available to the scientific community via a dedicated webpage (https://larva3dnovosparc.ijm.fr). The webpage layout consists of several components:

- dropdown menus to select the genes to be visualized (up to 3 genes can be visualized at once).
- color pickers to choose the color for each gene.
- threshold inputs for each gene to adjust the range of values for visualization.
- a removal threshold for filtering out low-expression data.
- axis filters (x, y, z) via sliders to perform virtual sections along each of the three axes.
- a download option to export the filtered dataset as a CSV file.

The main feature of the webpage is a 3D scatter plot that allows the visualization of the data points in space, colored by the presence or absence of selected gene expression. Users can blend colors for up to 3 genes and adjust the range of values to be visualized by setting minimum and maximum thresholds for each gene. The plot updates dynamically based on user input. The axes can be filtered according to the selected ranges, and colors are updated in real time. Finally, the user can download the filtered data in a CSV format. The basic functionalities of the webpage can be seen in **Supplementary Figure S6**.

## Discussion

### How to make a neuronal three-dimensional gene expression atlas?

Here, we present the pipeline that we followed to generate a three-dimensional gene expression atlas of a complex developing neuronal structure, the *Drosophila* optic lobe, using Novosparc. This computational approach allowed us to infer the spatial distribution of cell types and gene expression, providing a valuable resource for studying the organization of the developing nervous system. By leveraging single-cell mRNA sequencing data and spatial transcriptomic reconstruction, we identified spatially compartmentalized cell types, detected patterns of transcription factor expression, and predicted regions of programmed cell death. This work highlights the power of computational methods in augmenting single-cell sequencing datasets and provides insights into the spatial organization of neuronal development.

We also present the necessary steps needed to perform the reconstruction of a complex neuronal structure. First and foremost, one needs to take into account the developmental origin of the different neuronal types. While the hypothesis that cells that are close to each other are more likely to resemble each other transcriptomically is on average correct, it falls short when structures with different developmental origins juxtapose each other; cells with shared lineage tend to localize together, suggesting that spatial organization is not solely dictated by signaling cues but also by intrinsic lineage constraints. Therefore, to perform a 3D reconstruction with Novosparc one should divide the tissue in regions based on their developmental origin. Then, we identified markers that would separate these regions, as well as some of the cell types that we already knew in our tissue of interest. Finally, we tested different algorithm parameters, although this had the least impact in the resulting reconstruction.

Finally, we provide further proof that Novosparc is an efficient tool to generate 3D reconstructions. Barring some significant limitations that are described below, Novosparc managed to reconstruct a complex neuronal structure with significant accuracy, using the expression of only 43 genes as landmarks. We believe that Novosparc provides complementary information to spatial transcriptomic approaches, as it uses the entire transcriptome to assign cells to positions instead of a few hundreds of marker genes that are usually used in standard spatial transcriptomic approaches. Moreover, it can achieve single cell resolution, as we can have equal number of points in the reconstructed object as cells.

### Study limitations

Novosparc presents also limitations compared to other spatial transcriptomic approaches. First, it relies heavily on existing knowledge: in the absence of any landmark expression information or developmental origin knowledge, Novosparc is not performing satisfactorily in complex tissues. As a corollary, this limits its capacity to identify new “regions” that would deviate from the rule that physical proximity correlates with transcriptomic similarity. Second, using Novosparc we are restricting each cell to a single vertex. While this may not be a problem for different tissues, neurons are very complex cells with great neurite diversity, which often translates to localization of specific transcripts in different subcellular locations (Glock et al., 2021; van Oostrum and Schuman, 2025). This type of information is completely inaccessible in a Novosparc reconstruction and requires other spatial transcriptomic approaches for this complexity to be uncovered. Finally, Novosparc does not predict absolute expression levels but rather reconstructs spatial gene expression by assigning probabilities of expression to different locations. This represents a limitation because it does not directly capture the quantitative differences in gene expression across the tissue. Instead, it provides a probabilistic estimate of where a gene is likely to be expressed based on spatial constraints and similarity to reference data. As a result, while Novosparc can reveal spatial trends and organization, it may not accurately reflect expression gradients or absolute transcript abundances, potentially limiting its application in cases requiring precise quantitative measurements.

One final limitation of our study is that, while we tried to reconstruct the entire developing optic lobe, our focus was predominantly the neurons of the medulla neuropil that originate from the main OPC. This means that we have lower spatial resolution in lamina neurons and the neurons that come from the IPC, as well as glia, which were essentially only assigned to two regions: the last medulla temporal window and the wrapping glia region. Nonetheless, we do provide the pipeline, the know-how, and the code for future studies to elaborate further on either of these two populations.

### Temporal and spatial patterning of programmed cell death and terminal selector expression

Despite the above limitations, we were able to use the atlas to study programmed cell death and terminal selector regulation. Programmed cell death is a critical mechanism ensuring proper neuronal diversity and numbers during development (Pinto-Teixeira et al., 2016). Our 3D reconstruction of the *Drosophila* optic lobe revealed that the expression of key proapoptotic genes, *reaper, grim,* and *skl*, is not random but instead follows specific spatial and temporal patterns. These genes are expressed in defined neuronal populations, suggesting that programmed cell death is tightly regulated by developmental specification mechanisms. In contrast, *hid* expression was observed broadly, including in neurons that do not undergo apoptosis, hinting at a potential non-apoptotic role or stress response function. The spatially restricted expression of *reaper, grim*, and *skl* aligns with previous findings that neuroblast lineage and temporal transcription factors influence cell fate decisions, including apoptosis (Bertet et al., 2014) and raises evolutionary hypotheses regarding the role of programmed cell death in neuronal type evolution (Prieto-Godino et al., 2020). These findings reinforce the idea that temporal and spatial transcriptional programs not only define neuronal identity but also dictate survival, ensuring the proper assembly of the optic lobe’s complex circuitry.

Moreover, our 3D reconstruction of the *Drosophila* optic lobe revealed that terminal selector gene expression is closely linked to temporal transcription factor dynamics. Specifically, a significant fraction of candidate terminal selectors follows concentric expression patterns in the medulla cortex, suggesting regulation by temporal transcription factors. Additionally, genes with more complex or spatially restricted expression patterns indicate that some terminal selectors integrate both spatial and temporal cues. By analyzing gene expression along the anteroposterior axis, we could predict whether neurons expressing these genes originated from early, mid, or late temporal windows. Similarly, the medial-lateral spread of gene expression in the medulla cortex allowed us to infer neuroblast domain origins. These findings reinforce the idea that temporal transcription factors play a crucial role in coordinating neuronal diversity by guiding terminal selector expression. Furthermore, genes with complex regulatory patterns, such as *Sox21B* and *tup*, highlight the existence of multi-layered regulatory mechanisms. Our results demonstrate that the 3D spatial reconstruction of the optic lobe is a powerful tool for dissecting developmental gene regulation, providing a framework to investigate how temporal and spatial factors interact to shape neuronal identity.

### Evolution of developing nervous systems

The know-how gained from reconstructing spatial gene expression can be applied to studying the evolution of developing nervous systems by enabling cross-species comparisons of spatial organization. With the increasing availability of single-cell sequencing datasets across diverse species, we now have an unprecedented opportunity to investigate how neurodevelopmental programs, such as spatial and temporal patterning (Filippopoulou and Konstantinides, 2023), have evolved. By using spatial reconstructions of single-cell data, we can identify conserved and divergent patterns of gene expression, cellular arrangements, and developmental trajectories across taxa. This approach can reveal how lineage constraints shape nervous system architecture and how novel structures emerge through evolutionary modifications (Roberts et al., 2022; Tosches, 2017). Additionally, comparing the spatial distribution of homologous genes across species can help disentangle the relative contributions of conserved regulatory networks and species-specific adaptations. The ability to reconstruct and analyze spatial gene expression in a three-dimensional context further enhances our capacity to study neuronal diversification at multiple levels, from gene regulatory dynamics to whole-brain patterning. As more single-cell and spatial datasets become available, comparative studies will provide deeper insights into the fundamental principles that govern nervous system development and its evolutionary transformations.

## Materials and Methods

### *Drosophila* husbandry and genetics

Fly stocks and crosses were maintained on standard food at 25°C for experiments. The *Drosophila melanogaster* stocks used can be found in Supplementary Table S3.

### HCR-FISH

The HCR protocol for *Drosophila* larval brains used was as specified previously (Ferreira et al., 2021).

### Antibody staining

Wandering third instar larval *Drosophila* optic lobes were fixed in 4% formaldehyde for 15–20 min at room temperature. After washing, the samples were incubated for 2 days with primary antibodies at 4 °C. After washing the primary antibodies, the brains were incubated with the secondary antibodies overnight at 4 °C. The secondary antibodies were washed, and the brains were mounted in Slowfade and imaged at a confocal microscope (Leica SP8) using a ×63 glycerol objective. Images were processed in Fiji.

Details of the origin of all individual antibodies can be found in Supplementary Table S3.

### Novosparc - Blender

To build the atlas, we used the 3D software Blender. We imported CSV files of coordinates via the scripting tab in Blender and a Python script. Once imported, we assigned the xcoord, ycoord, and zcoord columns of the CSV file to the x, y, and z positions in Blender using Geometry Nodes. We first added a “Set Position” node between the input node and the output node. Then, we added a “Combine XYZ” node connected to the position socket of the “Set Position” node. We connected each socket of the “Combine XYZ” node (x, y, and z) to a “Named Attribute” node. We selected either xcoord, ycoord, or zcoord for each attribute, depending on which socket it was connected to. Finally, we applied the Geometry Nodes Modifier.

Layer by layer, we selected the points we wanted to associate with an atlas region and separated them into a new mesh, which we named according to the color associated with that region. After doing this for all the points, we exported the Blender file to an OBJ file that included all the meshes and their coordinates. Then, we created a new CSV file containing all the positions and added new columns indicating which mesh each position belonged to. For example, if a coordinate was in the "blue" mesh, we wrote 1 in the blue column; if not, 0. We applied the same method to generate all subregions.

### Novosparc – R

Then, we prepared the published data for Novosparc. First, we needed to switch the published RDS object to an AnnData object that contained the normalized but not scaled data in the correct location. An AnnData object has multiple layers; Novosparc accesses a specific layer of this object (the primary layer), so it is crucial to ensure that the desired data for reconstruction is stored in that layer. To achieve this, we read the .rds dataset into R as a Seurat object, which we then converted to an AnnData object, ensuring the appropriate data was placed in the primary layer. The script Conversion_To_AnnData_Only_Normalized can be found in the GitHub repository.

### Novosparc - Overview of the Algorithm and Workflow

The Novosparc algorithm is based on the hypothesis that there is a correspondence between the structure of positions in physical space and the structure of cells in gene expression space. Therefore, there is a correspondence between pairwise distances of locations in physical space and pairwise distances of the cells in gene expression space. This is why the first step of the Novosparc algorithm is to compute cost matrices based on the construction of k-nearest-neighbors (kNN) graphs in both physical space and gene expression space. Using the first two inputs, the locations file (CSV file of coordinates) and the single-cell dataset, Novosparc computes two cost matrices: the cell-cell cost matrix and the location-location cost matrix. These consist of the pairwise distances between cells andlocations, computed as the shortest path according to the kNN graph. As we use another input here, the reference atlas (locations file with marker genes’ binarized expression), Novosparc computes an additional cost matrix: the cell-location cost matrix. This matrix captures the discrepancy between the expression of the marker genes in each cell and in each location of the target space.

Using these cost matrices, Novosparc computes the optimal transport cost matrix, which probabilistically assigns each cell to a location. This assignment depends on different parameters, such as the alpha parameter, which allows us to decide the importance we want to assign to the reference atlas. A value of alpha near 1 indicates high fidelity to the reference atlas, while a value near 0 reflects higher fidelity to the de novo reconstruction (with alpha=0 being the de novo reconstruction).

The optimal transport matrix allows us to map the single-cell dataset to the locations file. We can then export the final file to a CSV file, with the first three columns being the coordinates (xcoord, ycoord, and zcoord) and all the other columns containing the gene expression data. Each row represents a cell, and each column (except for the first three) represents a gene.

The dataset used to compute the optimal transport matrix consisted of the gene expression values of the 5000 most variable genes, as using the most variable genes has been proven to improve the Novosparc reconstruction (Moriel et al., 2021).

### Novosparc - fine tuning of Novosparc parameters

When running Novosparc, there are several parameters that can be adjusted depending on the specifics of the reconstruction:

- **Alpha parameter**: This parameter controls the interpolation between two modes of reconstruction: de novo spatial reconstruction (α = 0) and reconstruction based solely on the information provided by the reference atlas (α = 1).
- **Number of genes**: This defines the number of genes used for the reconstruction. It is generally advisable to include only the most differentially expressed genes, as these are more likely to be spatially informative (i.e., expressed in some cells but not others, ideally across a well-defined spatial domain). However, the markers used for reconstruction are not always the most differentially expressed genes. For instance, if a marker is among the top 5000 differentially expressed genes, you must include at least the top 5000 genes in the reconstruction to incorporate that marker. Balancing this parameter is crucial for achieving accurate results.
- **Number of k-neighbors**: This parameter determines the number of neighbors used to construct the k-nearest neighbors (kNN) graphs for cells and spatial locations.
- **Epsilon**: This parameter is associated with the entropic regularization term. A low epsilon value leads to more localized mapping, while a higher epsilon value results in higher-entropy mappings, approaching a uniform distribution.

One of the challenges we encountered was measuring the reconstruction efficiency in an unbiased way. The goal of this fine-tuning was to run Novosparc with a set of different parameters and, based on the quality of the results, select the optimal parameters to use. To assess the efficiency of the reconstruction, we first chose a set of genes (*acj6, dpn, eya, toy, shg, elav, bsh, ap, hth, ey, slp1, tll, opa, tj, hbn, pxb, Optix, Vsx1, Rx,* and *wg*) and visualized their localization using the embedding function from Novosparc. The efficiency was initially evaluated through visual observation of the reconstructed gene expression across 20 z-stacks.

Additionally, for each set of parameters, the entropy of the transportation of individual cells was measured, and the different parameter sets were compared using histograms and boxplots (**Supplementary Figure S3**). Lower entropy was associated with a more efficient mapping. To ensure consistency, only one parameter was varied at a time. We established a default set of parameters: alpha = 0.5, number of genes = 2000, k neighbors = 15, and epsilon = 5e-3. We then varied each parameter individually.

#### a) Alpha parameter

The reconstructed images were quite similar, making it difficult to make a decision based solely on them. However, an analysis of individual cell entropies revealed a clear trend: entropy decreases as the alpha parameter increases, at least up to an alpha value of 0.5 (**Supplementary Figure S3A**). Based on this observation, we decided not to use alpha values of 0.1 and 0.3. Ultimately, we chose to proceed with an alpha value of 0.5, aiming for a reconstruction that incorporates a balance between de novo and marker-based approaches.

#### B) Epsilon parameter

This parameter was the most complicated to evaluate. Epsilon directly influences entropy, with lower epsilon values resulting in lower entropy. Also, using low epsilon values significantly increased computation time, with the code taking two days to run for epsilon 0.001 and even longer for epsilon 0.0005, compared to just 8–10 hours for the other values.

As expected, lower epsilon values resulted in lower entropy values (**Supplementary Figure S3B**). We decided to keep epsilon = 0.005 for a balance between timing and reconstruction efficiency.

#### C) K-Nearest neighbors

In the Novosparc code, there are two parameters defined for nearest neighbors: num_neighbors_s and num_neighbors_t, which are typically set to the same value, for example: num_neighbors_s = num_neighbors_t = 15. These parameters assume the existence of an expression “niche” or microenvironment around a cell, where expression differences within the niche are subtle.

- num_neighbors_s is chosen based on the number of immediate neighbors of a point, which depends on the grid’s dimensionality. In our case, the grid is 3D, so this value falls between 17 and 26.
- num_neighbors_t can be adjusted based on factors such as the total number of cells, levels of noise, and the dimensionality of the grid. The Novosparc paper suggests that values for num_neighbors_t between 3 and 15 result in robust spatial embedding.

The entropy analysis showed almost no differences when using different k-nearest neighbors parameters (**Supplementary Figure S3C**).

#### D) Number of genes

Upon reviewing the images, we observed that the reconstructions most closely resembling reality were those using higher gene counts, specifically 5000 and 8000 genes. This observation was further supported by entropy analysis, which showed a decreasing entropy value as the number of genes increased (**Supplementary Figure S3D**). Based on this analysis, we decided to use 5000 genes for the reconstruction.

### Novosparc – final code

The code used to run Novosparc can be found in the GitHub repository (**Novosparc_End**).

### Novosparc - website

To visualize the data a website was developed; the code is available in GitHub (**Web App**). This Python script implements a web application for visualizing and interacting with the Novosparc reconstruction. The application, built using Dash, allows users to select genes, apply thresholds, and choose colors to represent gene expression levels in a 3D scatter plot. Additionally, users can filter the data along the x, y, and z axes and download the filtered dataset.

### Key Features

1. Google Cloud Storage Integration: The script begins by downloading a CSV file from a Google Cloud Storage bucket using the google.cloud library. The file contains reconstructed single-cell data with coordinates and gene expression levels, is saved to the local file system and then read into memory in chunks to manage large files. This file contains the results of the Novosparc code. Several prior operations were performed on this file: first, reuniting the xyz data, which is not initially included in the file. To achieve this, the xcoord, ycoord, and zcoord columns from the atlas are merged with the file. Next, the data in the file is linearly normalized to a range of 0 to 1, ensuring that no values have more than three decimal places. This normalization facilitates visualization in the code and reduces the size of the CSV file. The code used for normalization is available in GitHub (**Linear_Normalization**); the code used for reducing the numbers of decimals is also available (**Less_Decimals**).
2. Data Preprocessing: The data is loaded and processed, with coordinates (xcoord, ycoord, zcoord) and gene expression values extracted. The script prepares options for gene and color selection, allowing the user to overlay gene expression values with different color schemes (e.g., greyscale, yellow, green, etc.).
3. Dash Web Interface: The application layout consists of several components:

- Dropdown menus to select the first and optionally the second gene.
- Color pickers to choose the color for each gene.
- Threshold inputs for each gene to adjust the range of values for visualization.
- A suppression threshold for filtering out low-expression data.
- Axis filters (x, y, z) via sliders to zoom in on specific regions of the 3D space.
- A download button to export the filtered dataset as a CSV file.
4. Dynamic 3D Plot: The main feature of the website is a 3D scatter plot that visualizes the data points in space, colored by the selected genes. Users can blend colors for three genes and adjust the color intensity by setting minimum and maximum thresholds for each gene.
5. Interactivity: The plot updates dynamically based on user input. The axes are filtered according to the selected ranges, and colors are updated in real time. The second gene visualization can be toggled on/off via a checkbox.
6. Data Download: The app includes a feature to download the filtered data, which is triggered by a button. When clicked, the filtered dataset is converted to CSV format and made available for download.

This Dash-based application provides an interactive platform for visualizing and analyzing the Novosparc reconstructed data. Users can customize gene selection, color schemes, and visualization thresholds, and easily export the filtered data for further analysis.

## Acknowledgements

We thank the members of the Konstantinides lab and, in particular, Elisavet Iliopoulou for valuable discussions. We also thank Nikos Karaiskos and Shuonan He for help with the implementation of Novosparc and Volker Hartenstein for sharing images of the developing fly brain. Finally, we thank Joel Marchand for informatic assistance during the implementation of the project. This work is supported by the European Research Council (ERC) under the European Union’s Horizon 2020 research and innovation programme (grant agreement No. 949500) and the HORIZON-WIDERA-2023-ACCESS-02 grant no. 101159925 - SCENTINEL. I.H. is supported by a Marie Skłodowska-Curie Postdoctoral Fellowship (101154260).

## Data and resource availability

All data, related code, as well as the final resource are available in GitHub (https://github.com/leonardotadini/Leonardo-Scripts), GEO (under accession number GSE167266), and in the dedicated webpage (https://larva3dnovosparc.ijm.fr).

## Supplementary Figure legends

**Supplementary Figure S1: Single-cell sequencing of the developing optic lobe**

(A) UMAP plot of the single-cell mRNA sequencing experiment of the third larval instar developing optic lobe. The medulla, lamina, lobula plate neuronal types, as well as the glial clusters are color-coded. (B) UMAP plots showing the localization of the different neuronal types that come from the respective spatial domains.

**Supplementary Figure S2: Marker gene selection**

(A) UMAP plots showing the localization of the different central brain neuronal types, which were removed from downstream analyses. (B) UMAP plots showing the expression of gross cell types markers. *shg* marks neuro epithelium and neuroblasts, while wrapper marks non *repo*-expressing wrapping glia. (C) UMAP plots showing the expression of temporal and spatial transcription factors. Combinations of spatial and temporal transcription factors can differentiate between the different neuroepithelial regions, neuroblast and GMC temporal windows, as well as many neuronal types. (D) UMAP plots showing the expression of different neuronal transcription factors. Combinations of these markers can differentiate between the different neuronal types.

**Supplementary Figure S3: Parameter selection**

Novosparc offers the option to test different parameters. Their efficiency can be evaluated by plotting the entropy levels. The distribution and boxplot of entropy values of the transport matrices are shown in the upper and lower plot, respectively. We tested different values for the (A) alpha and (B) epsilon parameters, as well as for (C) k-nearest neighbors different and (D) different gene numbers. Asterisk indicates the selected parameter value.

**Supplementary Figure S4: Reconstructed marker gene expression**

(A) Novosparc-reconstructed expression of cell class markers as seen from a lateral (left) and dorsal (right) view. *Shg* is expressed in the neuroepithelium and neuroblasts, while *repo* is expressed in most glia and *wrapper* in wrapping glia. (B) Novosparc-reconstructed expression of spatial transcription factors, *Vsx1, Optix,* and *Rx* as seen from a lateral (left) and dorsal (right) view. (C) Novosparc-reconstructed expression of temporal transcription factors, *erm, ey, slp1, hbn, opa,* and *tll* as seen from a lateral (left) and dorsal (right) view. (D) Novosparc-reconstructed expression of neuropil markers, as seen from a lateral (left) and dorsal (right) view. (E) Comparison of reconstructed marker gene expression against immunofluorescence against the respective protein.

**Supplementary Figure S5: Reconstructed terminal selector expression**

Novosparc-reconstructed expression pattern of all candidate terminal selectors, as can be seen from a lateral (left) and dorsal (right) view. Their expression was used to categorize them to concentric genes, genes spatiotemporally regulated, genes with more complex regulation, neuropil-specific genes, genes with restricted gene expression, and broadly-expressed genes.

**Supplementary Figure S6: Basic functionalities of the website**

(A) Upon logging into the page https://larva3dnovosparc.ijm.fr, the user selects his first gene of interest. A calculation of the top20 most correlated genes (Spearman correlation) can be visualized. (B) A histogram of the expression distribution of the gene of interest is generated that allows the user to define visualization thresholds. (C) The user can then choose a specific color for the visualization of the first gene, as well as specific thresholds, based on the expression distribution. (D) If the user wants to visualize a second or third gene, they can enable their visualization and proceed with the same choices as before. (E) Axis sliders allow the user to perform virtual sections along each of the three axes: anteroposterior (x), dorsoventral (y), and/or mediolateral (z). (F) For all of these manipulations to take effect, the user has to “refresh scatter plot”. (G) A scatterplot that shows the distribution of expression of the gene of interest in the reconstructed 3D atlas is then displayed. The user can choose to remove the cells that don’t show expression (displayed in black) and rotate the brain using their mouse or the X, -X, Y, -Y, Z, and -Z buttons. The displayed data can then be downloaded as a CSV file using the dedicated button.

