## Supplementary Figures for "Gene expression cartography of a developing neuronal structure"

### Figure S1: Single-cell sequencing of the developing optic lobe

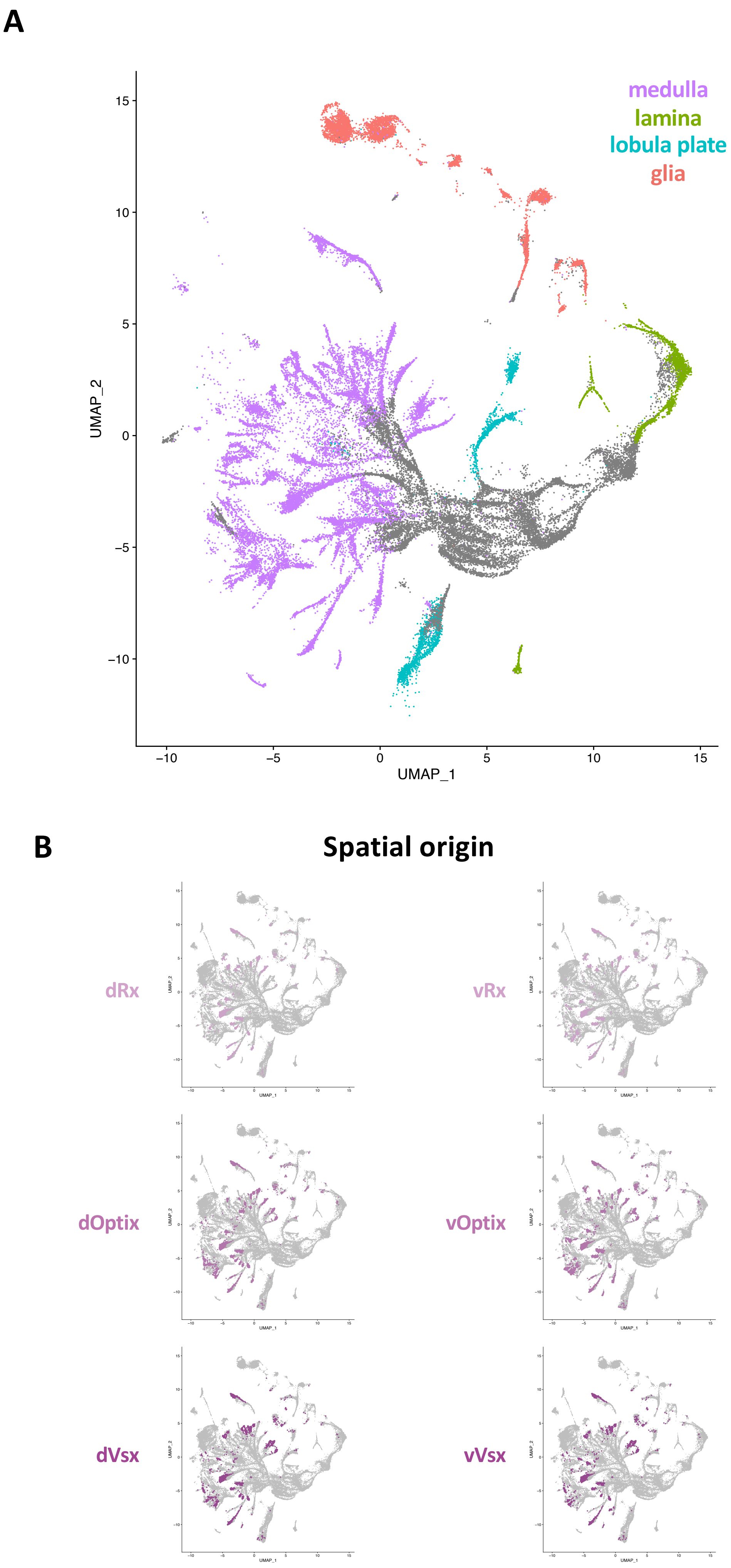

### Figure S2: Marker gene selection

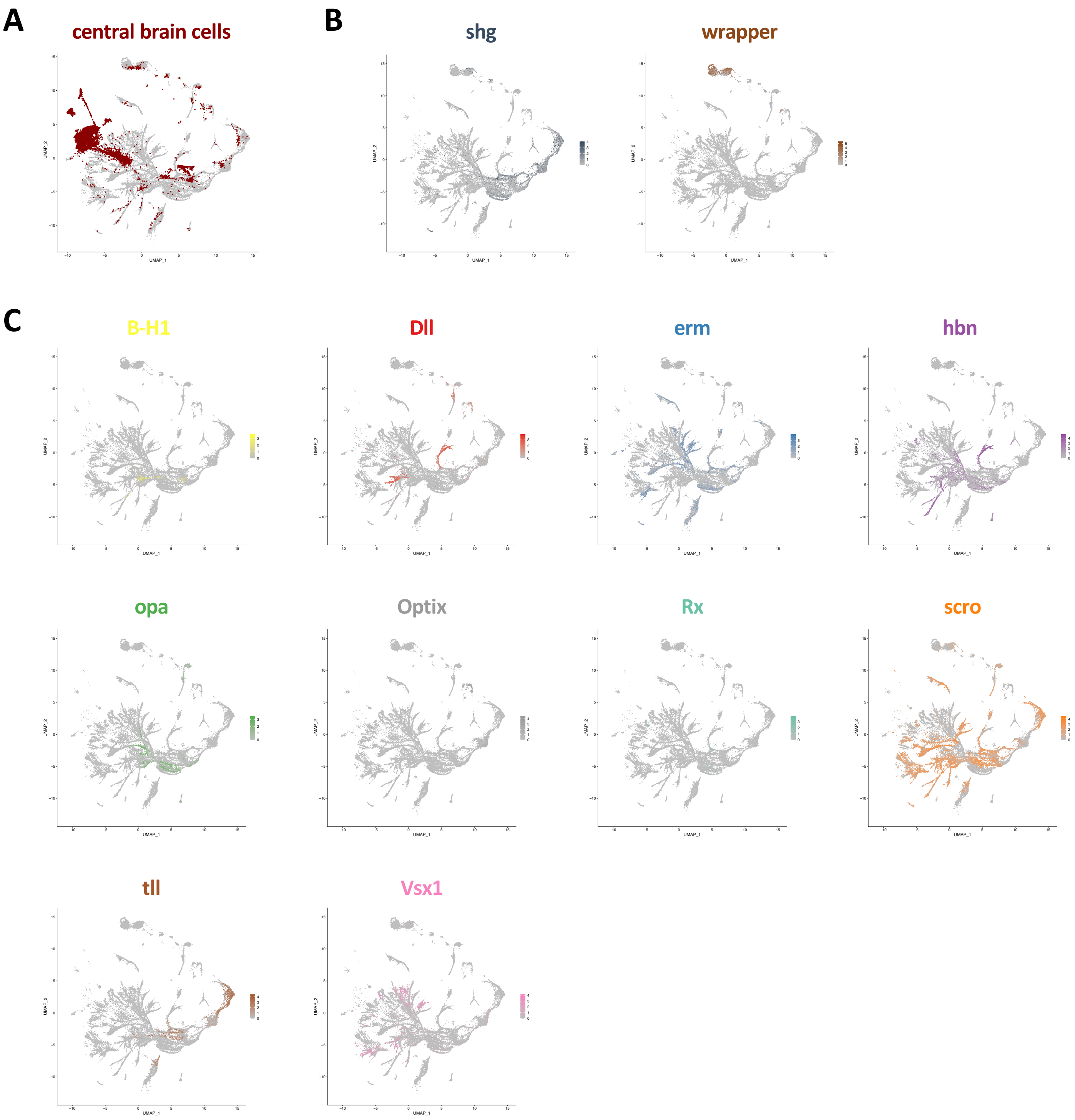

### Figure S2: Marker gene selection

D

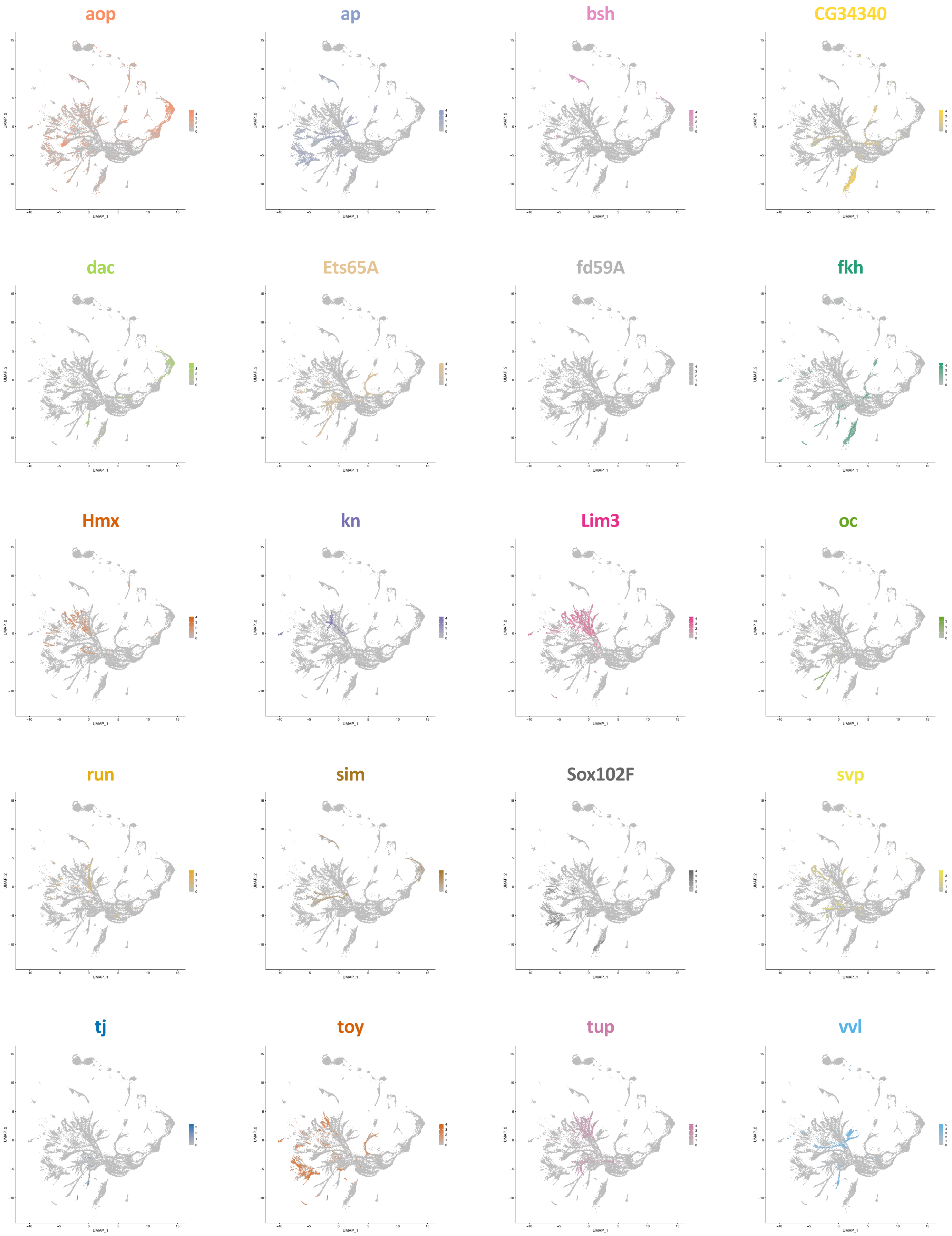

### Figure S3: Parameter selection

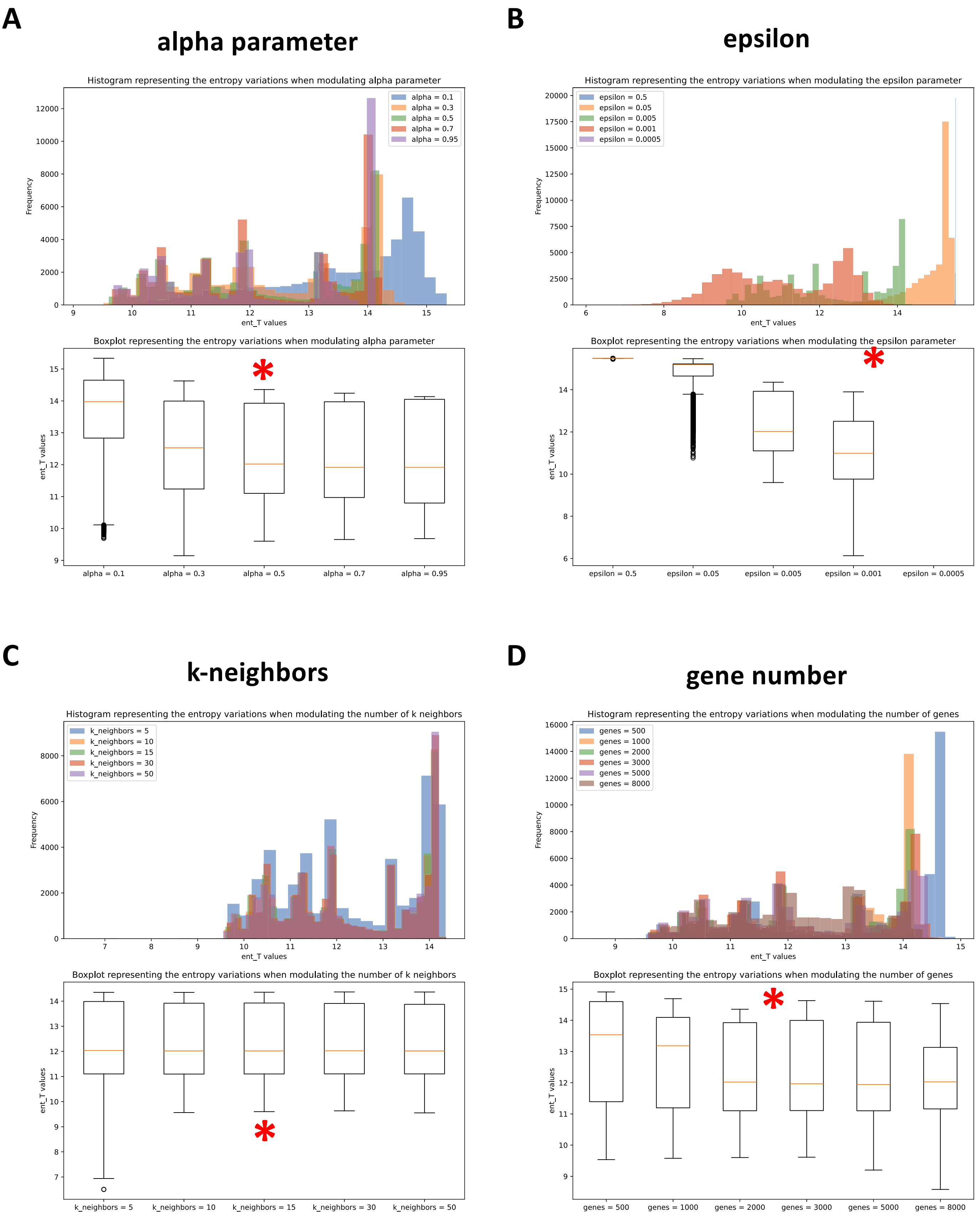

### Figure S4: Reconstructed gene expression of marker genes

**A** 

#### Gross cell type markers

***shg***

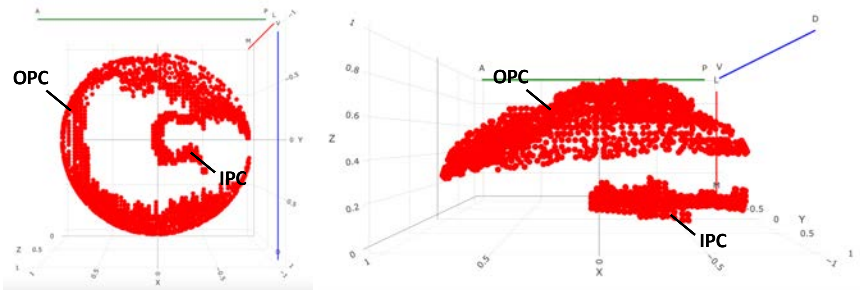


***repo***

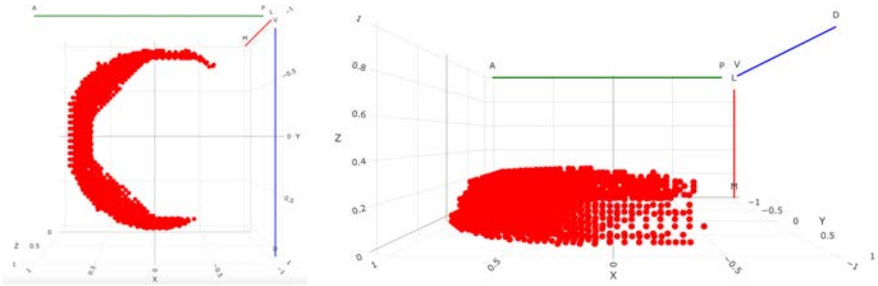

***wrapper***

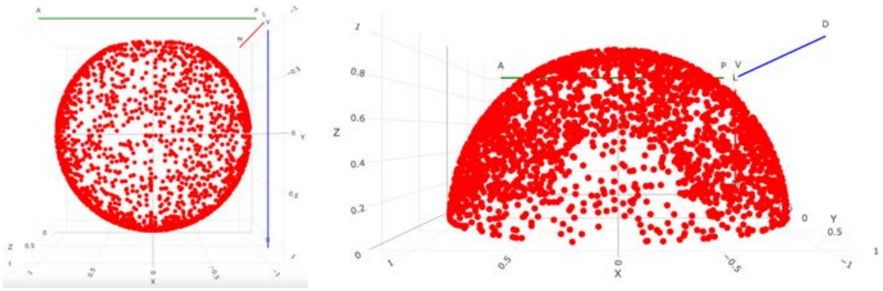

**B**

#### Spatial transcription factors

***Vsx1***

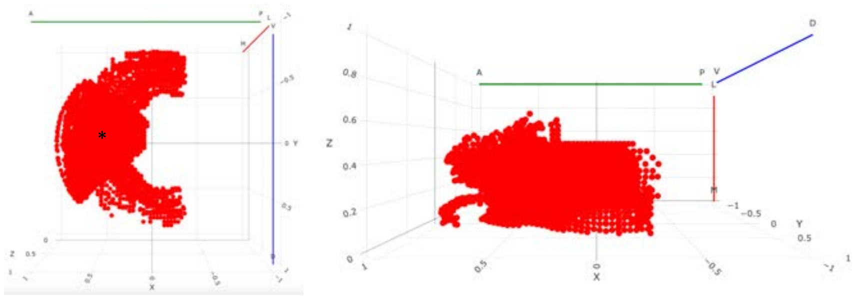

***Optix***

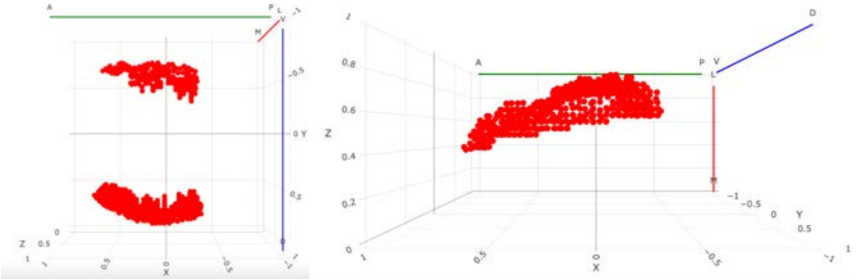

***Rx***

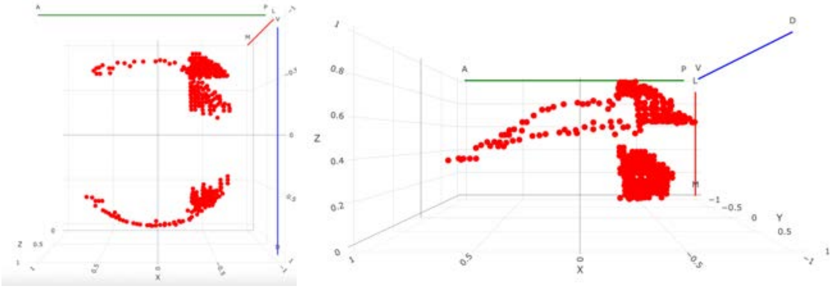

**C** **Temporal transcription factors**

***erm***

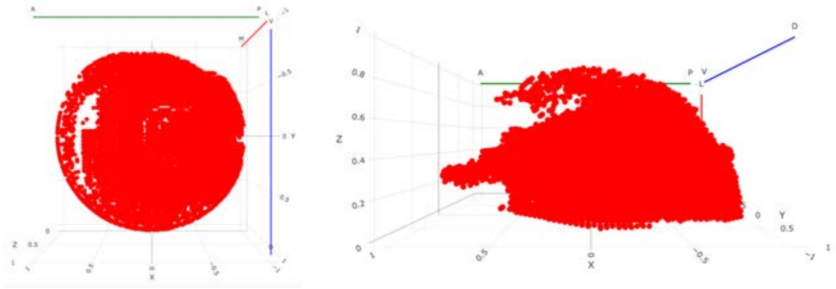

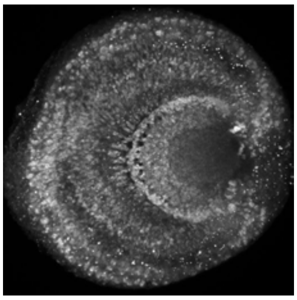

***ey***

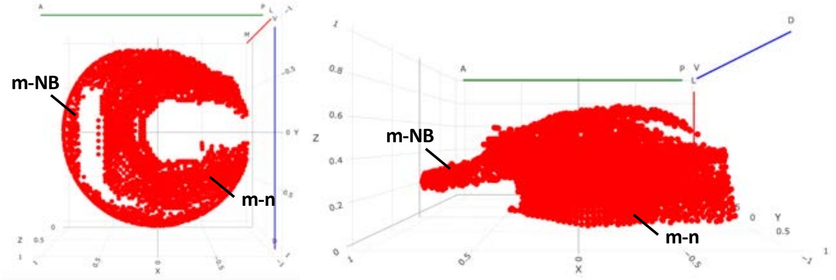

***slp1***

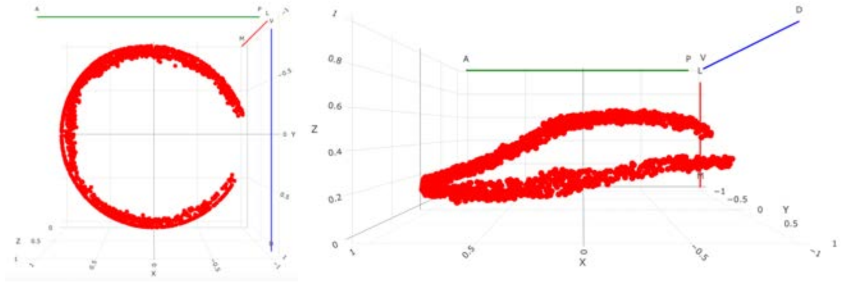

***hbn***

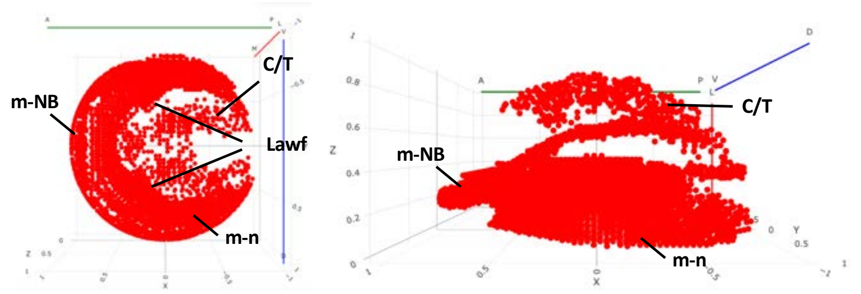


***opa***

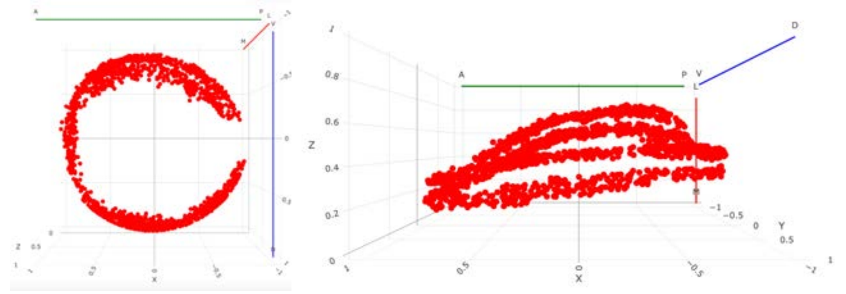

***tll***

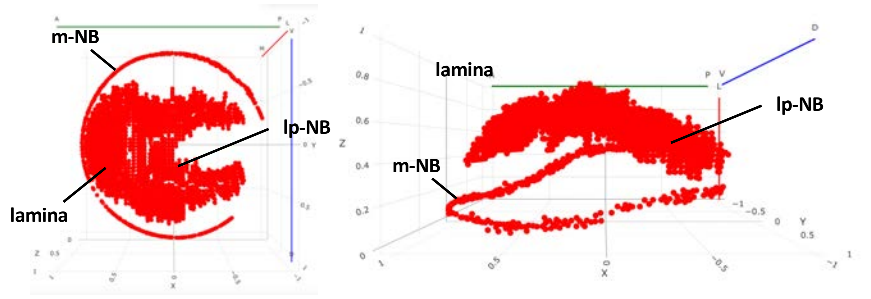


**D** **Neuropil markers**

***acj6***

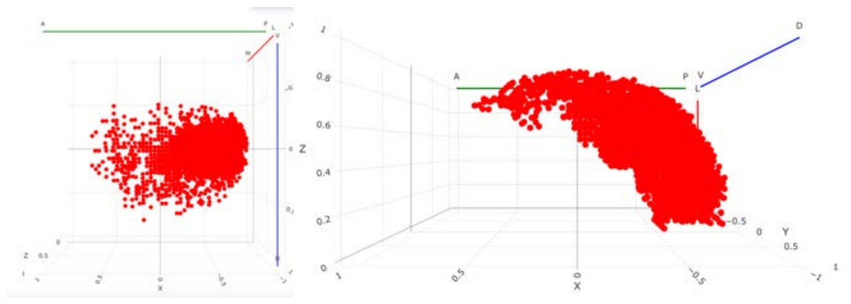

*eya*

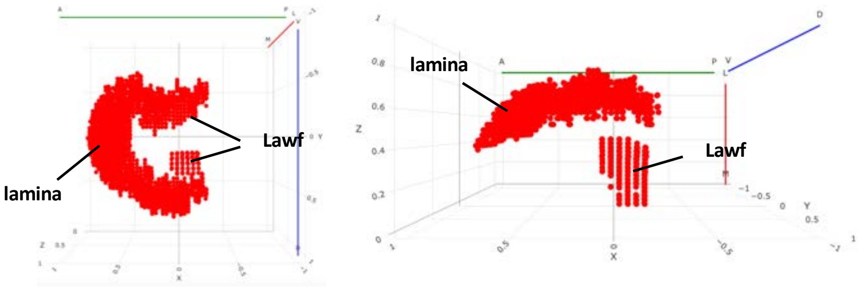


*Lim1*

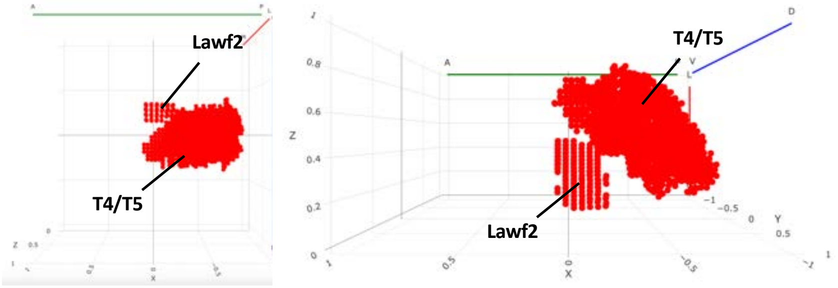


**E** **Neuronal type-specific genes**

*ap*

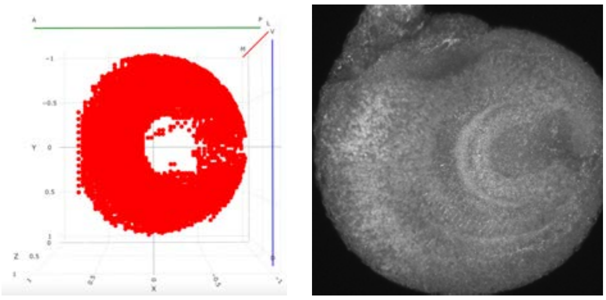

*dac*

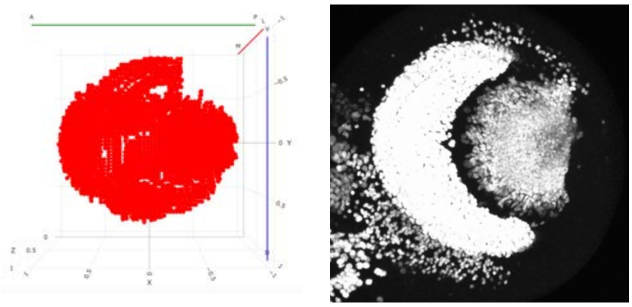

*dll*

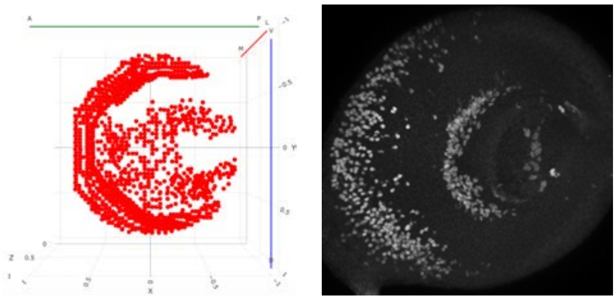

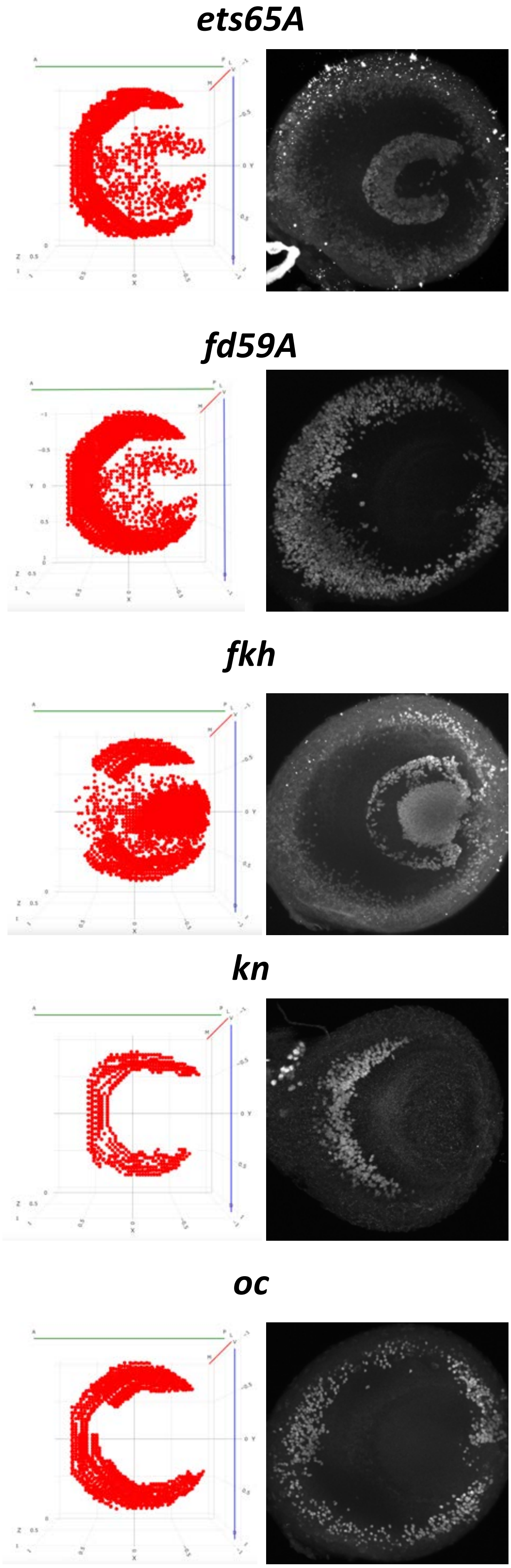
ets65A
fd59A
fkh
kn
oc

***svp***

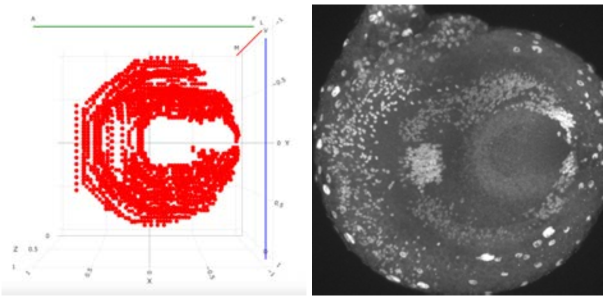

### Figure S5: Terminal selector expression

#### concentric genes

### ab

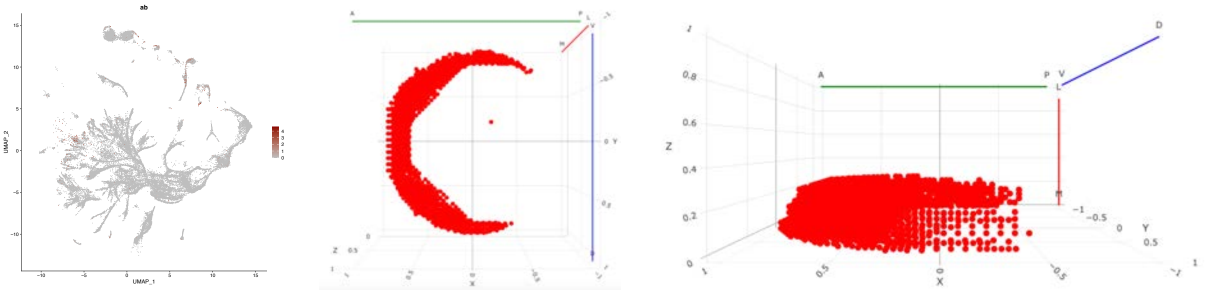

##### aop

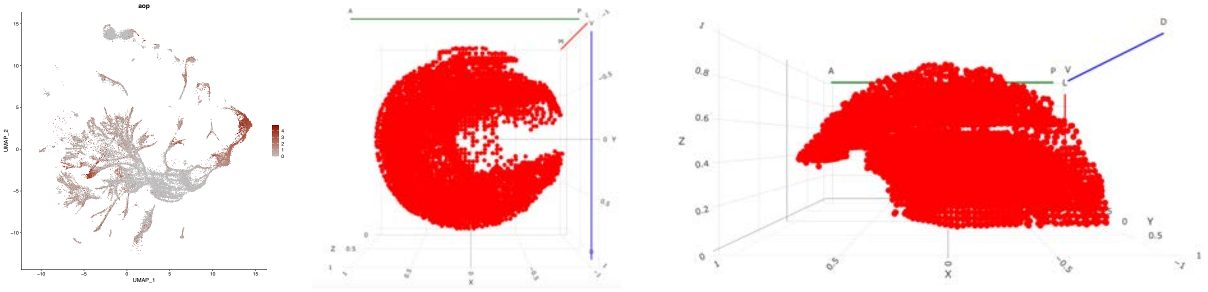

### ap

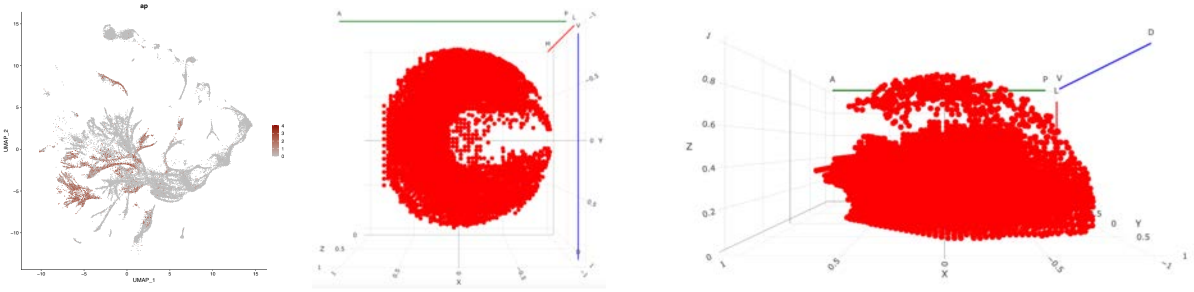

##### apt

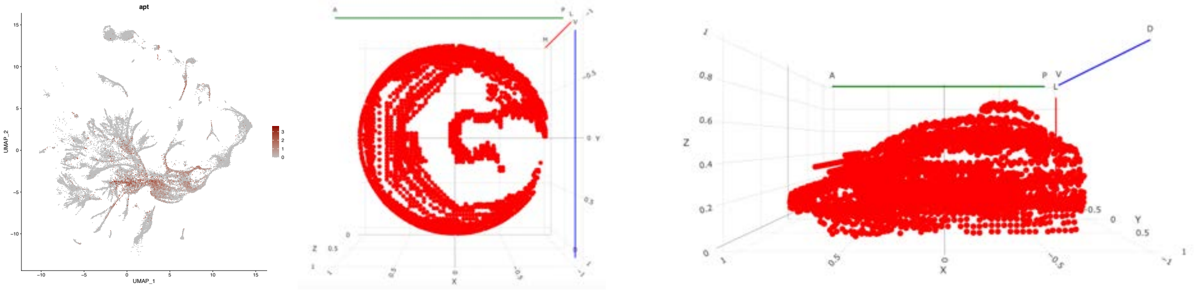

##### Atf3

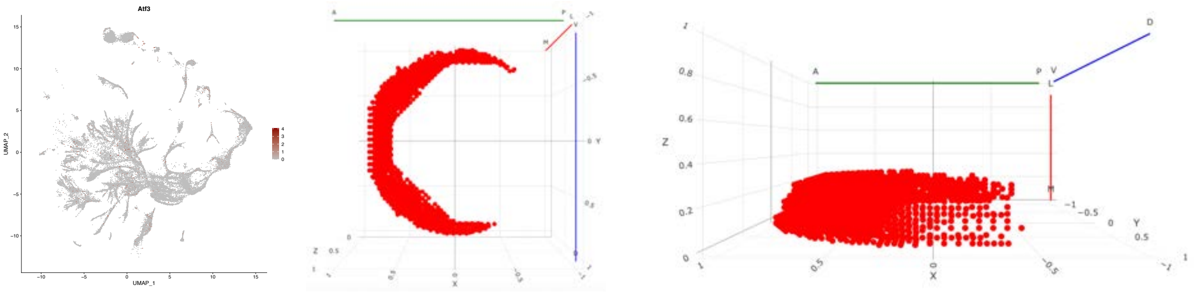

### bsh

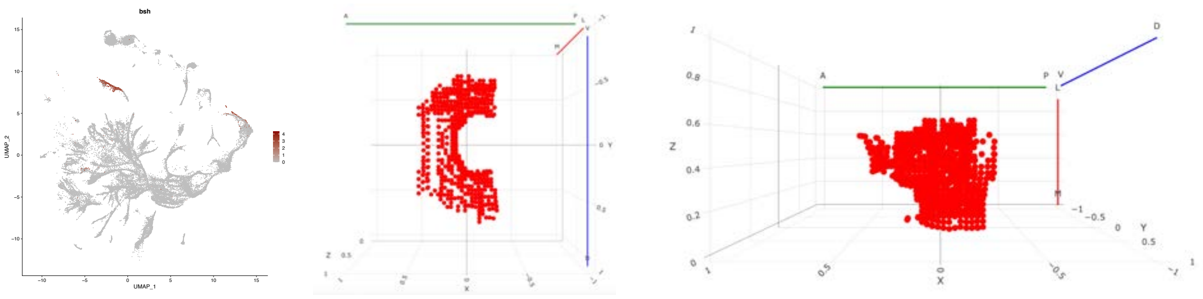


# CG11085

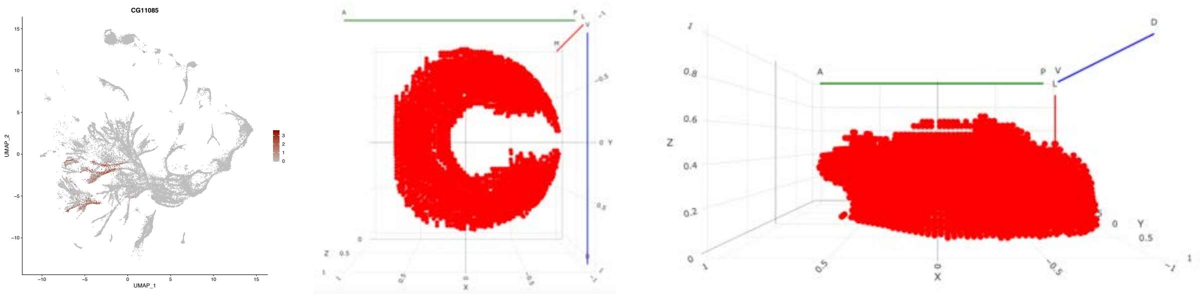


# CG11294

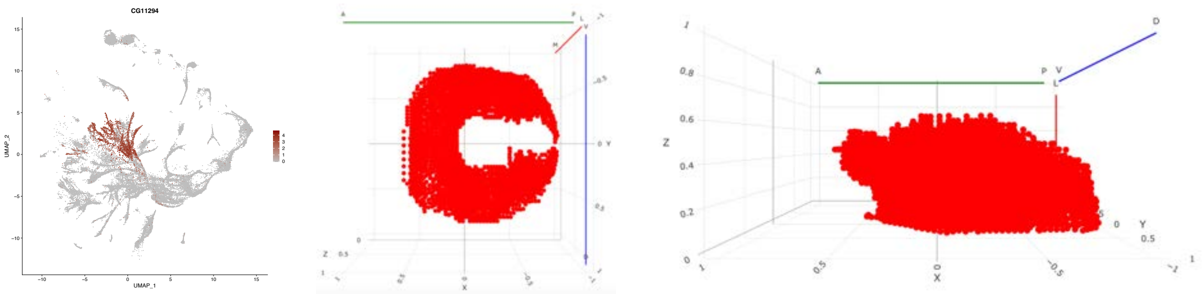


# CG43689

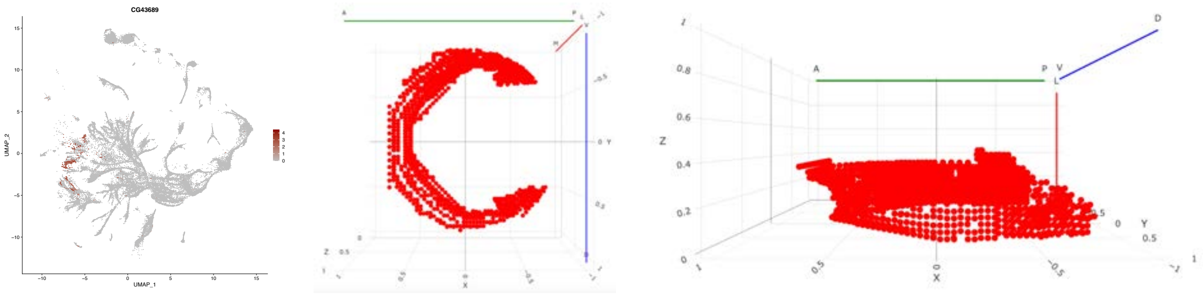


# dm

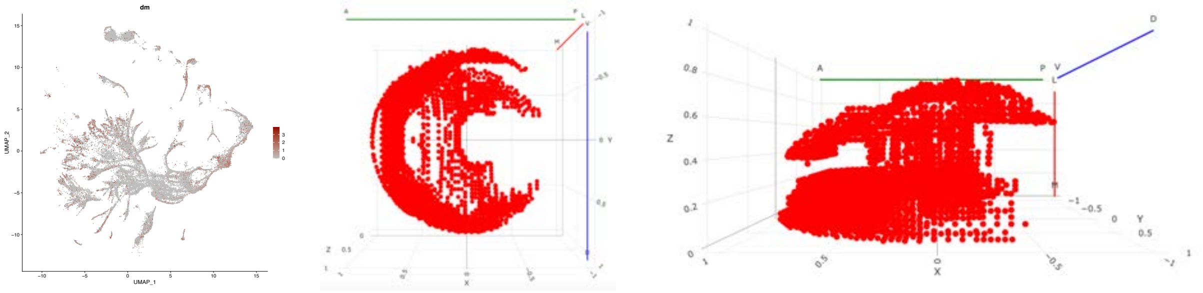

### dve

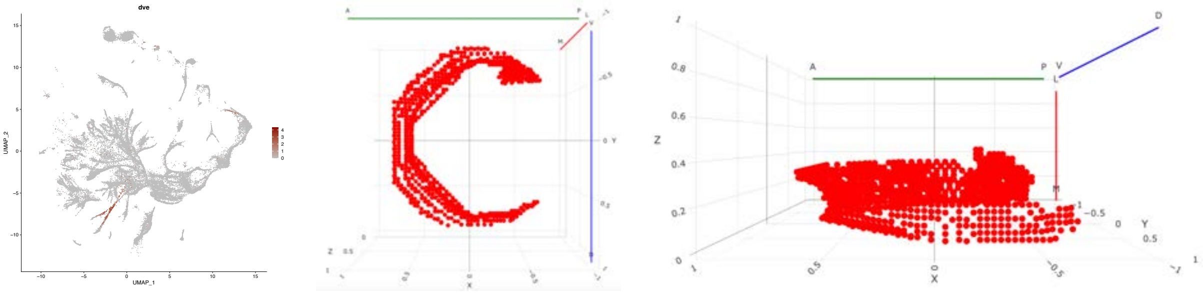


### elB

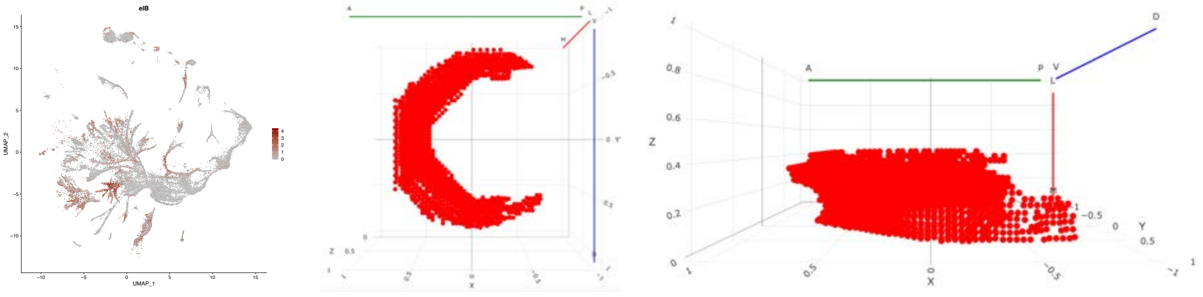


### ets65A

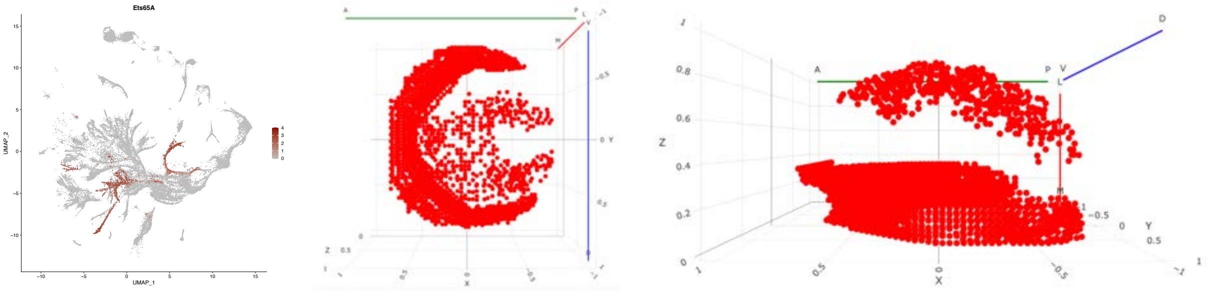


# ey

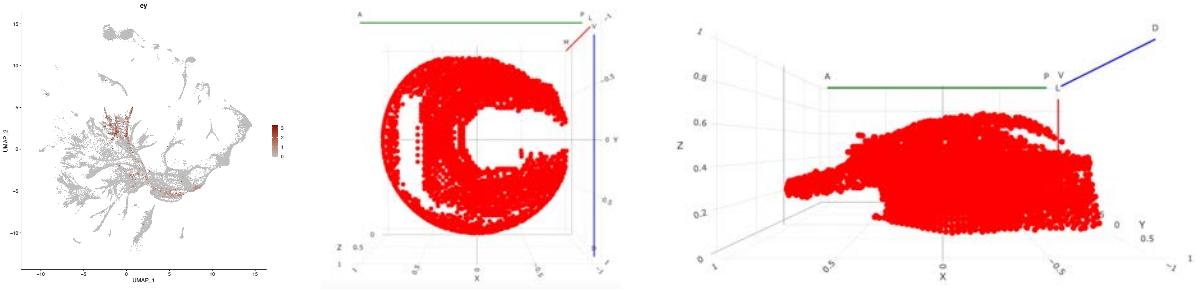


# fd59A

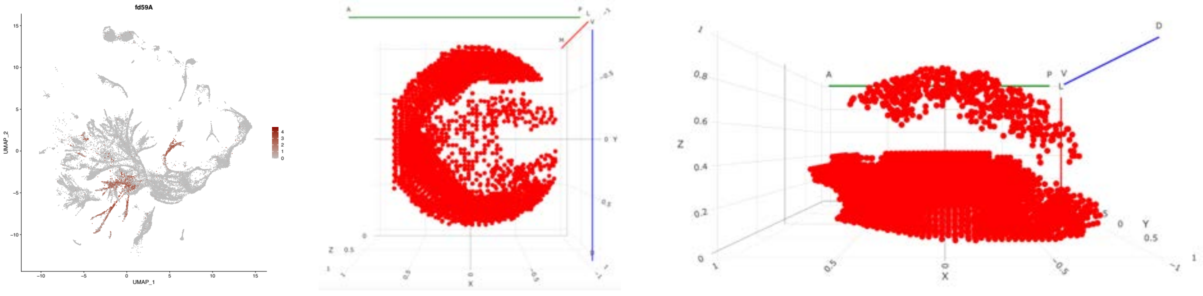

### foxo

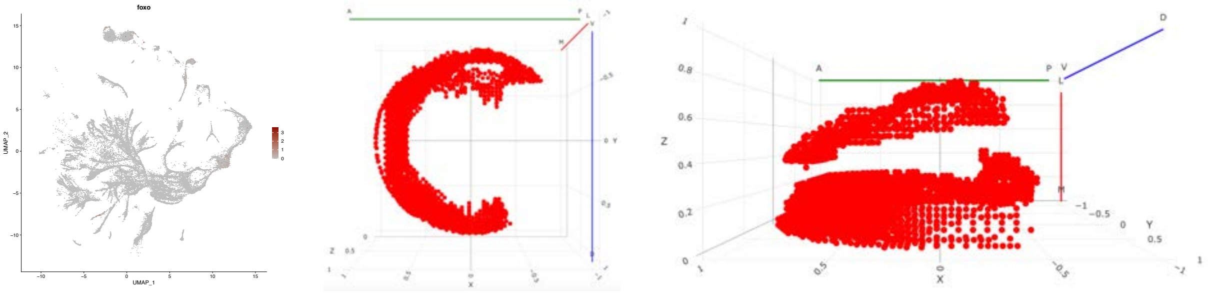


### ham

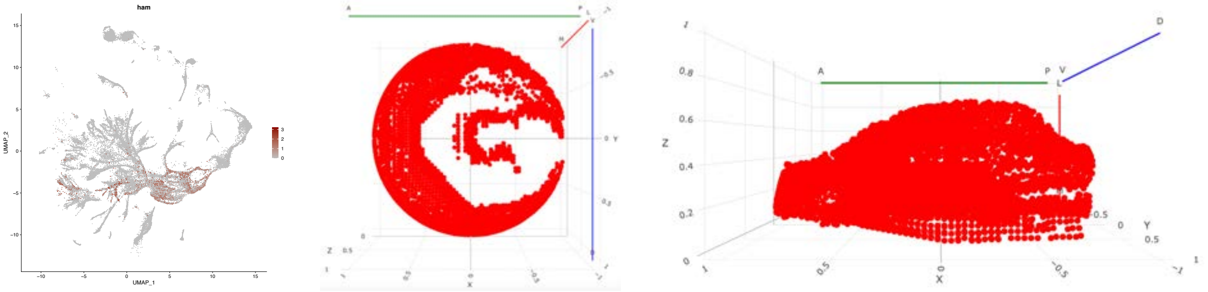


### hbn

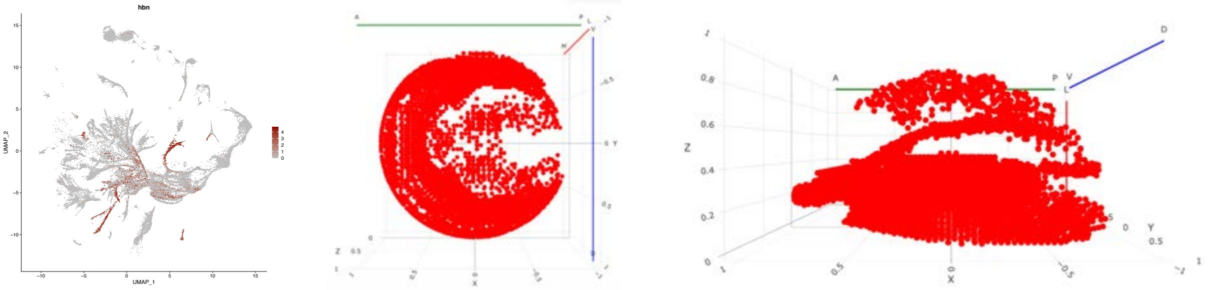


# kn

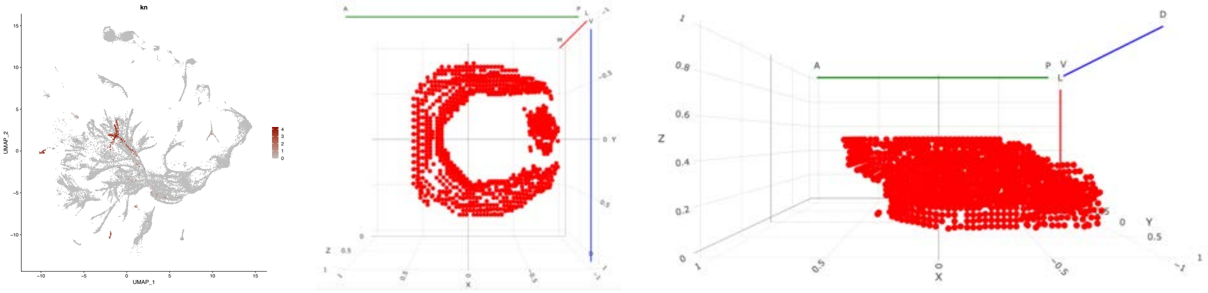


### Lim3

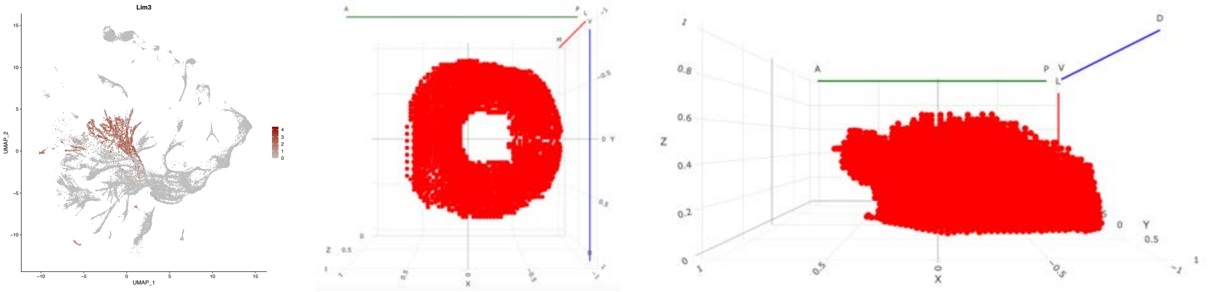

## NK7.1

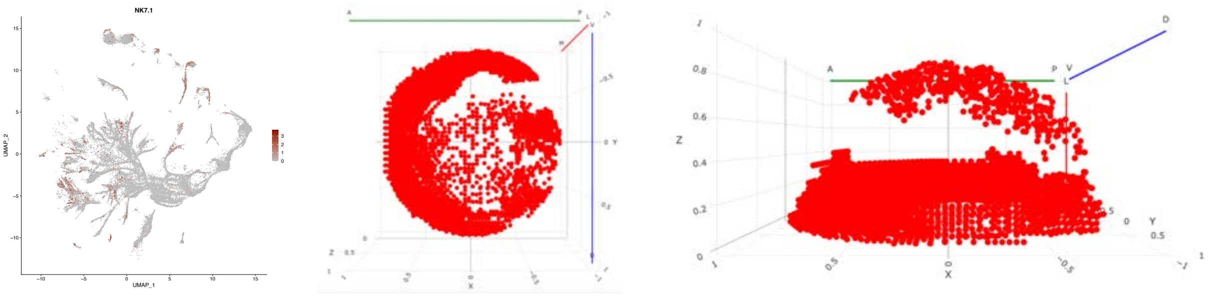


#### run

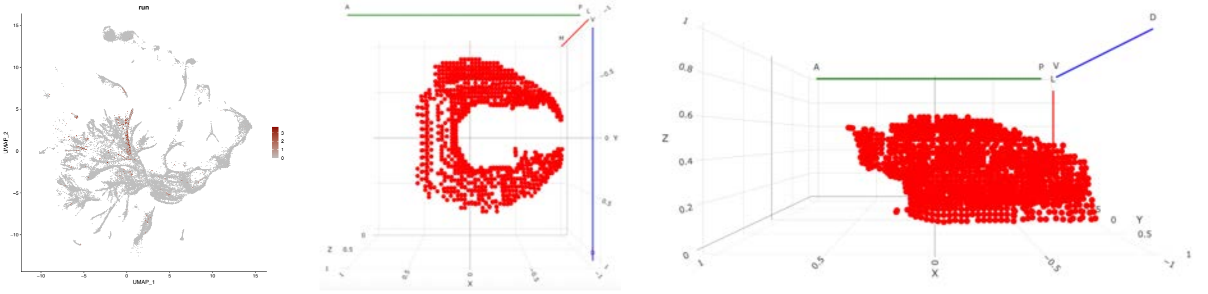


#### sim

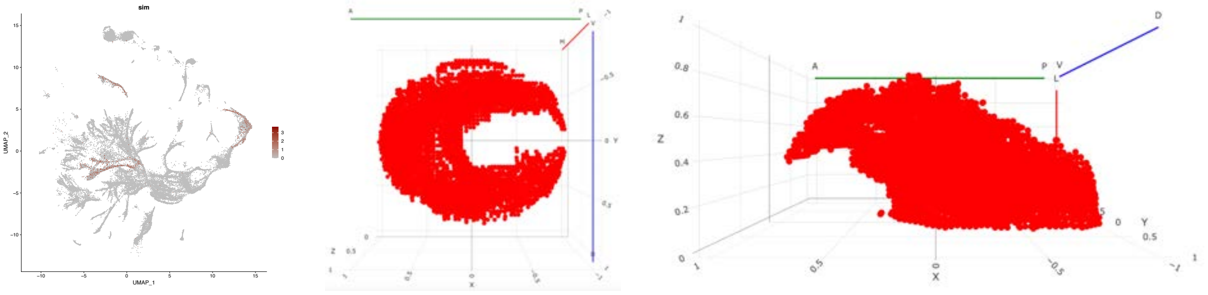


#### Sox102F

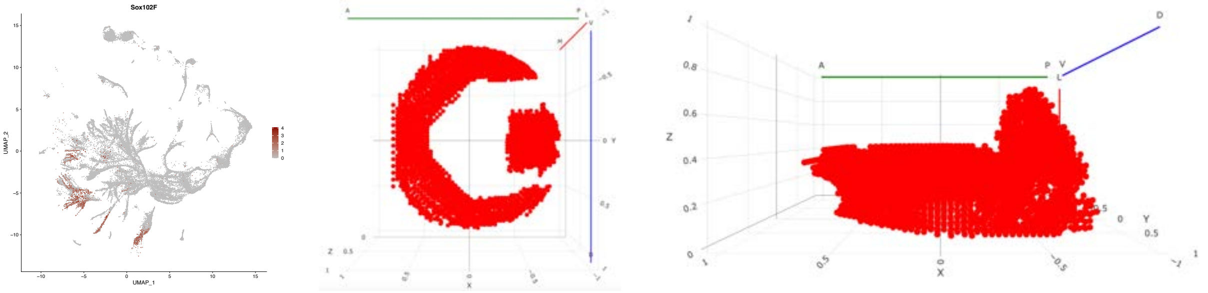


#### TfAP-2

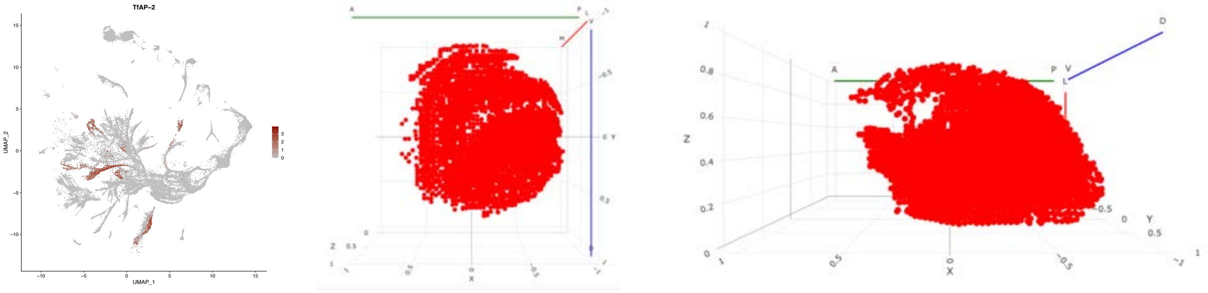

### toy

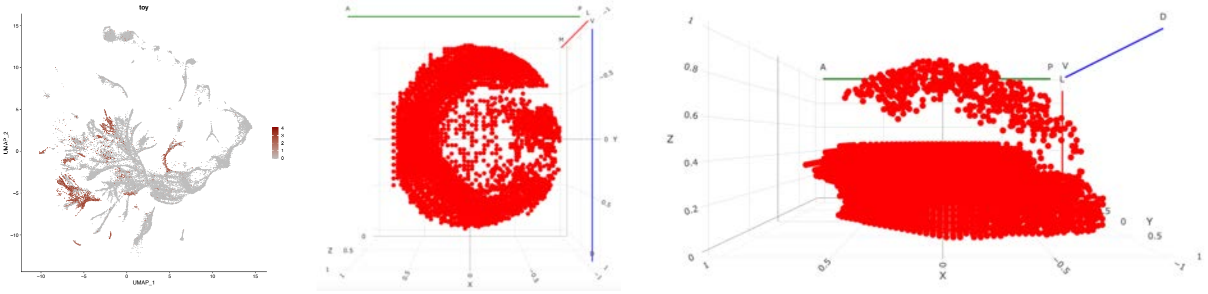


### tsh

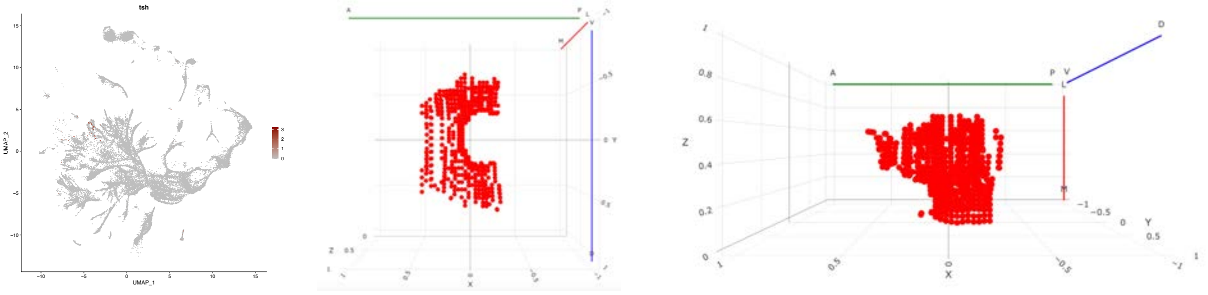


### tup

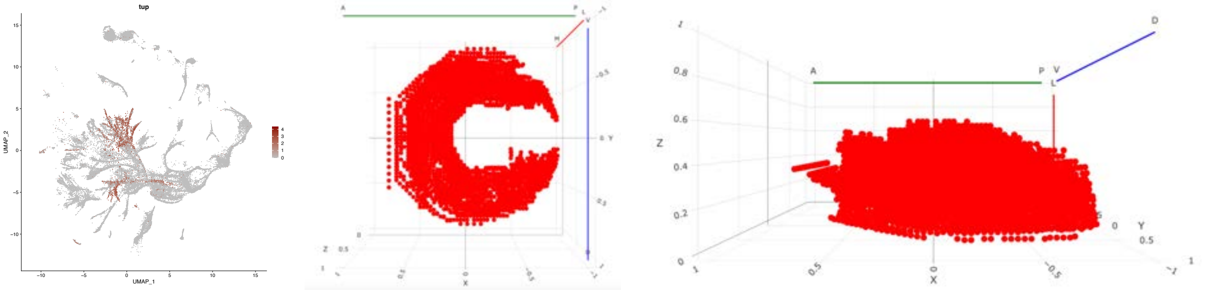


### vvl

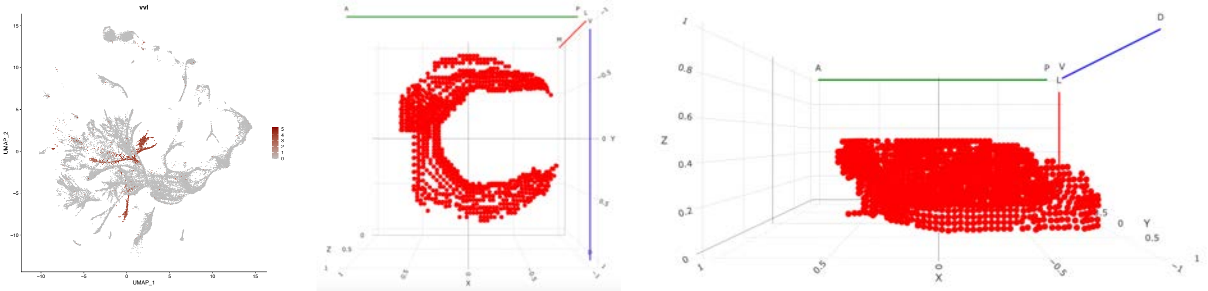


### zfh1

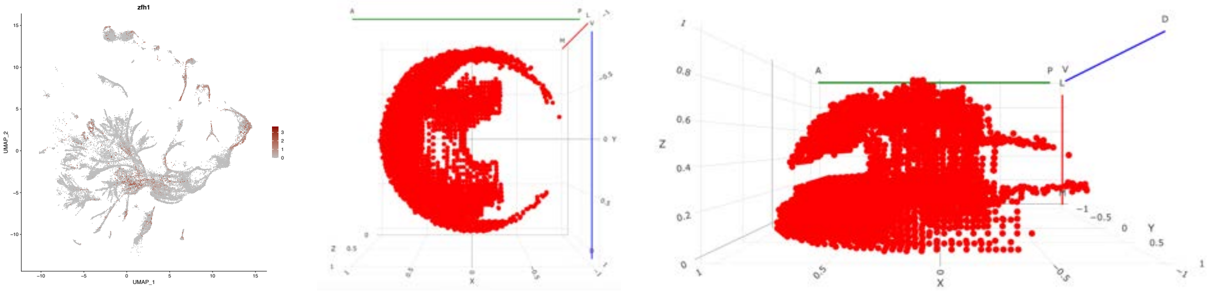

### zfh2

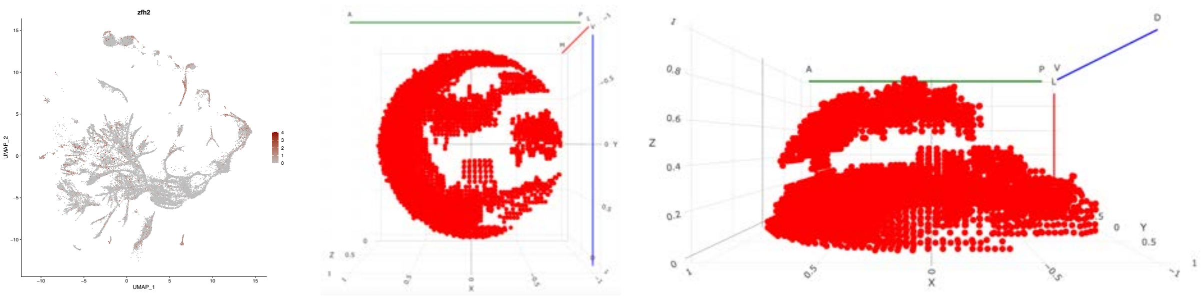

### spatiotemporal regulation

#### Awh

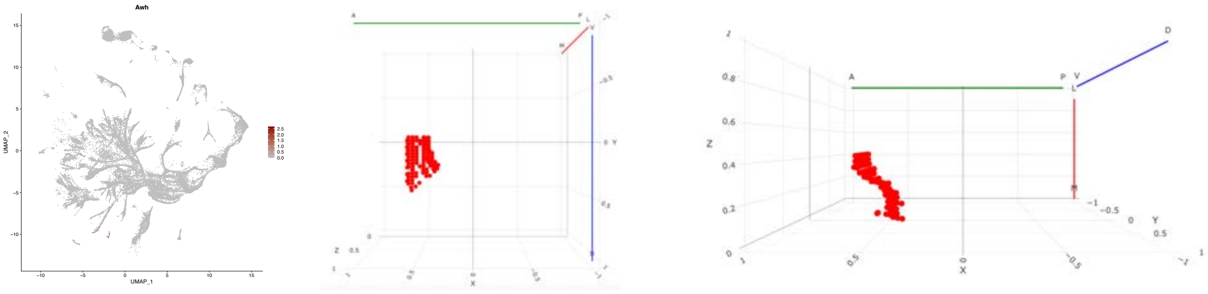

#### bab2

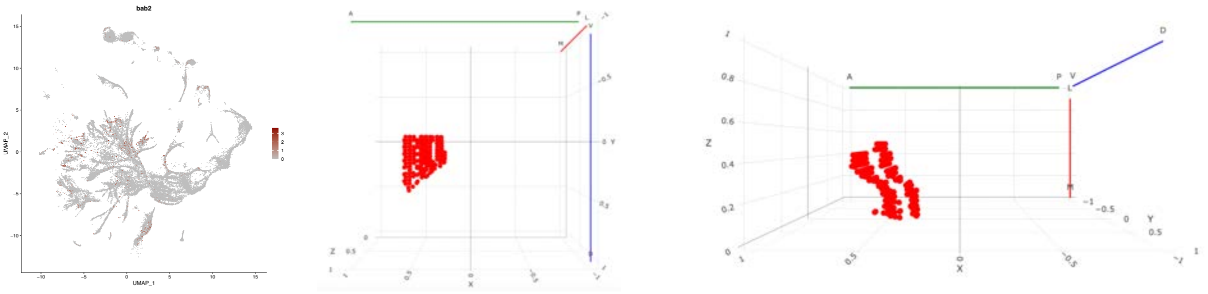

## CG3726

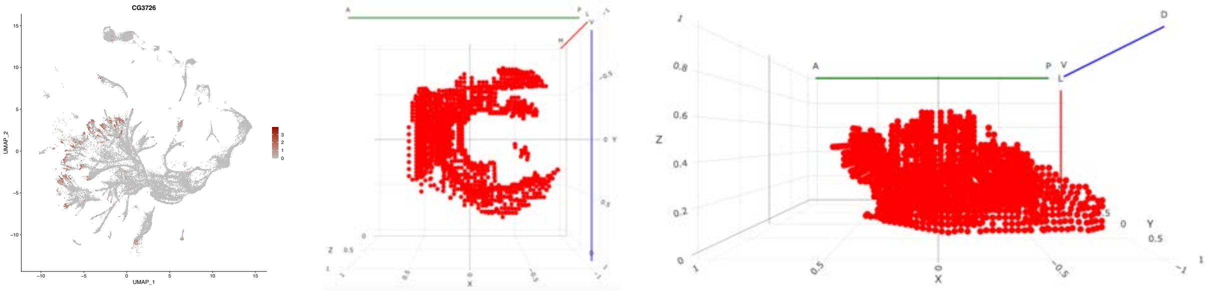

## CG32532

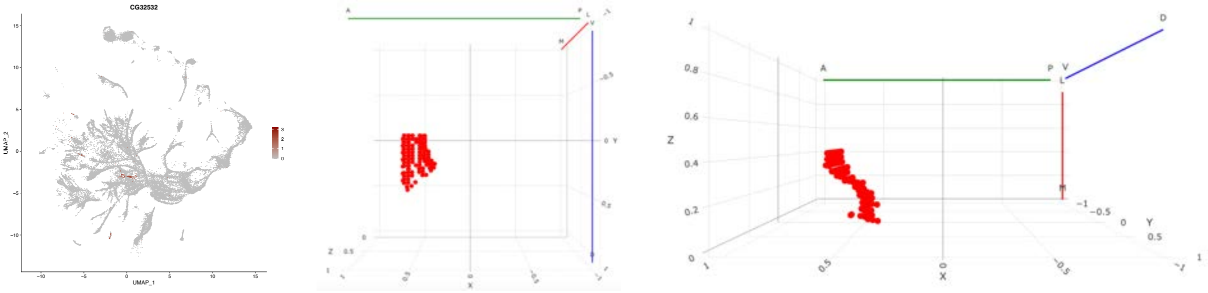

#### Dll

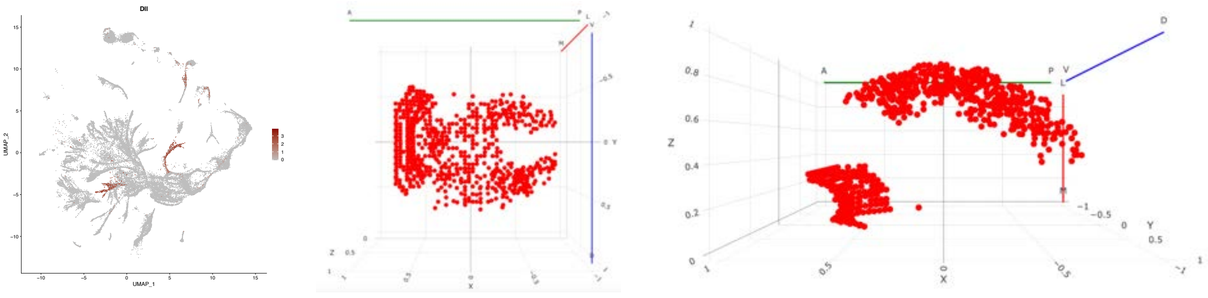

### disco

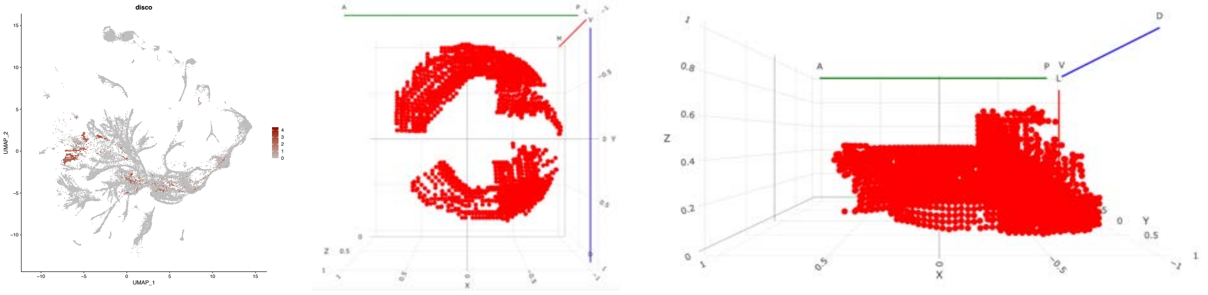

### disco-r

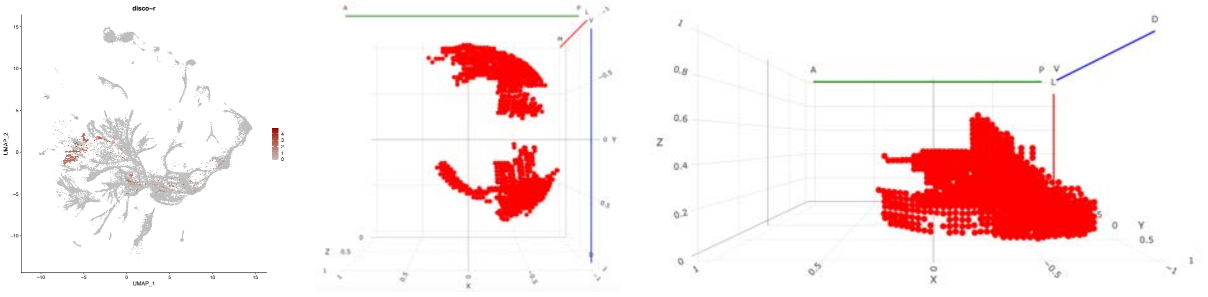

### fkh

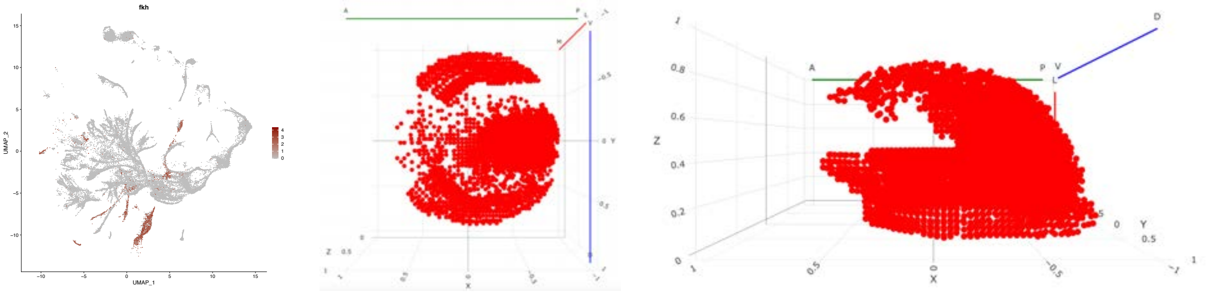

### Hmx

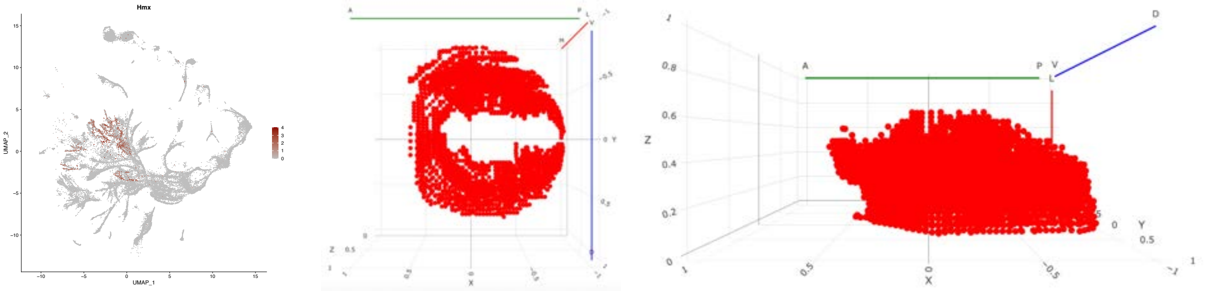

### lov

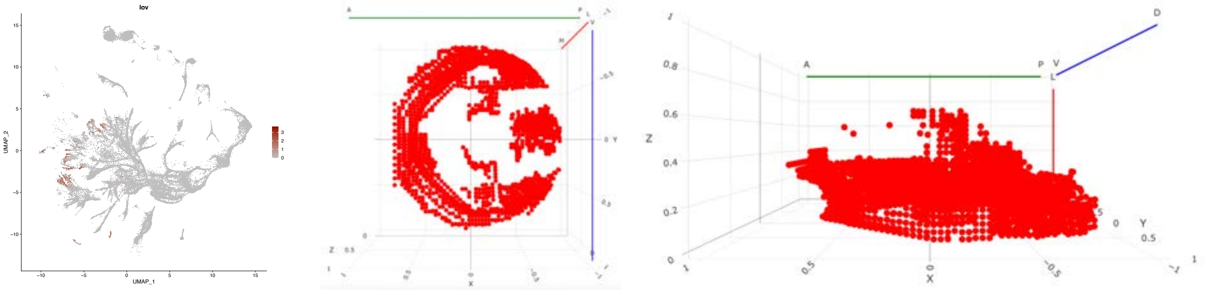

#### mirr

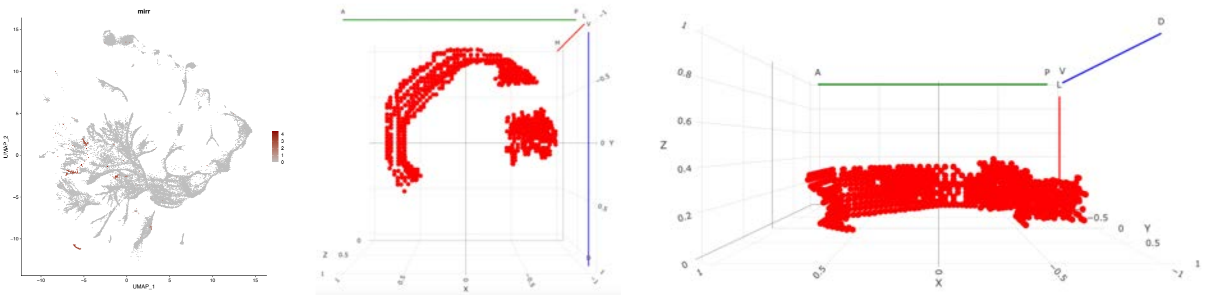

#### pdm3

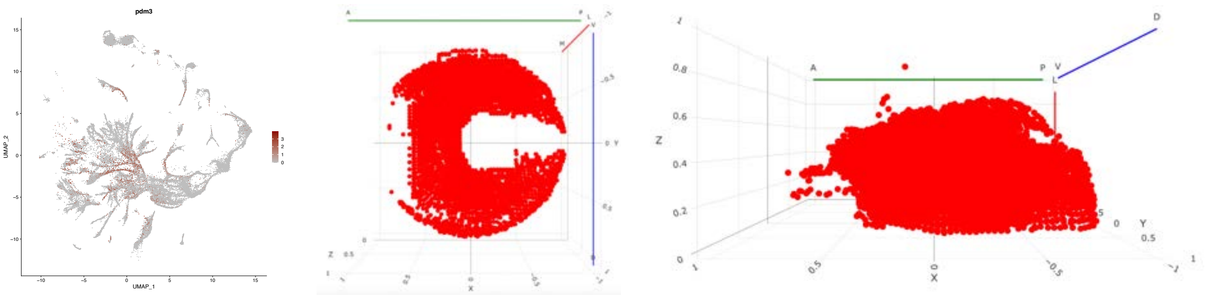

## rn

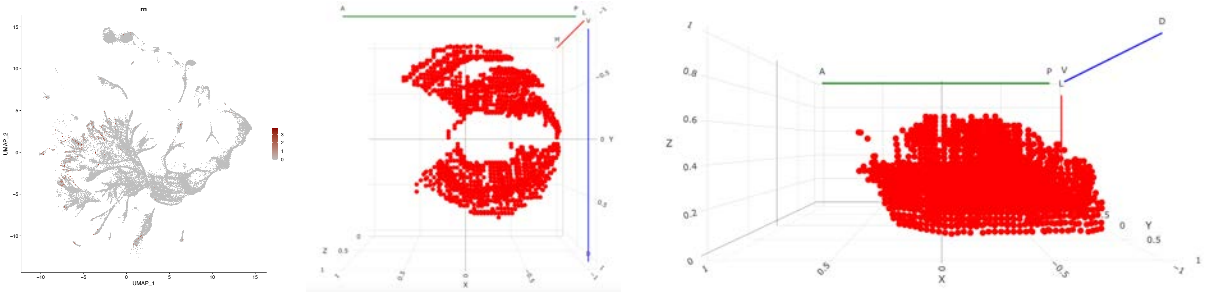

#### RunxA

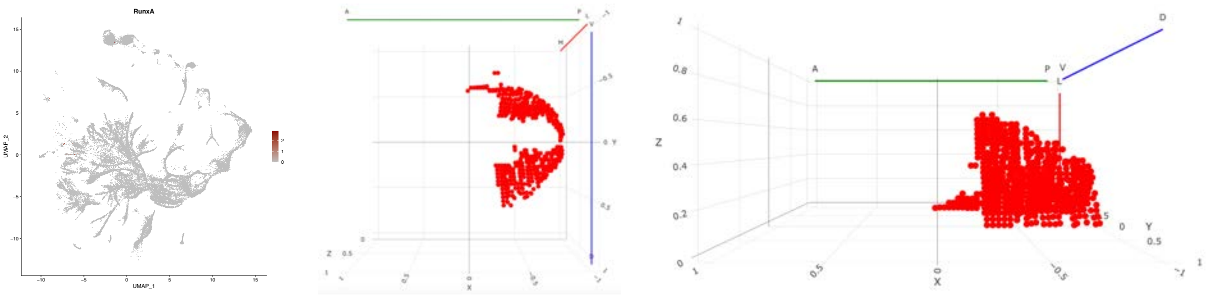

#### salm

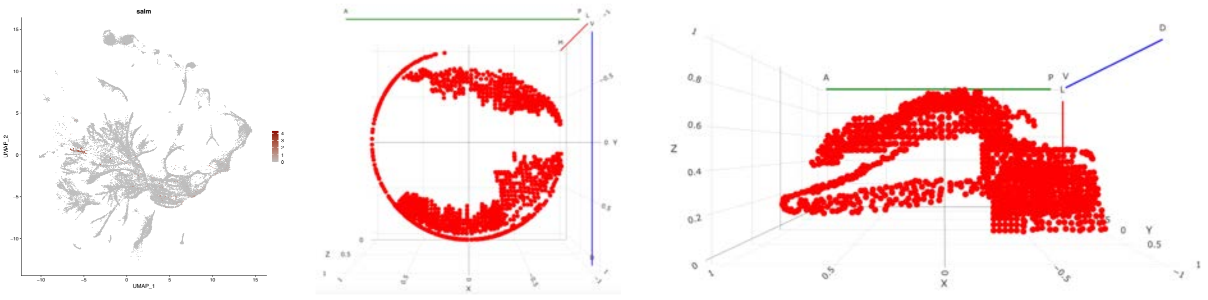

### salr

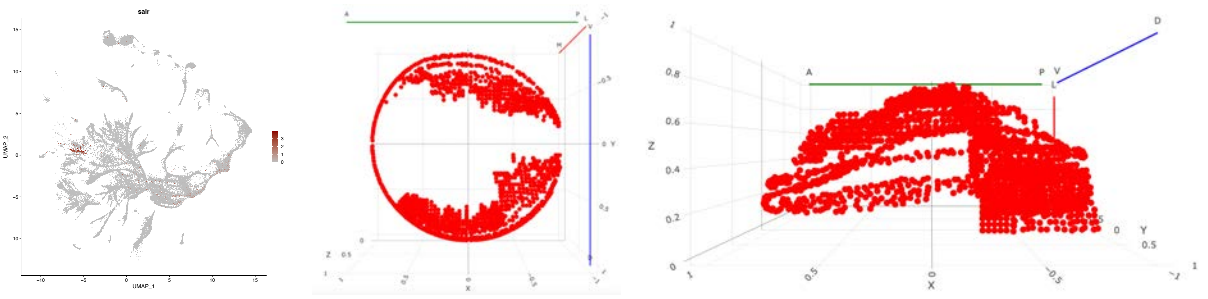
salr
UMAP_1
UMAP_2
A
P
V
L
D
X
Y
Z


# tj

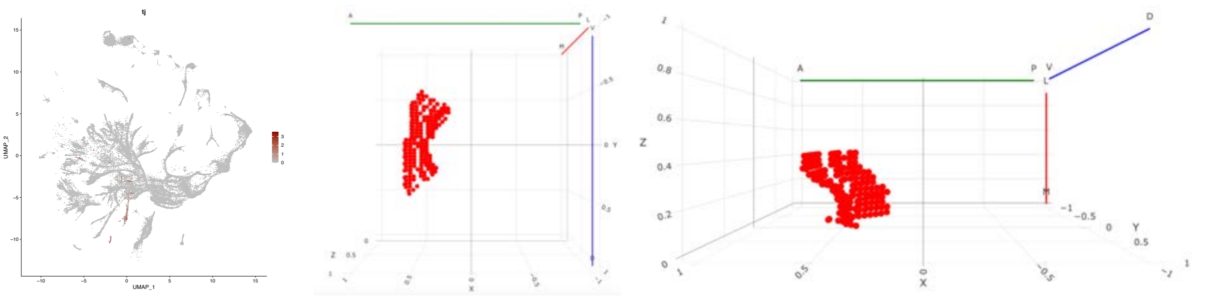
tj
UMAP_1
UMAP_2
A
P
V
L
D
X
Y
Z


### Vsx2

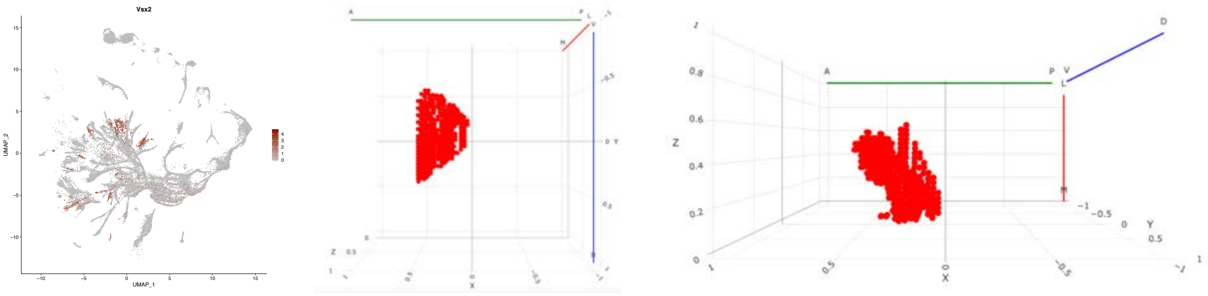
Vsx2
UMAP_1
UMAP_2
A
P
V
L
D
X
Y
Z

### complex regulation

#### bab1

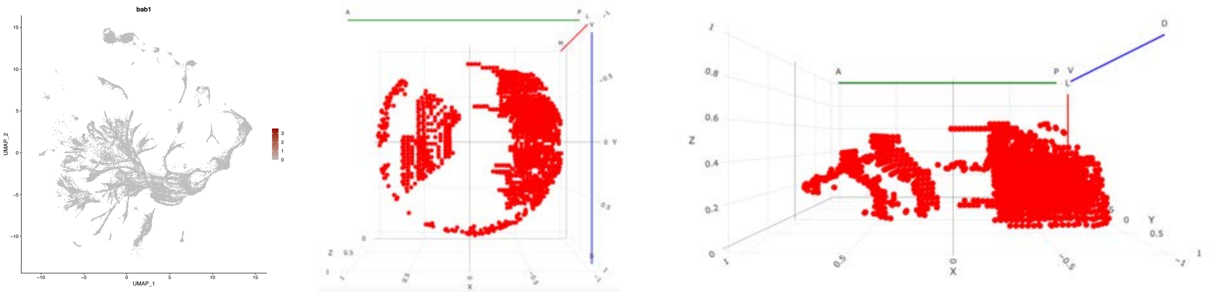

## bi

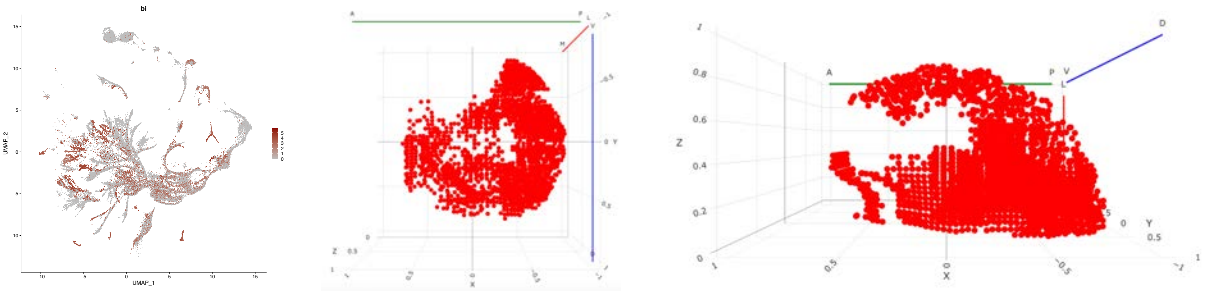

## CG9932

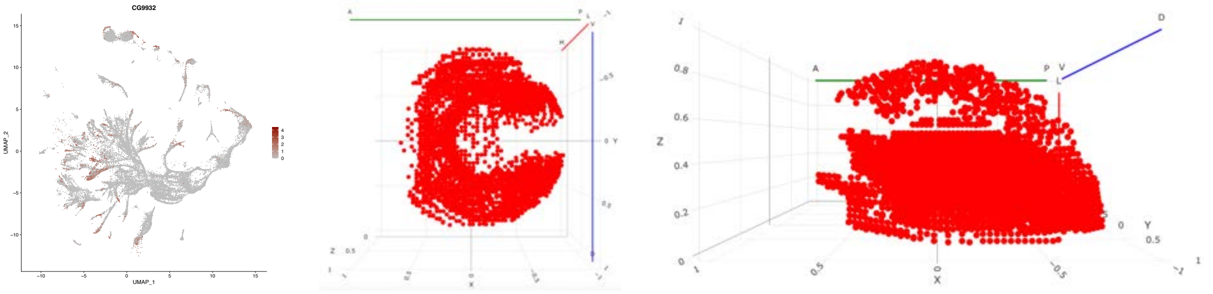

## D

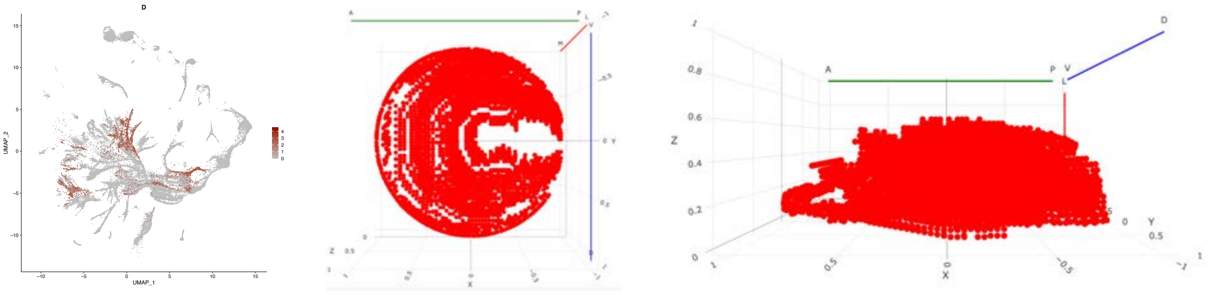

#### hth

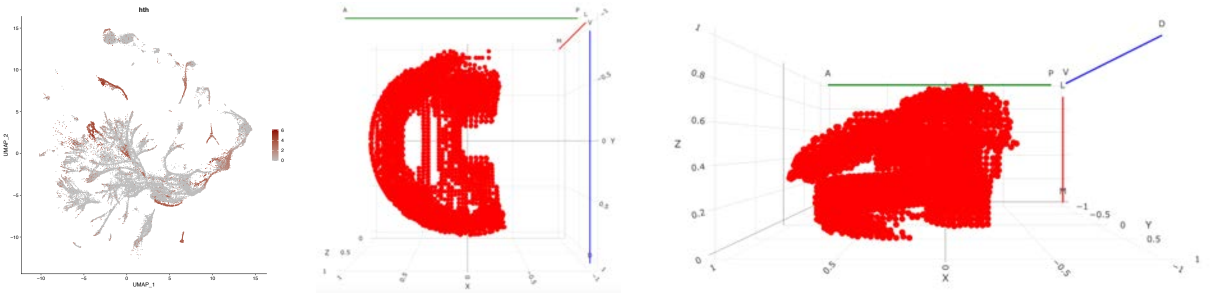

#### Sox21B

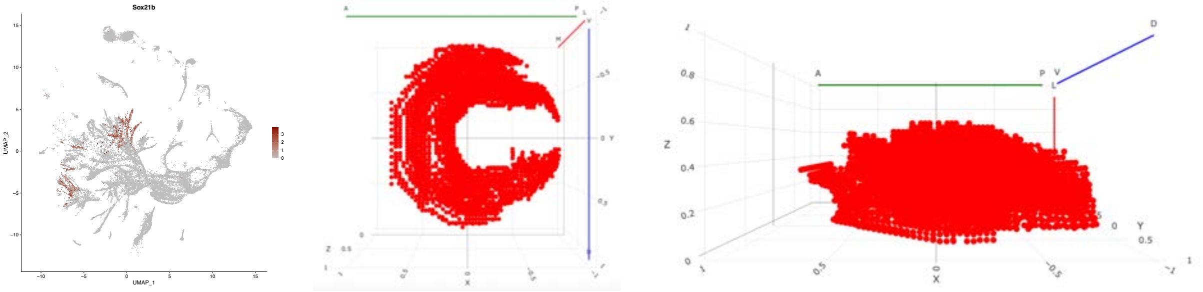


#### Stat92E

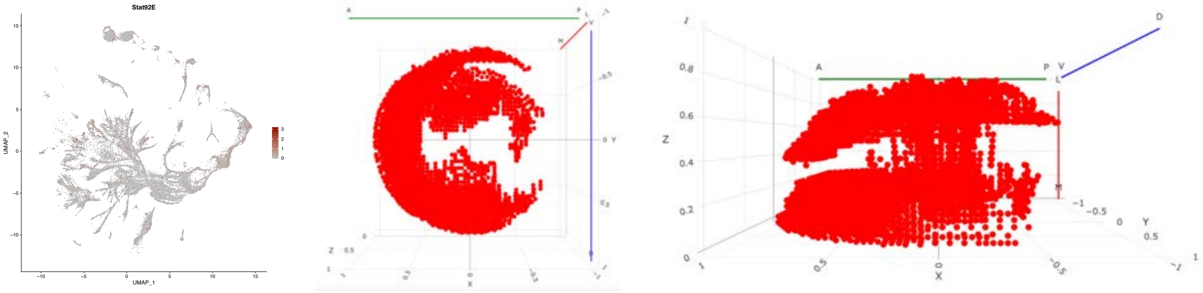


#### svp

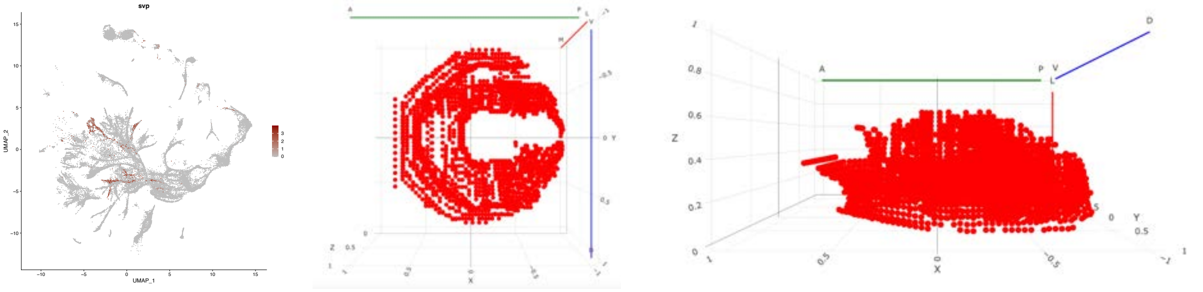


#### Vsx1

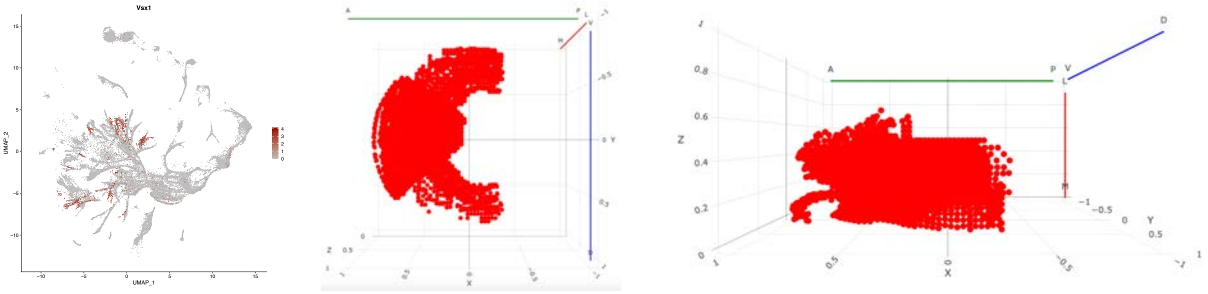

### neuropil-specific expression

#### acj6

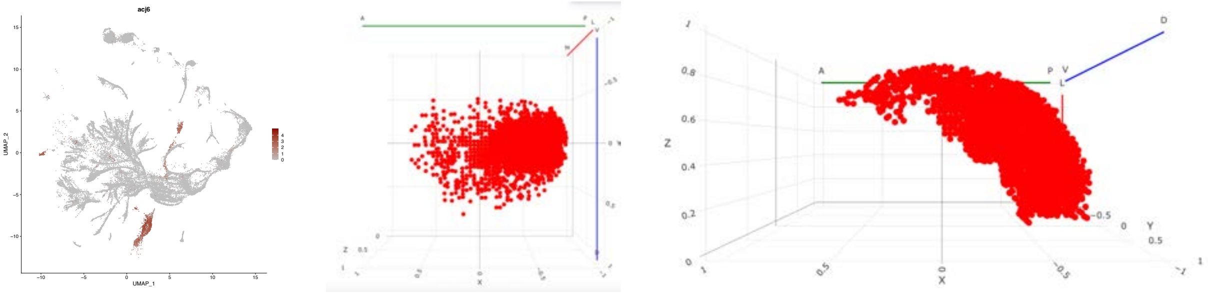


#### ato

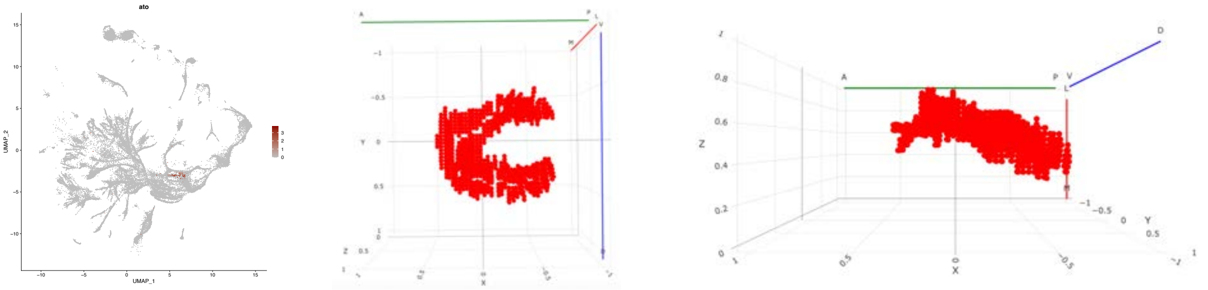


## CG34340

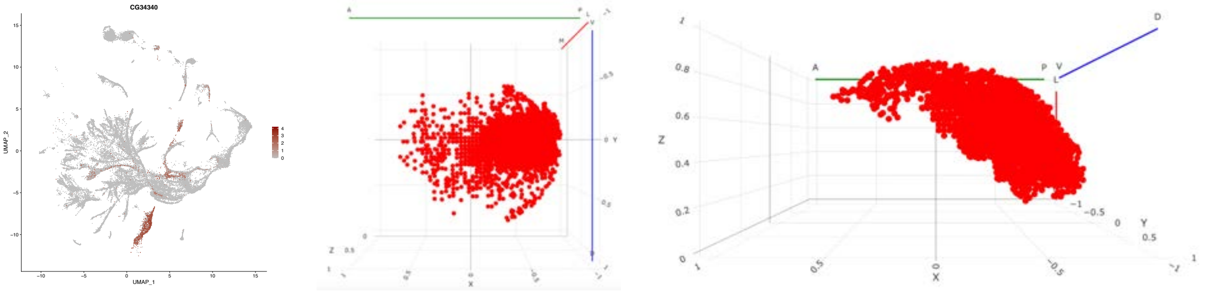


#### dac

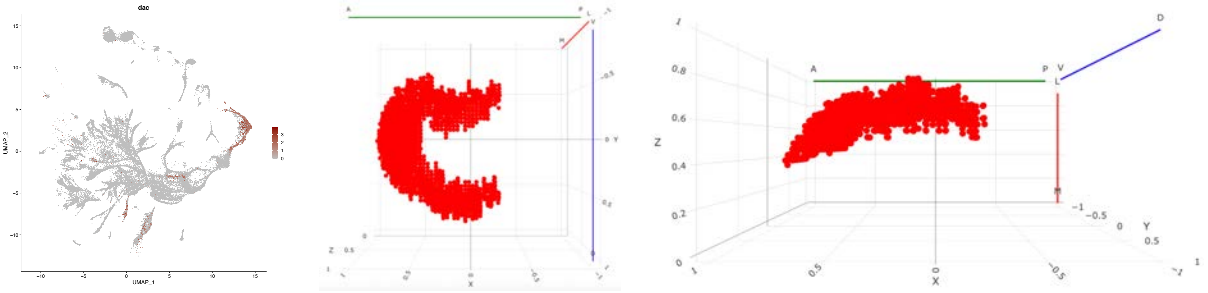


#### dan

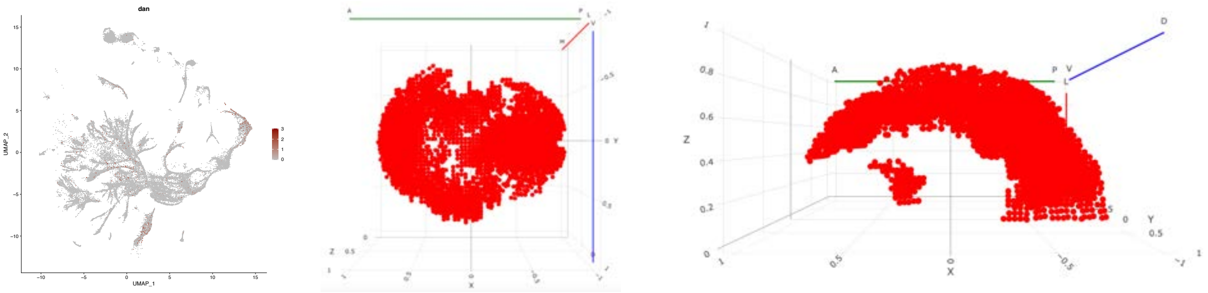

### danr

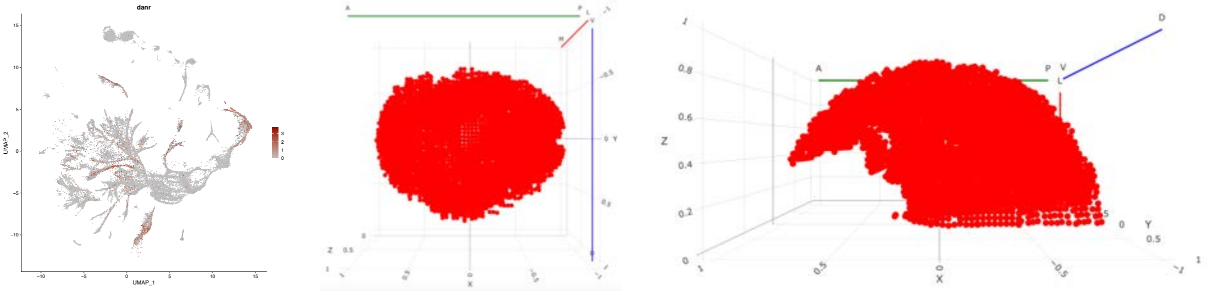

### fru

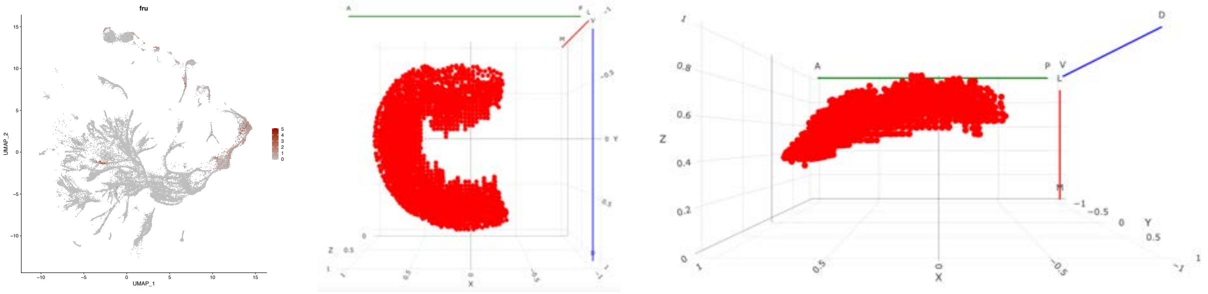

### grn

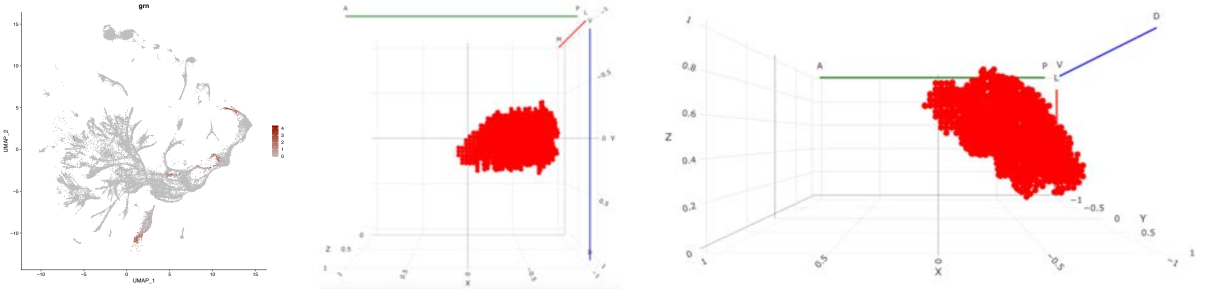

### RunxB

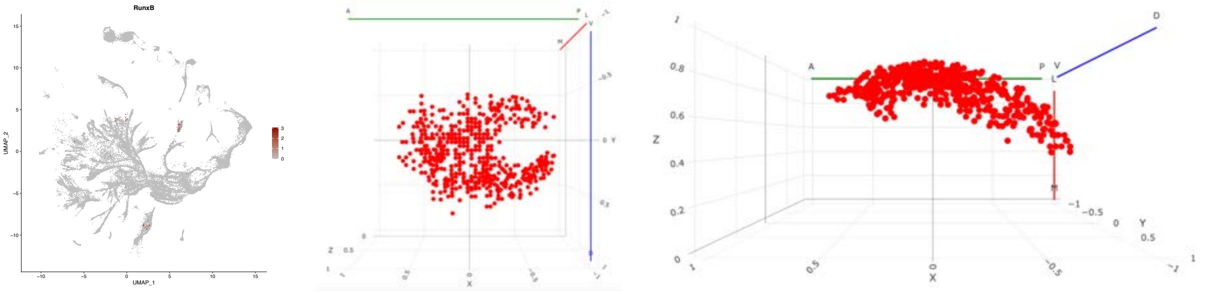

### restricted expression

#### ara

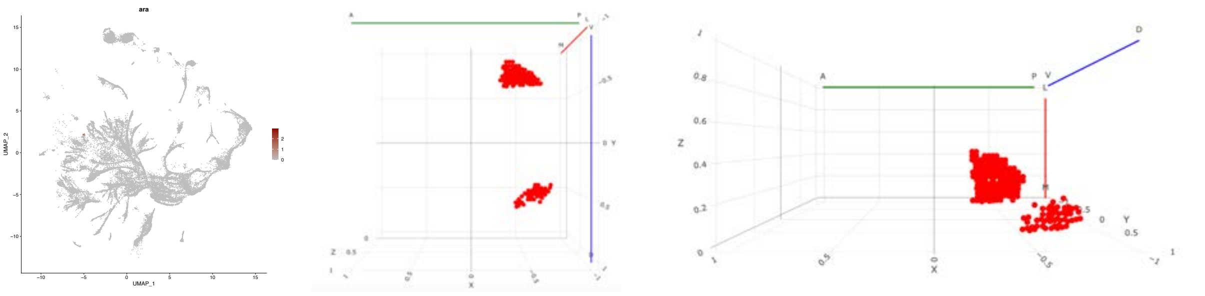

#### Camta

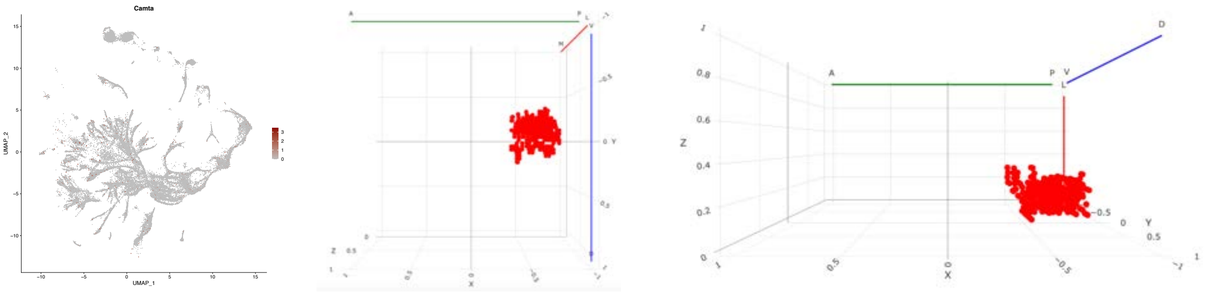

#### caup

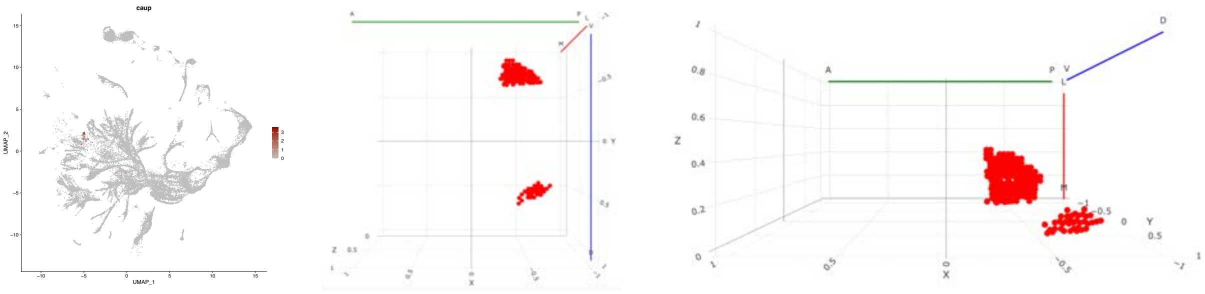

## CG4328

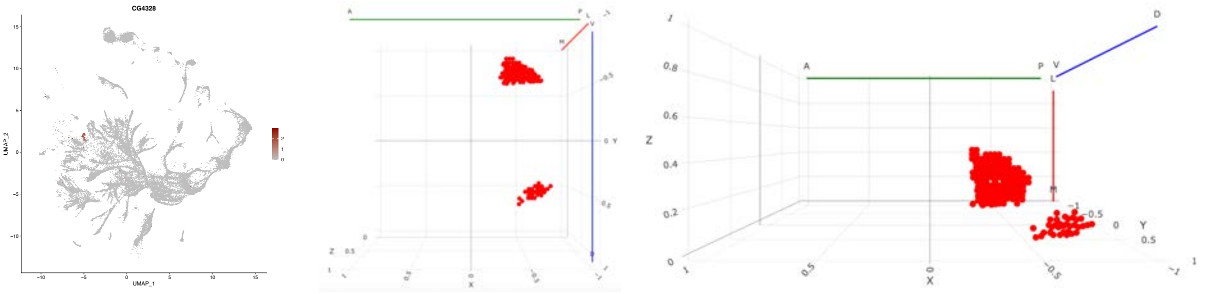

## CG32105

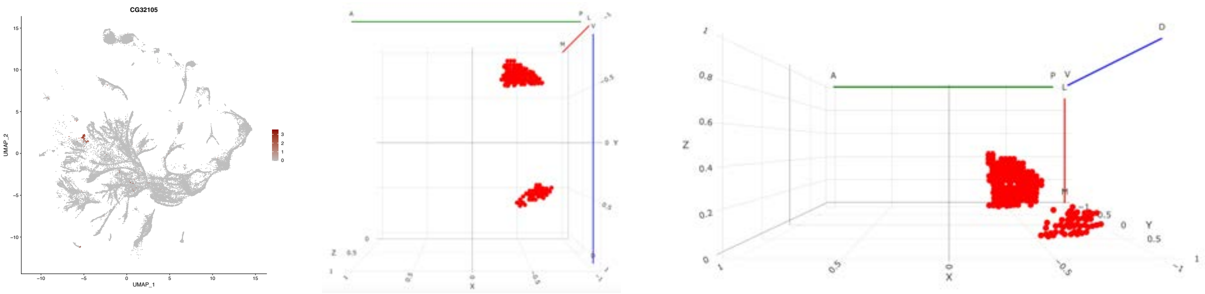

### dmrt99B

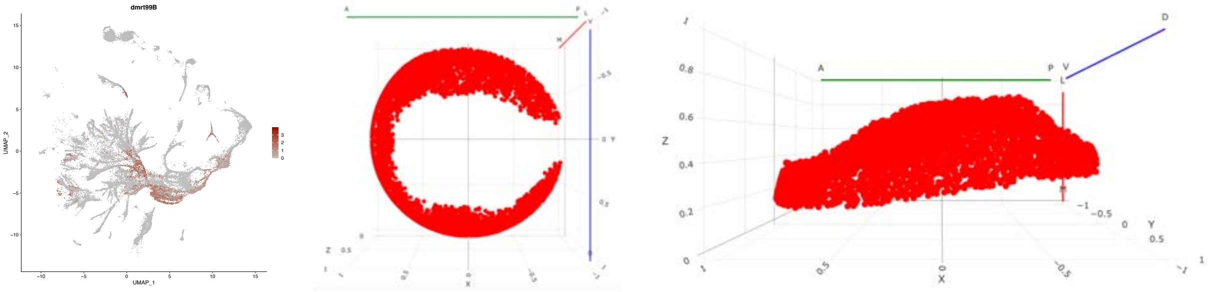


### Fer2

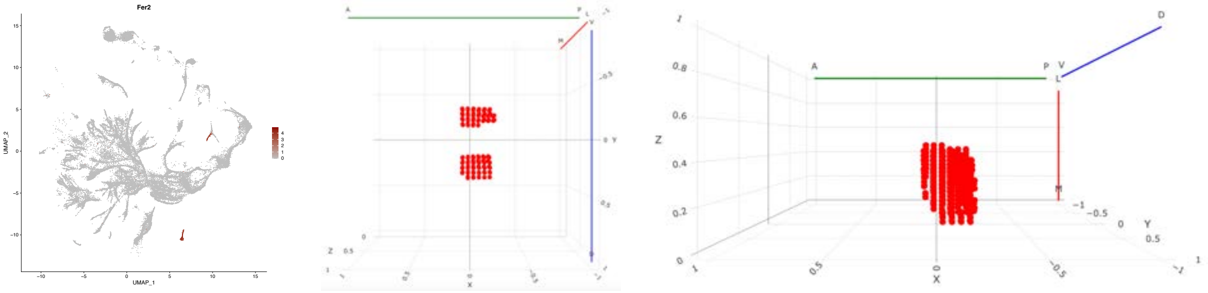


### klu

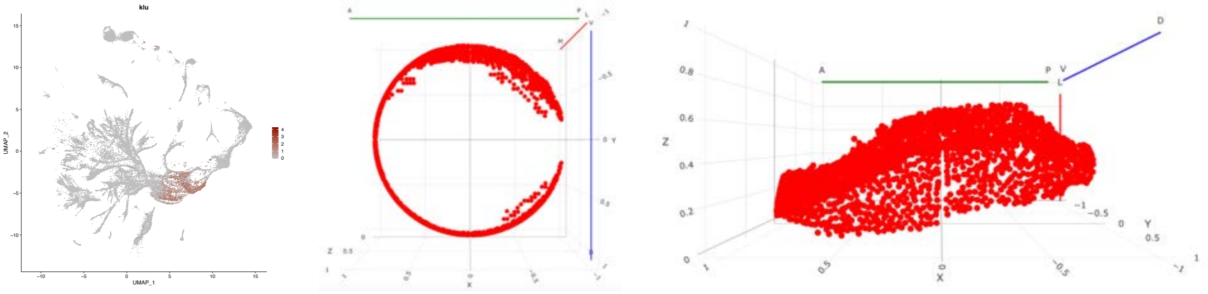


### Lim1

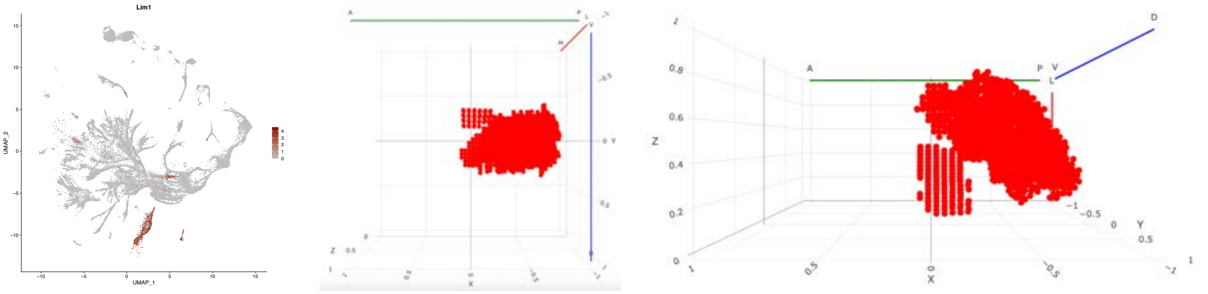


### Lin29

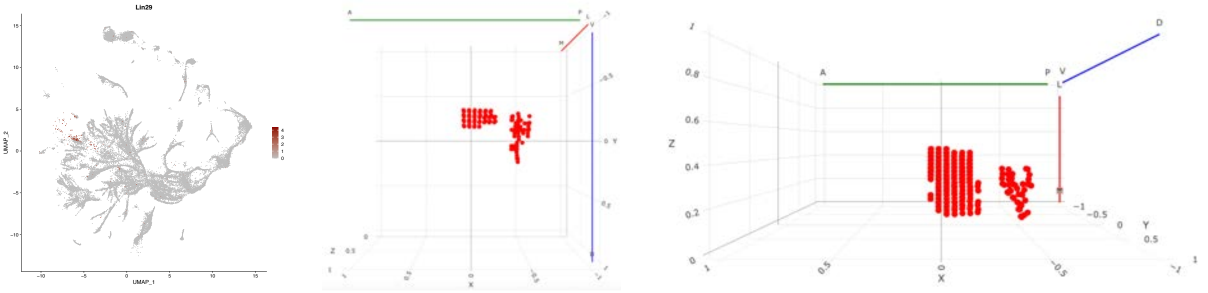

## oc

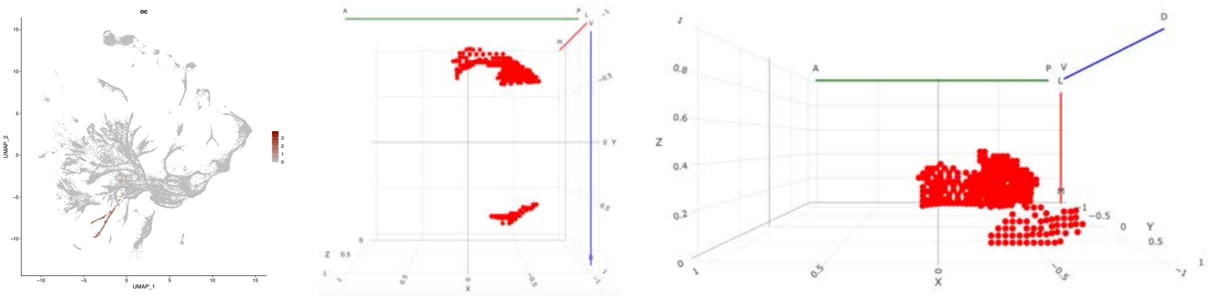

#### opa

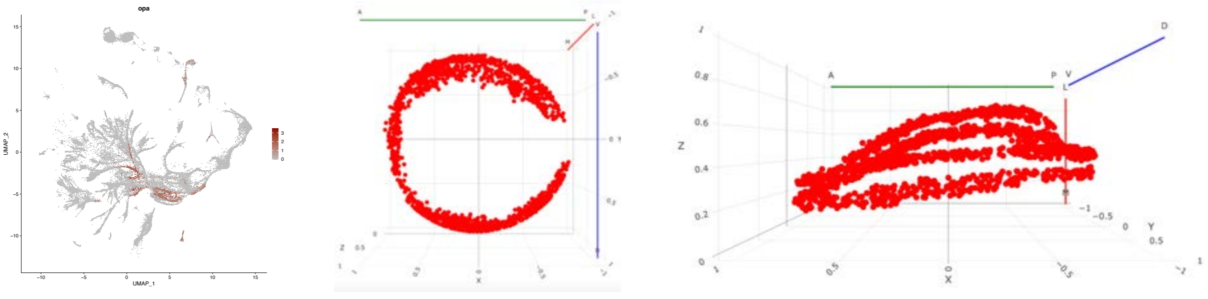

## Rx

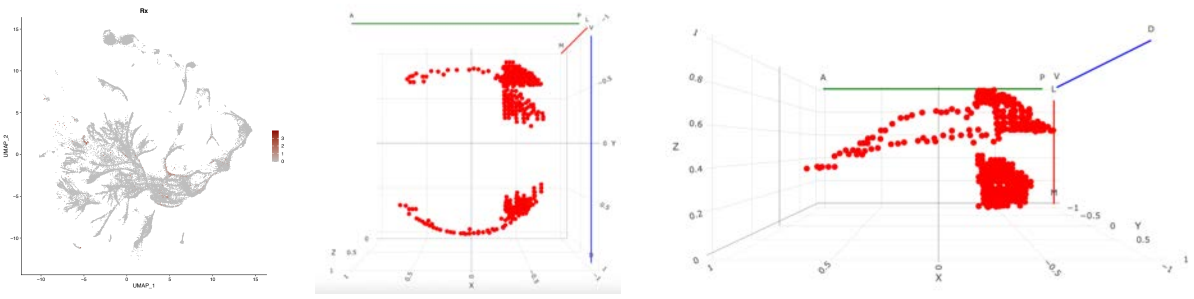

#### slou

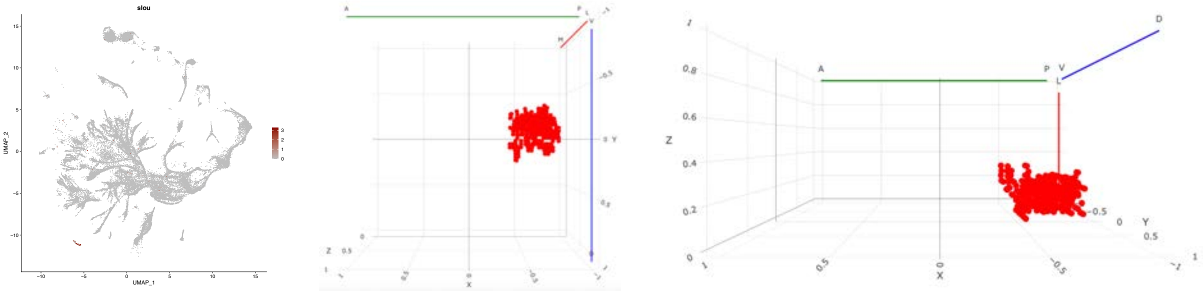

#### slp1

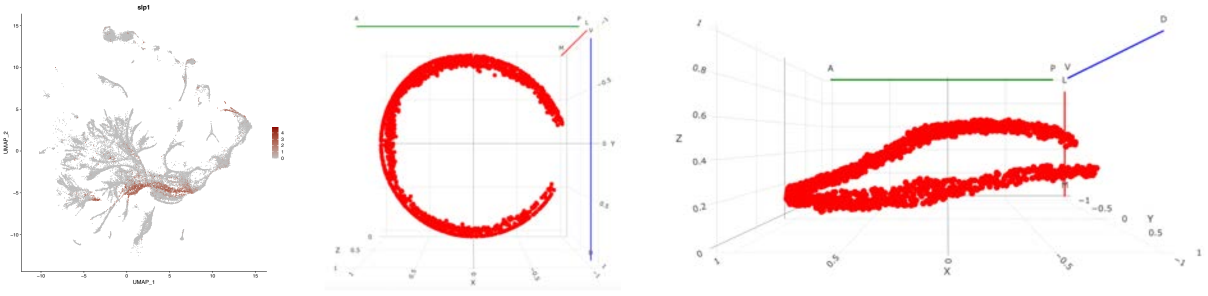

### slp2

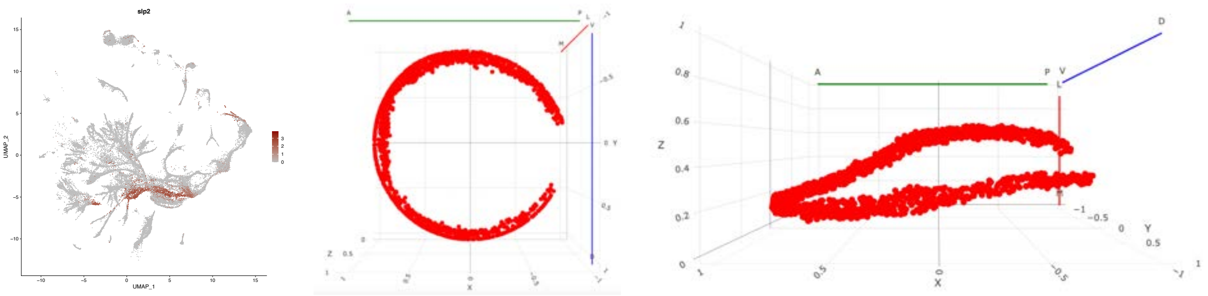

#### broad expression

### br

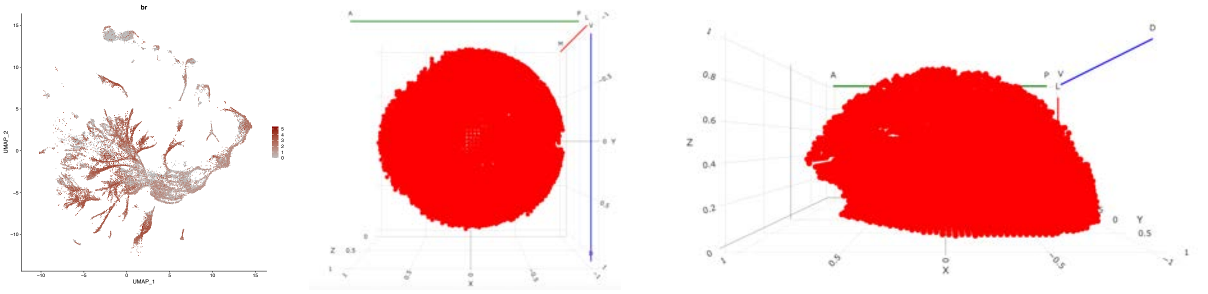

### CG9650

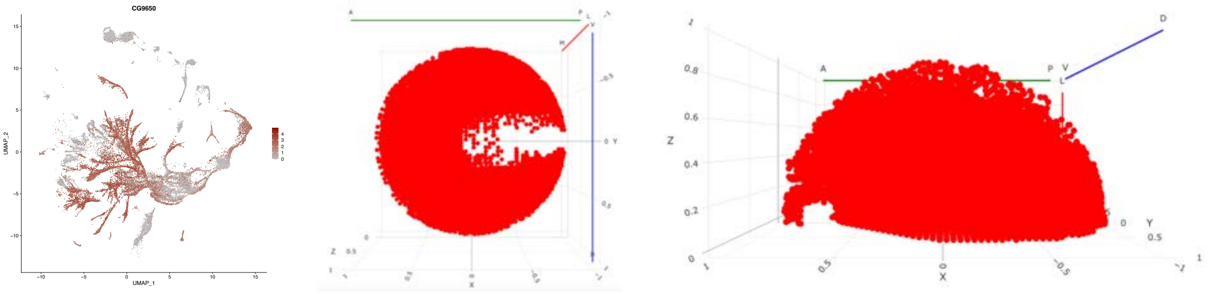

### ct

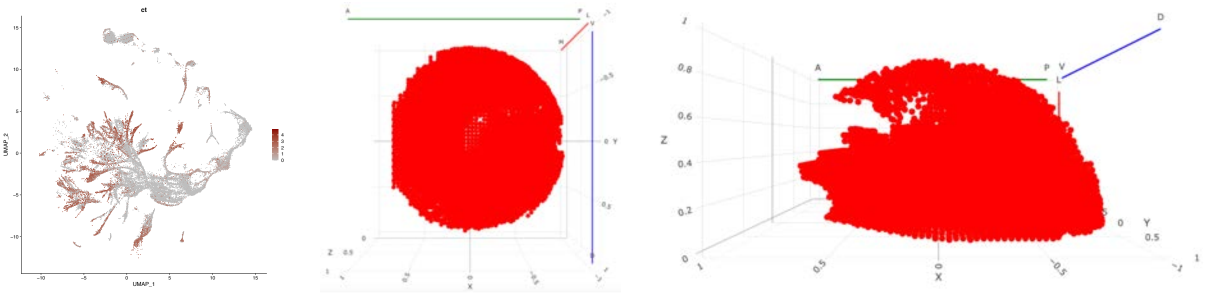

##### erm

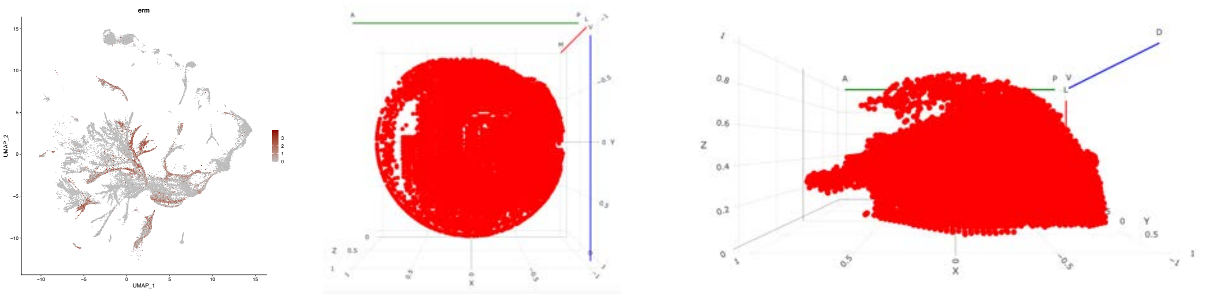

##### luna

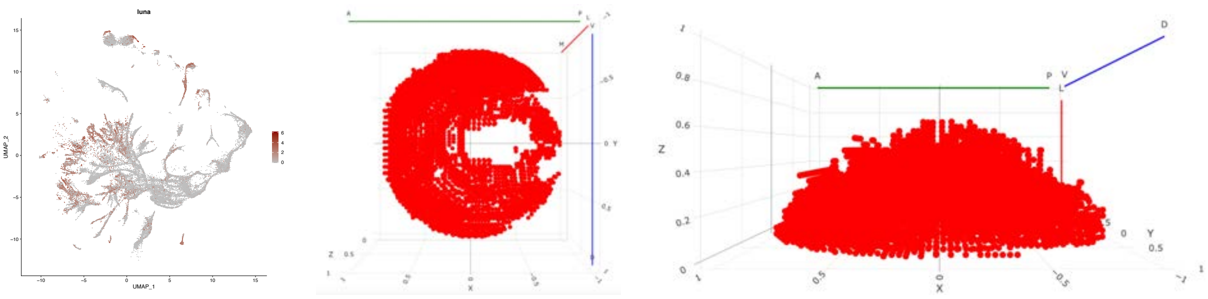

### noc

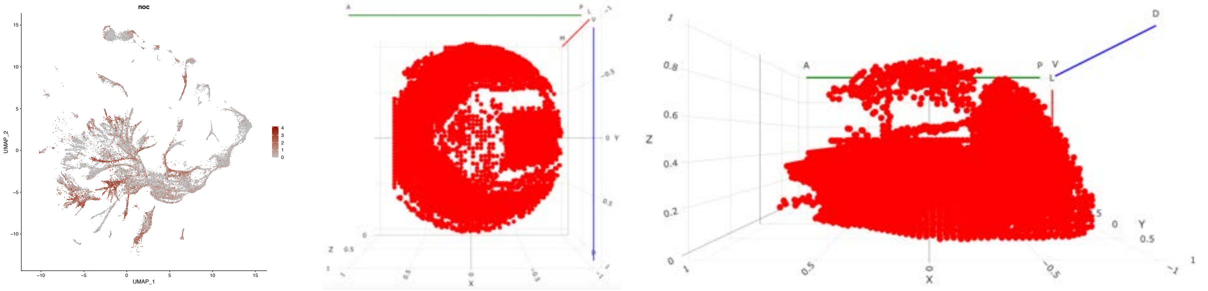

### pros

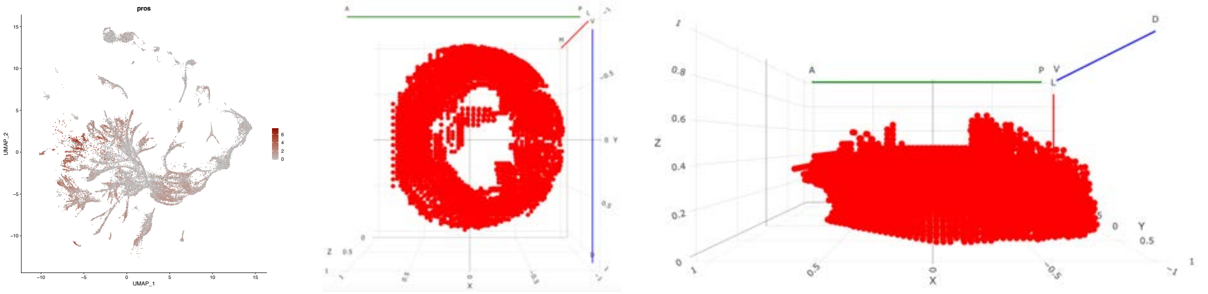

### scro

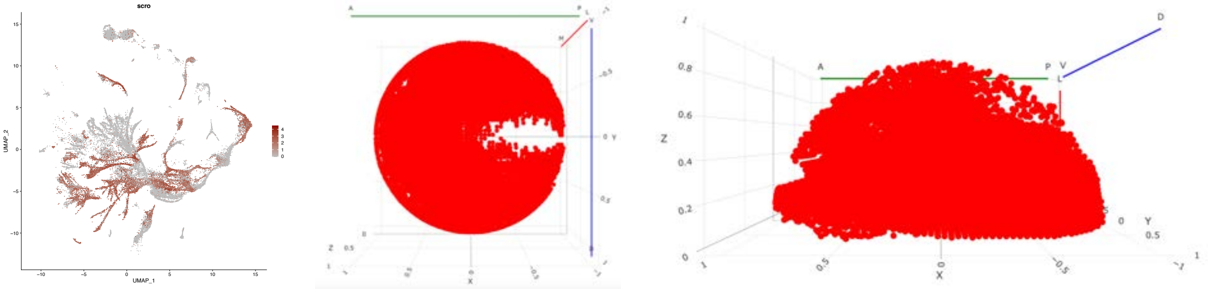

### Snoo

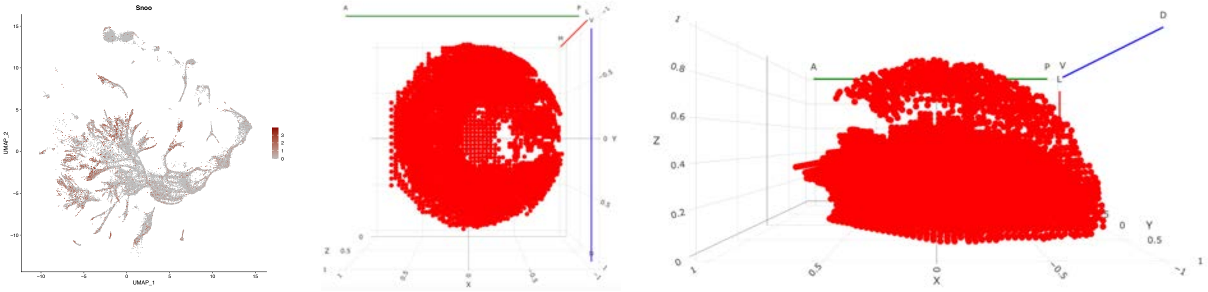

### SoxN

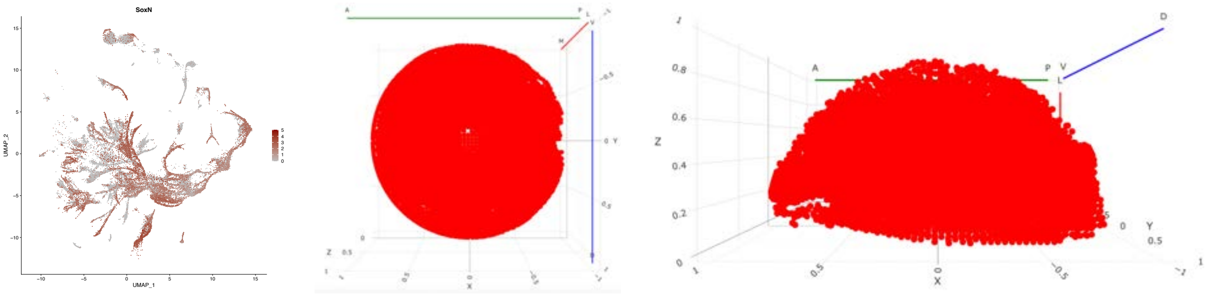

# sr

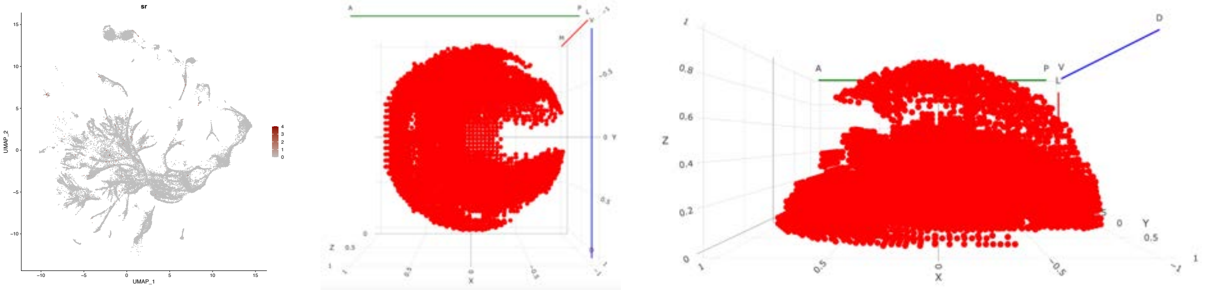


### vfl

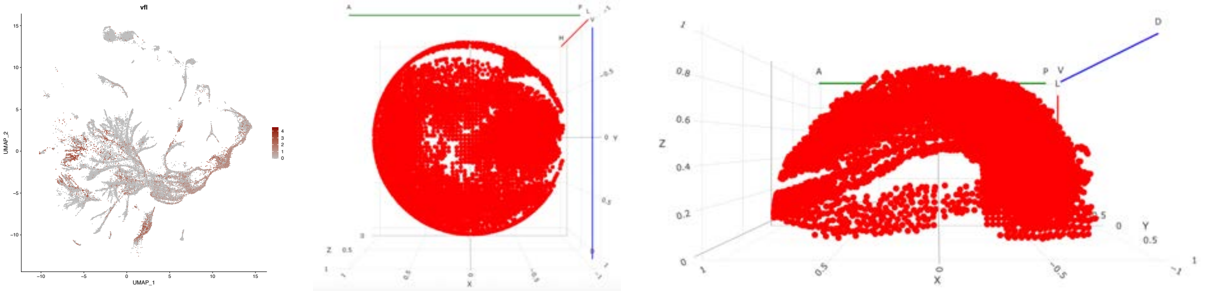

### Figure S6: Webpage description

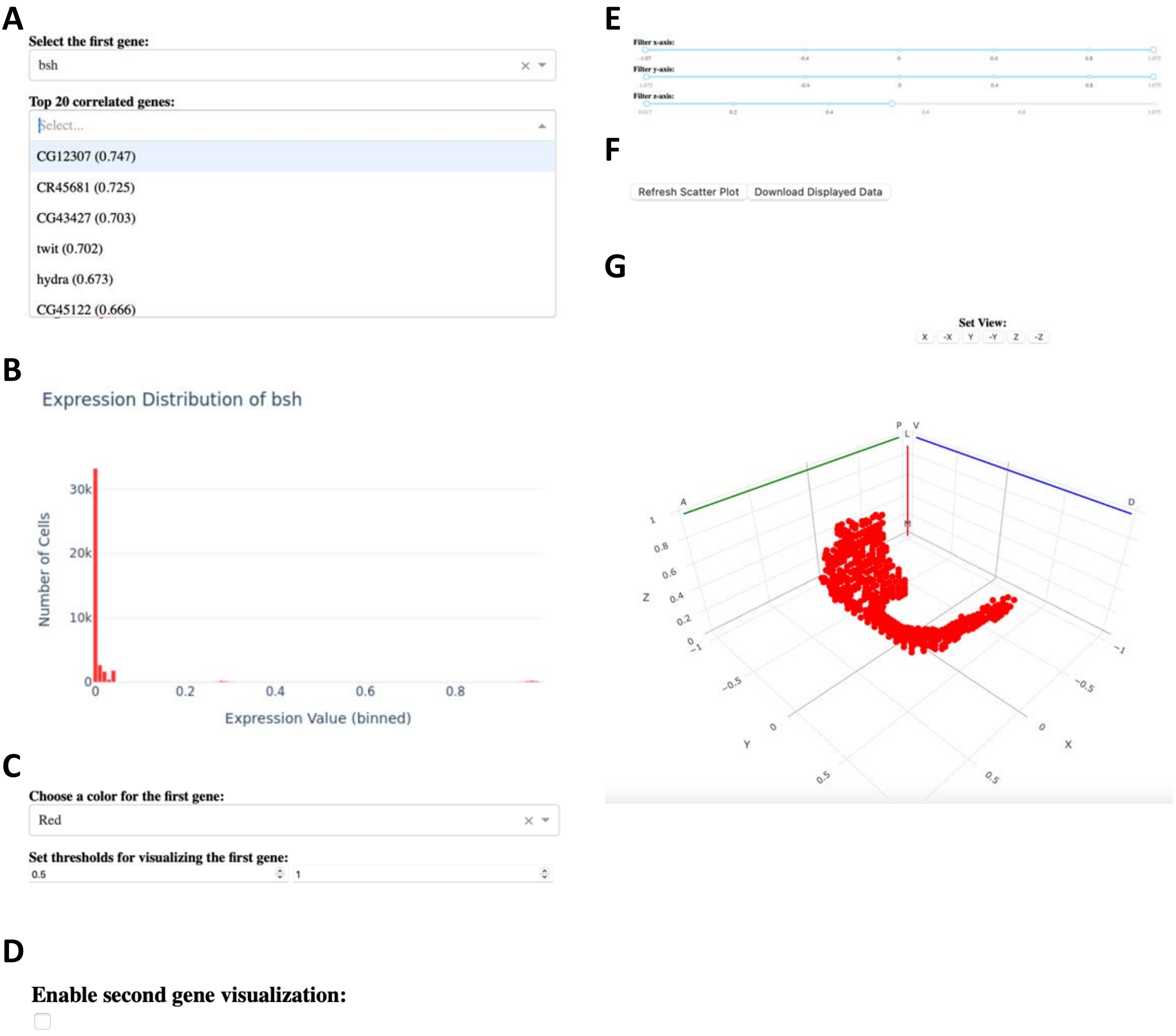
